# Non-cancer stem cell-derived Fibromodulin activates Integrin-dependent Notch signaling in endothelial cells to promote tumor angiogenesis and growth

**DOI:** 10.1101/2022.04.03.486893

**Authors:** Shreoshi Sengupta, Mainak Mondal, Kaval Reddy Prasasvi, Arani Mukherjee, Prerna Magod, Serge Urbach, Dinorah Friedmann-Morvinski, Philippe Marin, Kumaravel Somasundaram

**Affiliations:** Department of Microbiology and Cell Biology, Indian Institute of Science, Bangalore 560012, India; School of Neurobiology, Biochemistry and Biophysics, The George S. Wise Faculty of Life Sciences, Tel Aviv University, Tel Aviv 69978, Israel; Sagol School of Neuroscience, Tel Aviv University, Tel Aviv 69978, Israel; Institut de Génomique Fonctionnelle, Université de Montpellier, CNRS, INSERM, Montpellier, France

**Author notes:** Corresponding authors, Email: KS; PM; DFM. **Abbreviations**: FMOD, Fibromodulin; CSC, Cancer stem-like cell; GSC, Glioma stem-like cell; DGC, Differentiated glioma cell; GBM, Glioblastoma; CM, Conditioned medium; GSI, γ-secretase inhibitor, TDEC, Tumor derived endothelial cells; VM, Vascular mimicry.

**Keywords:** Cancer stem-like cells, Differentiated bulk tumor cells, Glioma stem-like cells, Differentiated glioma cells, Fibromodulin, Angiogenesis, Glioblastoma

## Abstract

Cancer stem cells alone can initiate and maintain tumors, but the function of non-cancer stem cells that form the tumor bulk remains poorly understood. Proteomic analysis showed a higher abundance of the extracellular matrix small leucine-rich proteoglycan Fibromodulin (FMOD) in the conditioned medium of non-cancer stem cells (DGCs; differentiated glioma cells) of glioma compared to that of glioma stem-like cells (GSCs). DGCs silenced for FMOD fail to cooperate with co-implanted GSCs to promote tumor growth. FMOD downregulation neither affects GSC growth and differentiation nor DGC growth and reprogramming *in vitro*. DGC-secreted FMOD promotes angiogenesis by activating Integrin-dependent Notch signaling in endothelial cells. Furthermore, conditional silencing of FMOD in newly generated DGCs *in vivo* inhibits the growth of GSC-initiated tumors due to poorly developed vasculature and increases mouse survival. Collectively, these findings demonstrate that DGC-secreted FMOD promotes glioma tumor angiogenesis and growth through paracrine signaling in endothelial cells and identifies a DGC-produced protein as a potential therapeutic target in glioma.

## Introduction

Tumors and their microenvironment form an ecosystem with many cell types that support tumor growth. The key constituents of this ecosystem include cancer stem-like cells (CSCs), non-cancer stem cells (non-CSCs) or differentiated cancer cells, and various other cell types that collectively make up the tumor stroma ***(Prager et al., 2019)***. It is well established that the tumor-initiating capacity lies solely with CSCs, thereby making them the crucial architects of tumor-stroma interactions that favor tumor growth and progression ***(Rheinbay et al., 2013)***. CSCs have a dichotomous division pattern, as they are capable of self-renewal and give rise to differentiated cells that form the bulk of the tumor ***(Olmeda and Ben Amar, 2019).*** The indispensable role of CSCs, which usually constitute only a minority population within tumors, is well documented in many solid tumors ***(Galli et al., 2004; Ignatova et al., 2002; Singh et al., 2004; Yang et al., 2020a)*.**

The tumor microenvironment is a vital driver of plasticity and heterogeneity in cancer ***(Carnero and Lleonart, 2016; Heddleston et al., 2010)***. The presence of hypoxic and necrotic regions is the hallmark of very aggressive tumors like glioblastoma (GBM), which have a highly vascular niche that supplies nutrients to cancer cells and makes a conducive environment for the tumor cells to thrive ***(Hambardzumyan and Bergers, 2015; Huang et al., 2016).*** Paracrine signaling mediated by proteins secreted from tumor cells, particularly glioma stem-like cells (GSCs), helps acquire this highly vascular phenotype by attracting blood vessels towards themselves and inducing pro-angiogenic signaling in endothelial cells through extracellular matrix remodeling ***(Dittmer and Leyh, 2014; Rupp et al., 2016).*** A reciprocal relationship exists between GSCs and endothelial cells by which endothelial cells induce stemness phenotype in cancer cells through activation of Notch, Sonic-Hedgehog, and Nitric Oxide Synthase signaling pathways ***(Jeon et al., 2014; Zhu et al., 2011)***, while GSCs drive vascularization of the tumor via endogenous endothelial cell stimulation, vascular mimicry, and GBM-endothelial cell transdifferentiation ***(Hardee and Zagzag, 2012; Soda et al., 2011)***. Recent reports have shown that CSCs induce such a high vascularization of tumors such as GBM by migrating along blood vessel scaffolds to invade novel vascular niches, thereby ensuring surplus and continuous blood supply at their disposal ***(Prager et al., 2020)***. In GBM, CD133+ and Nestin+ cells (representing GSCs) are located in close proximity of the tumor microvascular density (MVD), whereas a lower number of CD133- and Nestin-cells (representing differentiated glioma cells; DGCs) are located in the vicinity of the blood vessels. It has also been reported that the depletion of brain tumor blood vessels causes a decrease in the number of tumor-initiating GSCs ***(Calabrese et al., 2007)*.**

Besides CSC self-renewal, their differentiation to form the bulk cancer cells also plays a crucial role in tumor growth and maintenance ***(Jin et al., 2017).*** Epigenome unique to CSCs compared to differentiated cancer cells has been documented ***(Suva et al., 2014; Zhou et al., 2018).*** Reciprocally, a set of four reprogramming transcription factors, POU3F2, SOX2, SALL2, and OLIG2, is identified in GBM that are sufficient to reprogram DGCs and create the epigenetic landscape of native GSCs, thus creating “induced” cancer stem cells (***Suva et al., 2014***). The epigenetic regulation forms the basis of cellular plasticity, which creates a dynamic equilibrium between CSCs and differentiated cancer cells ***(Safa et al., 2015).*** Oncogene-induced dedifferentiation of mature cells in the brain was also reported using a mouse model of glioma, and the reprogrammed CSCs were proposed to contribute to the heterogeneous cell state populations observed in malignant gliomas ***(Friedmann-Morvinski et al., 2012; Friedmann-Morvinski and Verma, 2014***). Lineage tracing analyses revealed the reprogramming of DGCs to GSCs that act as a reservoir for initiating relapse of the tumors upon Temozolomide chemotherapy ***(Auffinger et al., 2014).*** Hypoxia has also been reported to reprogram differentiated cells to form CSCs in glioma, hepatoma, and lung cancer (***Prasad et al., 2017; Wang et al., 2017)***. Spontaneous conversion of differentiated cancer cells to CSCs has also been reported in breast cancer ***(Klevebring et al., 2014)*.**

Collectively, these studies highlight the crucial role of CSCs in cellular cross-talk in the tumor niche and establish CSCs as critical drivers of tumorigenesis. However, the massive imbalance in the proportions of CSCs and non-CSCs or differentiated cancer cells in tumors raises several important questions. Considering that differentiated cancer cells constitute the bulk of tumors, do they have specific functions, or do they only constitute the tumor mass? Do they contribute to the complex paracrine signaling occurring within the tumor microenvironment? Do they support tumor growth by promoting CSC growth and maintenance? It has been recently shown in GBM that DGCs cooperate with GSCs through a paracrine feedback loop involving neurotrophin signaling to promote tumor growth ***(Wang et al., 2018).*** While this study suggests a supporting role for differentiated cancer cells in tumor growth, the large proportion of them in tumors suggests a role in paracrine interactions with other stromal cells in the tumor niche.

We used quantitative proteomics to identify DGC-secreted proteins that might support their paracrine interactions within the tumor microenvironment. We show an essential role of Fibromodulin (FMOD) secreted by DGCs in promoting tumor angiogenesis via a cross-talk with endothelial cells. FMOD promotes Integrin-dependent Notch signaling in endothelial cells to enhance their migratory and blood vessel forming capacity. These findings indicate that DGCs are crucial for supporting tumor growth in the complex tumor microenvironment by promoting multi-faceted interactions between tumor cells and the stroma.

## Results

### DGC and GSC secretomes have distinct proteomes revealed by tandem mass spectrometry

While GSCs alone can initiate a tumor, the overall tumor growth requires functional interactions between GSCs and DGCs ***(Singh et al., 2004; Wang et al., 2018).*** To further understand the respective roles of GSCs vs. DGCs in tumor growth, we compared the conditioned medium (CM) derived from three patient-derived human GSC cell lines (MGG4, MGG6, and MGG8) ***(Wakimoto et al., 2009)*** and their corresponding DGCs, using a quantitative proteomic strategy. Proteins in CMs were systematically analyzed by nano-flow liquid chromatography coupled to Fourier transform tandem mass spectrometry (nano-LC-FT-MS/MS), and their relative abundance in DGC vs. GSC CM was determined by label-free quantification. We found that 119 proteins are more abundant in GSC CM, while 185 proteins are more abundant in the DGC CM (p<0.05, **Figure 1A; Supplementary Table S1**). Analysis of overrepresented functional categories among proteins exhibiting differential abundances in GSC vs. DGC CMs using Perseus with a p< value 0.05 revealed that the DGC CM is enriched in proteins known to exhibit extracellular or cell surface localization, such as proteins annotated as extracellular matrix (ECM) organization while terms related to DNA replication and many signaling pathways are enriched in GSC CM (**Supplementary Figures 1A and B**).

**Figure 1:**
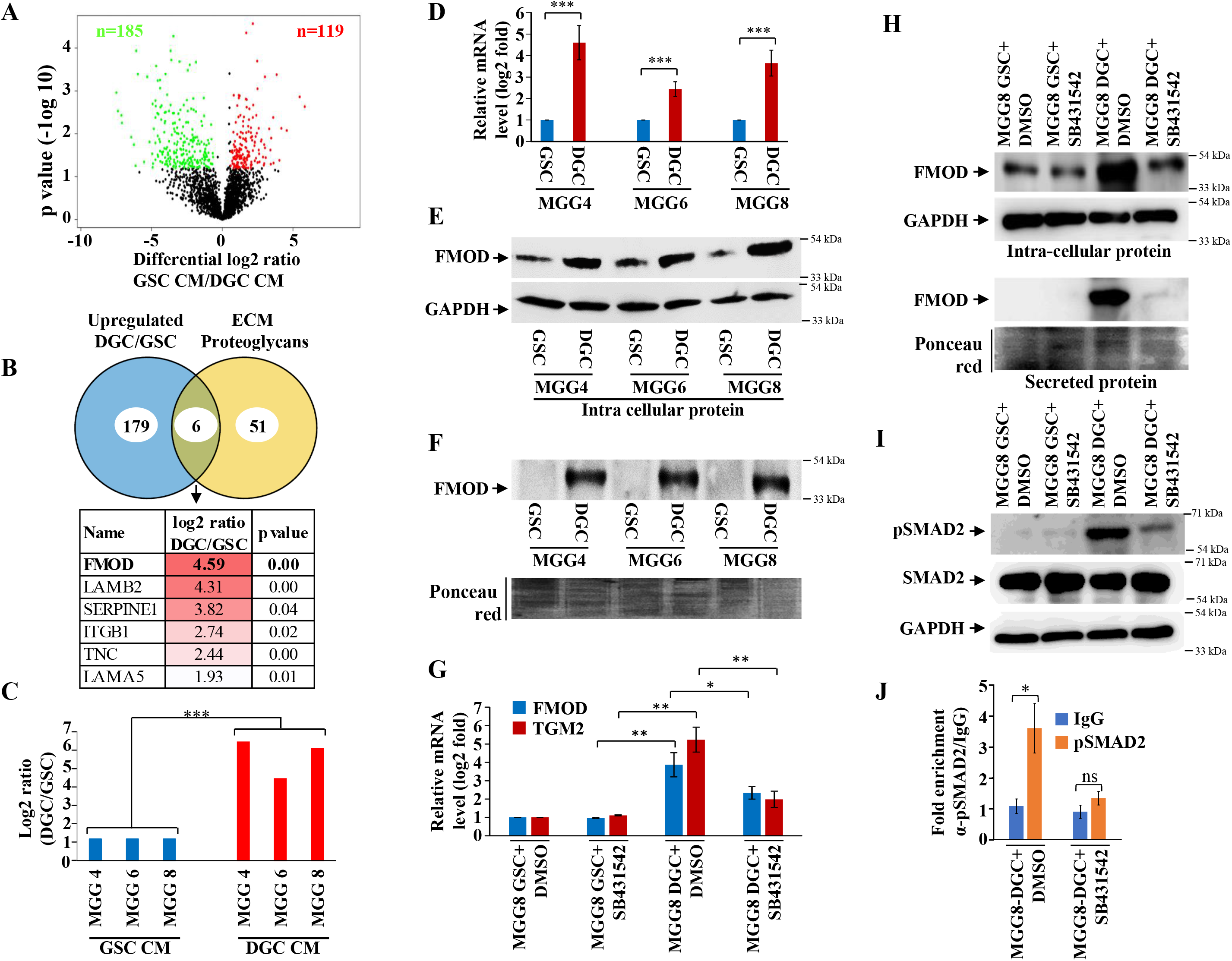
Quantitative proteomics shows higher abundance of fibromodulin under the control of TGFβ signaling in the DGC secretome. **A.** Volcano plot depicting relative protein abundance in GSC (MGG4, MGG6 and MGG8) *vs.* their corresponding DGC conditioned media (CMs). The black dots represent the non-significant proteins (p>0.05), while the red (higher abundance in GSC CM) and green (lower abundance in GSC CM) dots represent the significant ones (p<0.05) with a Log_2_ fold change cut-off of >0.58 or <-0.58. **B.** Venn diagram showing proteins upregulated in DGC CM and those annotated as ECM proteoglycans. Of the common proteins, FMOD exhibits the highest DGC/GSC ratio (indicated by the more intense red color). **C.** Label-free quantification (LFQ) of FMOD, expressed as Log_2_ fold change in GSCs *vs.* DGCs CM. **D.** Real-time qRT-PCR analysis shows upregulation of FMOD transcript in DGCs (red bars) *vs.* GSCs (blue bars). **E.** Western blotting shows the presence of higher amounts of intracellular FMOD in DGCs compared with corresponding GSCs. **F.** Top: Western blotting shows the presence of higher amounts of FMOD in the DGC CM *vs*. GSC CM. Bottom: Equal loading of the proteins assessed by Ponceau Red staining. **G.** Real-time qRT-PCR analysis showing showing reduction of high FMOD transcript level in DGCs upon treatment SB431542 (10 μM), a TGF-β inhibitor. Red bars indicate FMOD expression, and blue bars represent TGM2 (a bonafide TGF-β pathway target gene) expression. **H.** Western blotting shows that the high FMOD level in DGCs is inhibited by treating cells with SB431542 (10 μM); intracellular-top, and secreted-bottom. Equal loading of the secreted proteins assessed by Ponceau Red staining (bottom). **I.** Western blotting shows higher expression of pSAMD2 in DGCs than in GSCs, which is reduced by SB431542 treatment. **J.** Real-time qRT-PCR shows significantly higher fold enrichment of pSMAD2 in the FMOD promoter, which is inhibited upon SB431542 treatment (10 μM). for panels C, D, G, and J, p-value is calculated by unpaired t test with Welch’s correction. p value less than 0.05 is considered significant with *, **, *** representing p value less than 0.05, 0.01 and 0.001 respectively.

### TGFβ signaling controls the expression of Fibromodulin (FMOD) in DGCs

The enrichment of the “extracellular matrix” (ECM) annotation among proteins exhibiting higher abundance in DGC secretome prompted us to focus on ECM proteoglycans in line with their critical role in facilitating cancer cell signaling through their interaction with growth factor receptors, extracellular ligands and matrix components, and in promoting tumor-microenvironment interactions ***(Winkler et al., 2020)***. Six ECM proteoglycans were found to be more abundant in DGC CM compared with GSC CM (**Figure 1B**). The role of five of them (LAMB2, SERPINEE1, ITGB1, TNC, and LAMA5) in tumor growth has been well established ***(Angel et al., 2020; Bartolini et al., 2016; Long et al., 2016; Wang et al., 2021; Yang et al., 2020b).*** We thus focused on FMOD that exhibited the highest DGC CM/GSC CM protein ratio. FMOD is a small leucine-rich repeat proteoglycan upregulated in GBM due to the loss of promoter methylation orchestrated by TGFβ1-dependent epigenetic regulation ***(Mondal et al., 2017).*** FMOD promotes glioma cell migration through actin cytoskeleton remodeling mediated by an Integrin-FAK-Src-Rho-ROCK signaling pathway but does not affect colony-forming ability, growth on soft agar, chemosensitivity, and glioma cell proliferation ***(Mondal et al., 2017).*** We first confirmed the higher abundance of FMOD seen in DGC CM compared to GSC CM (**Figure 1C**) both at the transcript level (**Figure 1D)** and at the protein level (**Figures 1E and F**) in three GSC cell lines (MGG4, MGG6, and MGG8).

In line with our previous findings indicating that TGFβ signaling controls FMOD expression in glioma ***(Mondal et al., 2017),*** we next explored the possible role of this pathway in FMOD overexpression in DGCs. Gene Set Enrichment Analysis (GSEA) analysis of differentially regulated transcripts in GSC vs. DGC showed significant depletion of several TGFβ signaling pathway genes (**Supplementary Figure 2; Supplementary Table S2),** suggesting an enhanced TGFβ signaling in DGCs. Likewise, GSEA revealed an enrichment of several TGFβ signaling pathway genes in most GBM transcriptome datasets (**Supplementary Figure 3),** further supporting activation of TGFβ signaling in DGCs that represent the bulk of GBMs. In addition, the DGCs used in this study that express a high level of FMOD showed enrichment in mesenchymal signature, compared to GSCs (**Supplementary Figure 4**), consistent with the elevated TGF TGFβ signaling and FMOD levels we observed in the mesenchymal GBM subtype (**Supplementary Figures 5A and B**). Moreover, treating MGG8-DGCs with the TGFβ inhibitor (SB431542) significantly decreased luciferase activity of SBE–Luc (a TGFβ responsive reporter and contains Smad binding elements) and FMOD Promoter-Luc reporters (**Supplementary Figures 5C and D**). We also found higher levels of FMOD and TGM2 (a bonafide TGFβ target gene) transcripts and FMOD and pSMAD2 (an indicator of activated TGFβ signaling) proteins in MGG8-DGCs than MGG8-GSCs (**Figures 1G to 1I**). The addition of a TGFβ inhibitor (SB431542) significantly decreased transcript levels of FMOD and TGM2, and protein levels of FMOD and pSMAD2 in MGG8-DGCs (**Figures 1G to 1I**). Further, chromatin immunoprecipitation experiments revealed pSMAD2 occupancy on FMOD promoter in MGG8 DGCs that was significantly reduced by pretreating cells with SB431542 (**Figure 1J**). These results demonstrate a predominant expression and secretion of FMOD by DGCs that are promoted by TGFβ signaling.

### Tumor growth requires FMOD secreted by DGCs

We next investigated the role of FMOD in GSC and DGC growth and interconversion between both cell populations *in vitro* using two human (MGG8 and U251) and two murine (AGR53 and DBT-Luc) glioma cell lines. We found that the absence of FMOD neither affected GSC growth and differentiation to DGC (**Supplementary Figures 6 to 8**) nor DGC growth and reprogramming to form GSCs (**Supplementary Figures 9 to 11**; more details in **Supplementary information**), consistent with our previous findings showing that FMOD does not affect glioma cell proliferation *in vitro **(Mondal et al., 2017).*** In line with previous findings that DGCs cooperate with GSCs to promote tumor growth ***(Wang et al., 2018),*** we then evaluated the ability of DGCs silenced for FMOD to support the growth of tumors initiated by GSCs in co-implantation experiments in a syngeneic mouse model using GSCs and DGCs derived from DBT-Luc glioma cells. Reminiscent of our observations in MGG8 cell line, DBT-Luc-DGCs express higher levels of FMOD than DBT-Luc-GSCs (**Supplementary Figures 10C and D**). To silence the expression of FMOD in DBT-Luc-DGCs, we used a doxycycline-inducible construct that contains an inducible mCherry-shRNA downstream of the Tet-responsive element ***(Angel et al., 2020)*** (**Figure 2A**). The scheme of the co-implantation experiment is described on **Figure 2B**. DBT-Luc-GSC cells were coinjected with either DBT-Luc-DGC/miRNT (non-targeting shRNA) or DBT-Luc-DGC/miRFMOD (FMOD shRNA). In both groups, 50% of the mice received doxycycline on alternated days from day nine post-injection until the end of the experiment. Tumors in mice coinjected with DBT-Luc-GSCs and DBT-Luc-DGCs/miRNT grew much faster and reached a significantly larger size (measured by bioluminescence) than tumors in mice injected with DBT-Luc-GSCs alone, regardless of doxycycline treatment (**Figures 2B to 2D, compare black and purple lines with blue line; Supplementary Table S3**). Notably, mice treated with doxycycline did show mCherry expression in tumors (**Figure 2C**). In contrast, injected DBT-Luc-DGC/miRFMOD cells failed to support the growth of tumors initiated by DBT-Luc-GSCs in doxycycline-treated mice compared to doxycycline untreated mice (**Figures 2B to 2D; compare red line with orange line; Supplementary Table S3)**. While mice injected with DBT-Luc-GSCs + DBT-Luc-DGCs/miRFMOD (Dox+) showed an increase in tumor growth until the onset of doxycycline treatment (as seen in the rise in bioluminescence), subsequent tumor growth was drastically reduced. As expected, mice injected with DBT-Luc-DGCs alone developed substantially small tumors (**Figures 2C and D**). The small tumors formed in animals injected with either DBT-Luc-GSC + DBT-Luc-DGC/miRFMOD (Dox+) or DBT-Luc-GSC alone expressed significantly less FMOD protein than other tumors (**Supplementary Figure 12**). These results indicate that FMOD secreted by DGCs is essential for the growth of tumors initiated by GSCs.

**Figure 2:**
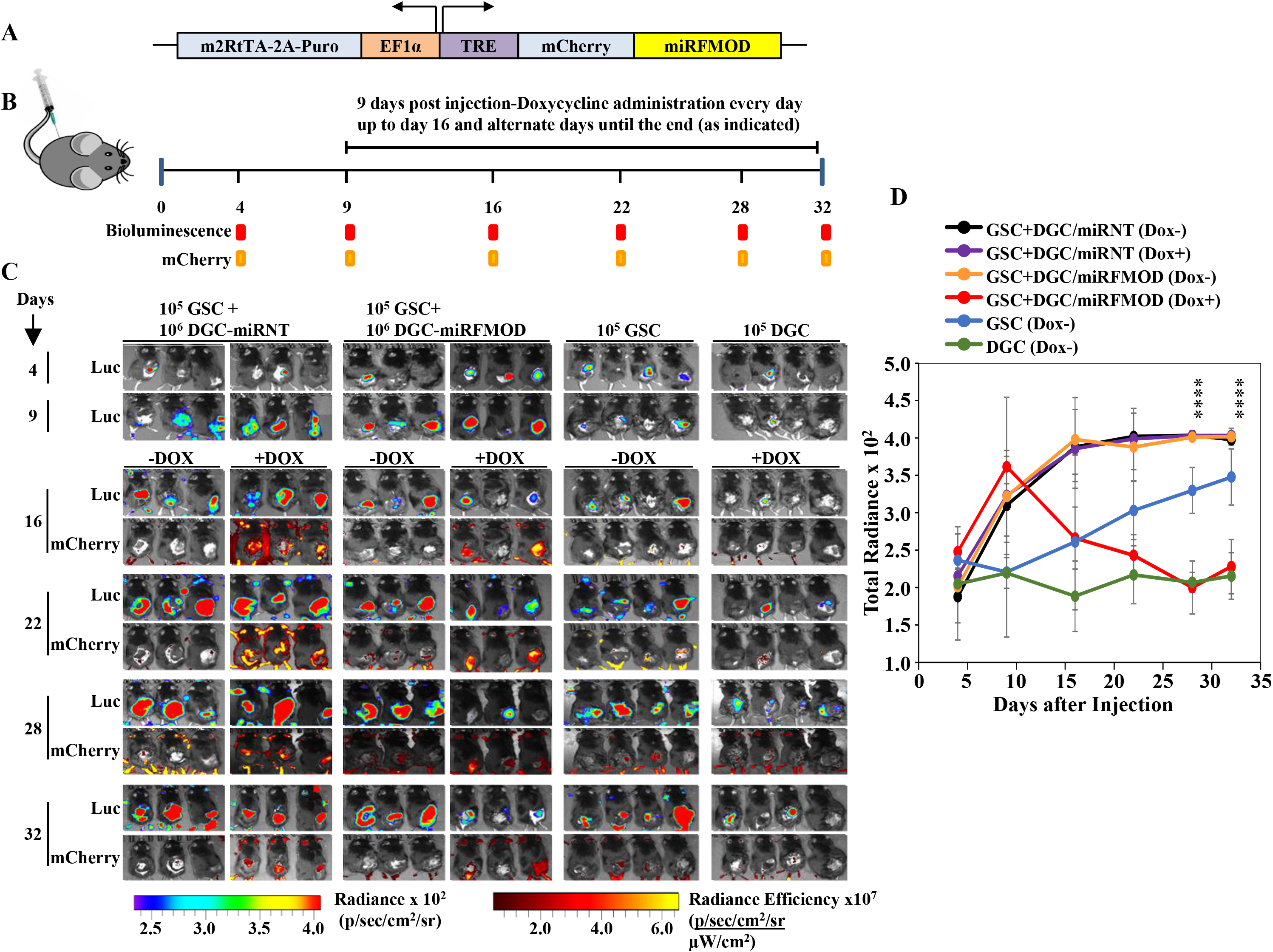
DGC-secreted FMOD is essential for tumor growth initiated by GSCs *in vivo* in a co-implantation experiment. **A.** Inducible shFMOD lentiviral construct. **B.** Schema depicting the GSC-DGC co-implantation experiment in C57BL/6 mice (n=5 for each group). Mice were injected with a combination of DBT-Luc-GSCs + DBT-Luc-DGCs (two groups for miRNT, Dox+ and Dox-. Same strategy for miRFMOD group.) The control groups were only injected with 10^5^ DBT-Luc-GSCs or 10^6^ DBT-Luc-DGCs and did not receive doxycycline. **C**. *In vivo* imaging of the injected mice, showing tumor growth over time by both bioluminescence and mCherry fluorescence, according to the timeline shown in **B. D.** Quantification of the total radiance. The different colors represent the different groups of animals. Significant differences between each of the groups were calculated using ANOVA. The p-values for days 28 and 32 are shown. A detailed comparison of the p values between different groups is provided in Supplementary Table 2.

### FMOD induces angiogenesis of host-derived and tumor-derived endothelial cells

Tumor cell interactions with stromal cells are critical for glioma tumor growth ***(Pine et al., 2020).*** Small leucine-rich proteoglycans such as FMOD promote angiogenesis in the context of cutaneous wound healing ***(Pang et al., 2019)***. In addition, we previously found a significant enrichment of the term “Angiogenesis” among differentially regulated genes in FMOD-silenced U251 glioma cells (***Mondal et al., 2017***). In light of these observations, we next examined the impact of FMOD on tumor angiogenesis. First, we tested the ability of glioma cell-derived FMOD to induce angiogenic network formation by immortalized human pulmonary microvascular endothelial cells (ST1). We used LN229 and U251 glioma cells, which express low and high levels of FMOD, respectively, for overexpression and silencing studies ***(Mondal et al., 2017)***. We found that the CM derived from LN229 cells stably expressing FMOD (LN229/FMOD) induced more angiogenesis than LN229/Vector stable cells (**Figures 3A and B**). Further, the CM of FMOD-silenced U251 cells was less efficient in promoting angiogenesis than the CM of cells expressing non-targeting siRNA (**Figure 3C; Supplementary Figure 13A)** or shRNA (**Supplementary Figure 13B**). The addition of recombinant human FMOD (rhFMOD) to the CM of U251/siFMOD cells rescued its ability to induce angiogenesis (**Figure 3C; Supplementary Figure 13A**). More importantly, the addition of rhFMOD directly to endothelial cells induced angiogenesis in the presence of a control antibody (IgG) but not in the presence of an FMOD neutralizing antibody (**Supplementary Figure 13C**).

**Figure 3:**
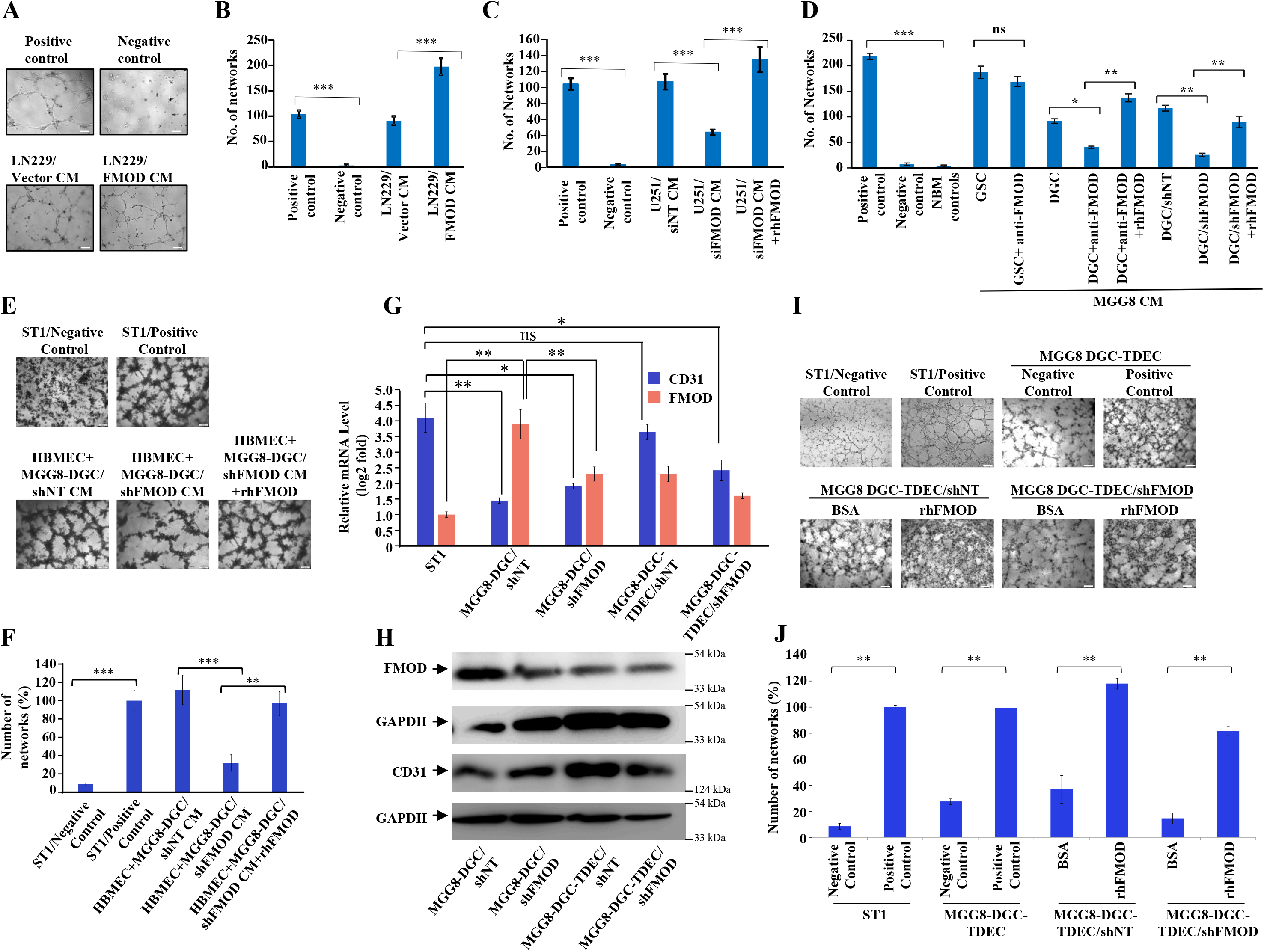
DGC-secreted FMOD promotes angiogenesis. **A.** Representative images of *in vitro* network formation by ST1 cells treated with LN229/Vector CM *vs.* LN229/FMOD CM. In the positive control condition (top left), cells are plated in complete endothelial cell media (Medium 199) supplemented with Endothelial Cell Growth Factors (ECGS) and 20% FBS, and in the negative control (top right), where cells are plated in incomplete Medium 199 (without serum and ECGS). Significantly more networks are formed when ST1 cells are treated with LN229/FMOD CM (right bottom), compared with cells treated with LN229/Vector CM (left bottom). Magnification 10X, Scale bar = 100 μm. **B**. Quantification of the number of complete networks formed in **A. C.** Quantification of the number of networks formed by ST1 cells shows that cells treated with U251-DGC/siFMOD (low FMOD) CM form a significantly lesser number of networks than cells treated with U251-DGC/siNT (high FMOD) CM. This reduction in the number of networks is rescued by addition of rhFMOD (400 nM) in U251-DGC/siFMOD CM. **D.** Quantification of the number of networks formed in the *in vitro* angiogenesis assay showing that DGC-secreted FMOD induces network formation by ST1 cells, which is reduced when an FMOD-neutralizing antibody is added to the CM and rescued by adding rhFMOD. Furthermore, MGG8-DGC/shFMOD CM forms lesser networks than MGG8-DGC/shNT CM, which is rescued by adding rhFMOD to the MGG8-DGC/shFMOD CM. **E.** Representative images of *in vitro* network formation by primary human brain-derived microvascular endothelial cells (HBMECs). Significant reduction in the number of networks formed when the cells are treated with MGG8-DGC/shFMOD CM compared with MGG8-DGC/shNT CM, which is rescued by the exogenous addition of rhFMOD. Magnification 4X, Scale bar = 200 μm. **F.** Quantification of the number of complete networks formed in **E. G.** Real-time qRT-PCR analysis showing CD31 (blue bars) and FMOD (orange bars) expression in ST1, MGG8-DGC/shNT and MGG8-DGC/shFMOD cells before and after transdifferentiation (the groups labeled as TDECs represent the transdifferentiated cells). **H.** Western blotting analysis showing the expression of FMOD and CD31 in MGG8-DGC/shNT and MGG8-DGC/shFMOD cells before and after transdifferentiation (the groups labeled as TDECs represent the transdifferentiated cells). **I.** Representative images of *in vitro* network formation by MGG8-DGC/shNT TDECs and MGG8-DGC/shFMOD TDECs upon BSA and rhFMOD treatments. Magnification 4X, Scale bar = 200 μm. **J.** Quantification of the number of complete networks formed in **I.** For panels B, C, D, F, G, and J, p-values were calculated by unpaired t-test with Welch’s correction. p values < 0.05 were considered significant with *, **, *** representing p values < 0.05, 0.01 and 0.001 respectively. ns stands for non-significant.

Both CMs derived from three DGCs and their corresponding GSCs also induced angiogenesis efficiently (**Supplementary Figure 13D**). Further, pretreating cells with an FMOD antibody significantly reduced the ability of MGG8-DGC CM, but not that of MGG8-GSC CM, to induce angiogenesis (**Figure 3D**). The reduced ability of the FMOD antibody-pretreated DGC CM to promote angiogenesis was rescued by the exogenous addition of an excess of rhFMOD (**Figure 3D**). Moreover, CM derived from FMOD-silenced MGG8-DGCs was less efficient in promoting angiogenesis than the CM of shNT (**Figure 3D**) or siNT (**Supplementary Figure 13E)** transfected MGG-DGC cells. The effect of FMOd silencing was rescued by adding exogenous rhFMOD (**Figure 3D; Supplementary Figure 13E**). Both rhFMOD and FMOD present in the CM collected from MGG8-DGC induced the migration and invasion but not the proliferation of ST1 cells (**Supplementary Figures 14A to 14E).** Further, CM from MGG8-DGC/shNT cells was more efficient than CM from MGG8-DGC/shFMOD cells to promote angiogenic network formation by human brain-derived primary endothelial cells (HBMECs) and mouse brain-derived immortalized endothelial cells (B.End3) (**Figures 3E and F; Supplementary Figure 14F**). Again, the effect of FMOD silencing was rescued by adding rhFMOD (**Figures 3 E and F**; **Supplementary Figure 14F).**

Vascular mimicry (VM) is one of the alternative mechanisms of angiogenesis wherein tumor-derived endothelial cells (TDECs) originate from glioblastoma cells ***(Angara et al., 2017; Ricci-Vitiani et al., 2010; Soda et al., 2011).*** To assess the ability of FMOD to induce TDECs derived from DGCs to form angiogenic networks, we used MGG8-DGC and U87 cells. MGG8-DGC/shNT cells grown in endothelial media (M199) under hypoxia (1% O_2_) differentiated to TDECS as evidenced by an increase in CD31 (**Figures 3 G and H**). MGG8-DGC/shFMOD also differentiated to form TDECs, albeit with less efficiency (**Figures 3G and H**). Further, the addition of rhFMOD induced both MGG8-DGC/shNT-TDEC and MGG8-DGC/shFMOD-TDEC cells to form angiogenic networks efficiently (**Figures 3 I and J**). Similarly, U87 cells differentiated to TDECs (**Supplementary Figures 15A and B),** which readily formed angiogenic networks in the presence of rhFMOD **(Supplementary Figures 15 C and D).** Collectively, these results demonstrate that DGC CM can induce angiogenesis and identify FMOD as a critical mediator of DGC-induced angiogenesis.

### FMOD activates Integrin/FAK/Src-dependent Notch pathway in endothelial cells to induce angiogenesis

To dissect the signaling mechanisms underlying FMOD-induced angiogenesis, we subjected protein extracts derived from ST1 endothelial cells treated or not with rhFMOD to Reverse Phase Protein Array (RPPA). A total of 12 proteins exhibited differential abundance in a time-dependent manner in rhFMOD-treated ST1 endothelial cells (**Figure 4A**). These include HES1, a downstream target of the Notch signaling pathway that has been shown to promote angiogenesis (***Zhao et al., 2017***). We thus investigated the possible involvement of Notch signaling in FMOD-induced angiogenesis. The addition of rhFMOD induced luciferase activity of Notch-dependent CSL-Luc and HES-Luc reporters in ST1 cells, but not in γ-secretase inhibitor- (GSI; a Notch pathway inhibitor) pretreated cells (**Supplementary Figures 16A and B**). The addition of rhFMOD also increased HES1 mRNA and protein levels in ST1 cells, an effect abolished by GSI pretreatment of cells **(Figures 4B and C).** rhFMOD treatment also resulted in the translocation of NICD (Notch intracellular domain) from the cytosol to the nucleus, as shown by subcellular fractionation and confocal microscopy **(Supplementary Figures 16C and D**). Furthermore, rhFMOD failed to induce angiogenic network formation by GSI-pretreated ST1 cells (**Figure 4D**). In addition, ST1 cells having a stable expression of NICD (ST1/NICD) showed enhanced angiogenic network formation than ST1 vector stable (ST1/Vector) cells (**Supplementary Figures 17A to 17D).** FMOD present in CM from MGG8-DGCs induced ST1/Vector cells, but not ST1/NICD cells to form more angiogenic networks, suggesting that Notch activation in endothelial cells is an essential step in FMOD-induced angiogenesis.

**Figure 4:**
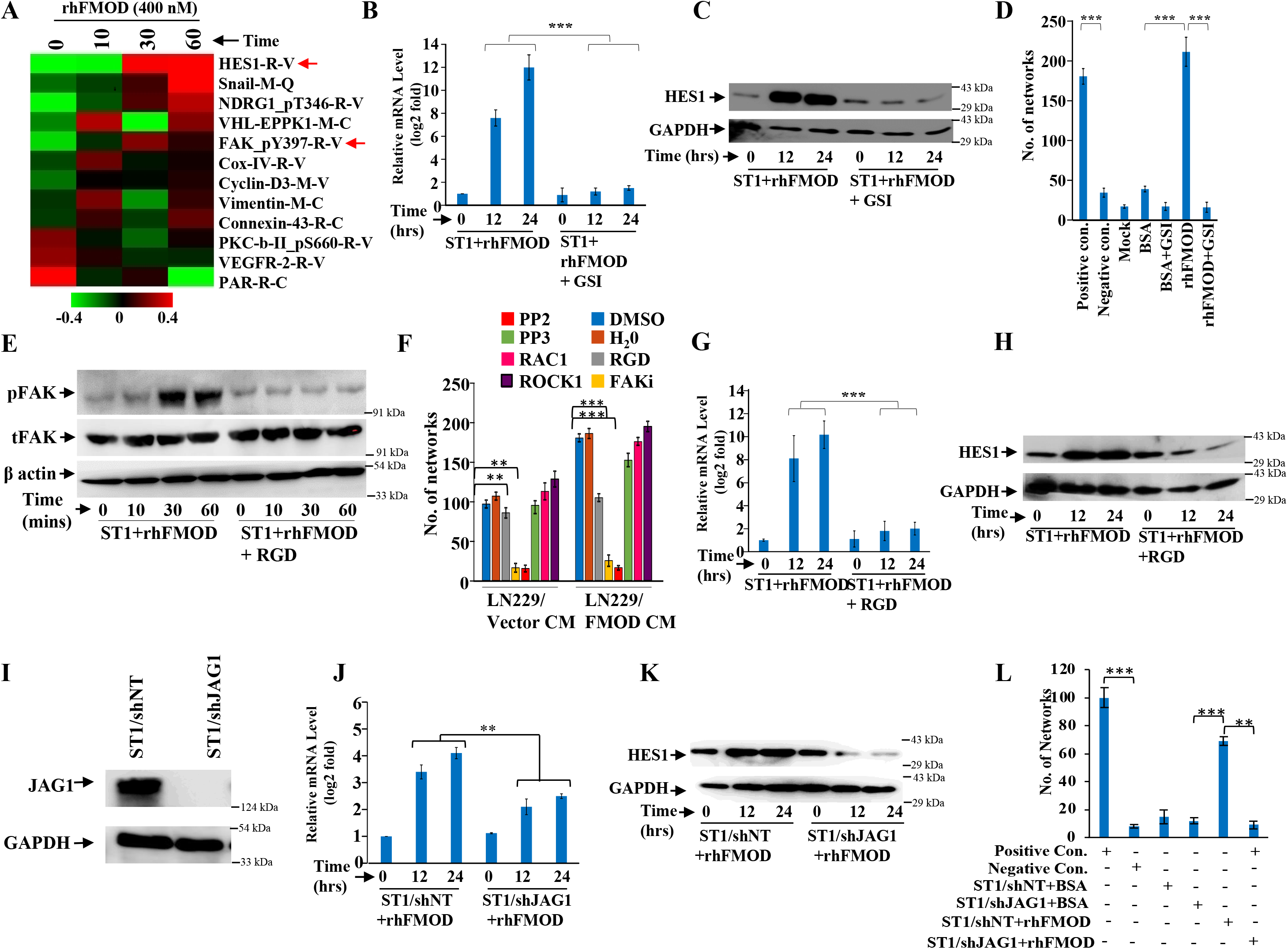
Integrin/FAK/Src/JAG1-dependent Notch pathway activation in endothelial cells mediates FMOD-induced angiogenesis. **A.** Heatmap showing differentially regulated proteins in endothelial cells (treated with vehicle or 400 nM rhFMOD for 10, 30 and 60 min, Log_2_ fold change >/< 0.2), assessed by RPPA (Reverse Phase Protein Array). Red and green depict upregulated and downregulated proteins in ST1 cells, respectively. The red arrows indicate HES1 and pFAK. **B.** qRT-PCR analysis shows that rhFMOD treatment of ST1 cells causes a time-dependent increase in HES1 mRNA, which is inhibited in cells pre-treated with GSI (10 μM). **C.** Western blotting showing rhFMOD treatment of ST1 cells causes an increase in HES1 protein, which is inhibited in cells pre-treated with GSI. **D.** Quantification of the number of networks formed in *in vitro* angiogenesis assay shows that the rhFMOD-mediated increase in the number of networks formed by ST1 cells is abolished in cells pre-treated with GSI. **E.** Western blotting shows a time-dependent increase in phospho-FAK level in FMOD-treated ST1 cells. This increase is suppressed when the cells are pre-treated with RGD peptide (10 μM), an Integrin inhibitor. **F.** Quantification of networks upon treatment of ST1 cells with either LN229/Vector CM or LN229/FMOD CM. ST1 cells are pretreated with the indicated inhibitors (PP2, PP3, and FAK inhibitor-PF573228 were used at a concentration of 10 μM, ROCK1 inhibitor-H1152 was used at a concentration of 0.5 μM, Rac1 inhibitor was used at a concentration of 10 μM). **G.** qRT-PCR analysis shows that rhFMOD treatment of ST1 cells causes a time-dependent increase in HES1 mRNA, which is inhibited in cells are pre-treated with RGD peptide. **H.** Western blotting showing rhFMOD treatment of ST1 cells causes an increase in HES1 protein level, which is inhibited in cells pre-treated with the RGD peptide. **I**. Western blotting validating the knockdown of JAG1 in shJAG1-transfected ST1 cells. **J.** qRT-PCR analysis shows that rhFMOD-induced expression of HES1 mRNA is significantly decreased in ST1/shJAG1 cells compared with ST1/shNT cells. **K.** rhFMOD-induced expression of HES1 protein is decreased in ST1/shJAG1 cells compared with ST1/shNT cells. **L.** Quantification of networks formed in *in vitro* angiogenesis assay upon treatment of ST1/shNT and ST1/shJAG1 cells with BSA or rhFMOD. For panels B, D, F, G, J, and L, the p-value calculated by unpaired t-test with Welch’s correction are indicated. p-value less than 0.05 was considered significant with *, **, *** representing p-value less than 0.05, 0.01 and 0.001 respectively.

The increase in phosphorylated FAK (pFAK, FAK_Py397-R-V; the molecule downstream of Integrin signaling) levels in rhFMOD-treated endothelial cells, as shown by RPPA (**Figure 4A**), also suggested a possible role of integrin signaling in FMOD-induced angiogenesis. This is consistent with our previous findings indicating that FMOD activates Integrin signaling *via* type I collagen to engage the FAK-Src-Rho-ROCK pathway and promote the migration of glioma cells ***(Mondal et al., 2017).*** We first confirmed the activation of Integrin signaling by FMOD, as assessed by increased pFAK in rhFMOD-treated ST1cells, but not in cells pretreated with RGD peptide, an Integrin inhibitor (**Figure 4E**). The addition of RGD peptide inhibited angiogenesis induced by LN229/FMOD CM (**Figure 4F**). Likewise, angiogenesis induced by LN229/FMOD CM was completely inhibited when ST1 cells were pretreated with inhibitors of FAK (FAKi) or Src (PP2), two signaling molecules downstream of Integrin (**Figure 4F**). These treatments also strongly reduced the basal level of angiogenesis elicited by the CM of LN229/Vector cells. In contrast, an inactive analog of Src inhibitor (PP3), as well as inhibitors of RAC1 and ROCK, failed to inhibit the ability of CM derived from LN229/FMOD cells to induce angiogenesis (**Figure 4F**). Our previous report also demonstrated that the interaction of FMOD with type I collagen is essential for integrin activation ***(Mondal et al., 2017).*** The C-terminal region of FMOD comprises 11 leucine-rich-repeats (LRRs), of which the 11^th^ repeat binds to type I collagen ***(Oldberg et al., 2007).*** A synthetic interfering peptide (**RLDGN*E*I*K*R**) corresponding to the 11^th^ LRR of type 1 collagen, but not a modified peptide (**RLDGN*Q*I*M*R**), competes with rhFMOD for binding to type I collagen to activate integrin signaling in glioma cells ***(Mondal et al., 2017; Oldberg et al., 2007).*** Consistent with these findings, rhFMOD-induced luciferase activity of CSL-Luc and HES-Luc (**Supplementary Figures 18A and B**) and angiogenesis by ST1 cells (**Supplementary Figure 18C**) were significantly inhibited by the interfering peptide, but not the modified peptide, suggesting a crucial role of type I collagen-dependent activation of Integrin signaling in FMOD-induced angiogenesis. To identify the α and β subunits of integrin involved in FMOD-mediated activation of integrin signaling in endothelial cells, we chose ITGA6, ITGB1, and ITGAV for investigation based on analysis of transcriptome data derived from laser capture dissected microvessels from the human brain (more detail in **Supplementary information**). Silencing either of the selected three integrin subunits in ST1 cells reduced significantly the ability of rhFMOD to activate integrin as assessed by reduced pFAK levels (**Supplementary Figures 18B to 18I**), thus demonstrating the involvement of αv/β1 and α6/β1 heterodimeric integrin receptors in FMOD activation of integrin signaling in endothelial cells.

Next, to examine a possible cross-talk between Integrin and Notch signaling in FMOD-treated endothelial cells, we tested the effect of the RGD peptide on the ability of rhFMOD to induce Notch signaling. Pretreatment of ST1 cells with RGD peptide significantly reduced rhFMOD-elicited CSL-Luc and HES-Luc activation (**Supplementary Figures 4E and F**). Likewise, pretreatment of cells with FAKi or PP2, but not PP3, significantly reduced rhFMOD-induced CSL-Luc and HES-Luc activity in ST1 cells **(Supplementary Figures 19A to 19D).** Further, rhFMOD failed to increase HES1 transcript and protein levels in ST1 cells treated with either RGD peptide, FAKi, or PP2, but not in cells treated with PP3 **(Figures 4G and H; Supplementary Figures 19E to 19H)**.

We next investigated the mechanistic link between Integrin and Notch signaling in FMOD-treated ST1 cells. Since Notch activation by FMOD is sensitive to GSI treatment, we explored activation of Notch ligands by Integrin-FAK signaling in FMOD-treated cells. While the addition of rhFMOD induced DLL3 and JAG1 transcripts in ST1 cells, RGD peptide pretreatment abolished JAG1 induction (**Supplementary Figure 20A**), suggesting that JAG1 might be a potential linking molecule. Consistently, pretreatment of cells with either FAKi or PP2, but not PP3, also abolished the ability of rhFMOD to induce JAG1 transcript in ST1 cells (**Supplementary Figures 20B and C)**. Further, JAG1 silencing significantly decreased rhFMOD-induced activity of CSL-Luc and HES-Luc reporters and HES1 transcript/protein levels in ST1 cells (**Supplementary Figures 20D and E; Figures 4I to 4K**). JAG1 silencing also abolished the ability of rhFMOD to induce angiogenic networks by ST1 cells (**Figure 4L**). In addition, rhFMOD induced expression of the FAK inducible transcription factor KLF8 mRNA in ST1 cells, an effect abolished by pretreating cells with RGD peptide (**Supplementary Figure 20F)**. Further, KLF8 silencing in ST1 cells significantly reduced the ability of rhFMOD to increase pFAK level (**Supplementary Figures 20G and H**), suggesting that KLF8 activates JAG1 through an integrin-dependent pathway in FMOD-treated endothelial cells. Collectively, these results identify JAG1 as a molecular link between Integrin and Notch signaling pathways in FMOD-treated endothelial cells and demonstrate a key role of Integrin-FAK-JAG1-Notch-HES1 signaling in FMOD-induced angiogenesis.

To further explore the clinical relevance of these findings, we interrogated transcriptome datasets from multiple sources (more detail in **Supplementary Information**). We found a significant upregulation of transcript levels of FMOD, JAG1 and HES1 in GBM from various datasets (**Supplementary Figures 21A to 21C).** We also found a positive correlation between FMOD and HES1 transcripts and between FMOD and JAG1 transcripts in the majority of GBM datasets analyzed (**Supplementary Figures 21D and E)**, which further substantiates the functional link between FMOD, JAG1, and HES1. We also show that high FMOD transcript levels and hypomethylation of FMOD promoter are associated with poor prognosis in most datasets (**Supplementary Figures 22F and G)**. These observations provide additional support for the activation of Integrin-Notch signaling in FMOD-treated endothelial cells.

### DGC-secreted FMOD is required for the growth of murine and human GSC-initiated tumors

While GSCs alone can initiate a tumor, tumor growth requires continuous differentiation to form DGCs, which form the bulk of the tumor mass. In line with our co-implantation experiments (**Figure 2**), we sought to define the importance of FMOD secreted by DGCs generated through a differentiation program initiated by GSCs during tumor growth *in vivo,* using a syngenic intracranial glioma mouse model. We injected AGR53-GSC-miRNT and AGR53-GSC-miRFMOD cells intracranially into C57/black mice and allowed them to form tumors. Thirteen days after intracranial injections, both groups received doxycycline as indicated (**Figure 5A**). The understanding is that as the tumors start growing, GSCs, in addition to their self-renewal, will start differentiating *de novo* to form DGCs, which would express high levels of FMOD. However, doxycycline treatment would inhibit FMOD expression, and thus one could investigate the importance of DGC-secreted FMOD in tumor growth. AGR53-GSC/miRNT and AGR53-GSC/miRFMOD cell-initiated tumors showed a similar size seven days after doxycycline treatment as shown by mCherry fluorescence (**Figures 5B and C; day 21)**. However, upon subsequent follow-up, doxycycline administration significantly inhibited the growth of AGR53-GSC/miRFMOD-initiated tumors over time but not that of AGR53-GSC/miRNT tumors (**Figures 5B, C, and E**). Further, doxycycline administration increased the survival of mice injected with AGR53-GSC/miRFMOD cells compared to AGR53-GSC/miRNT cells (**Figure 5D**). AGR53-GSC/miRFMOD initiated tumors showed decreased FMOD, mCherry, and GFP expression compared to AGR53-GSC/miRNT initiated tumors (**Figures 5F and G**). DBT-Luc GSC/miRFMOD, another murine glioma GSC cell line, produced similar results: FMOD silencing after doxycycline administration resulted in reduced tumor growth (**Supplementary Figures 23A and B**), increased mice survival (**Supplementary Figures 23C**), and decreased FMOD expression (**Supplementary Figure 23D**).

**Figure 5.**
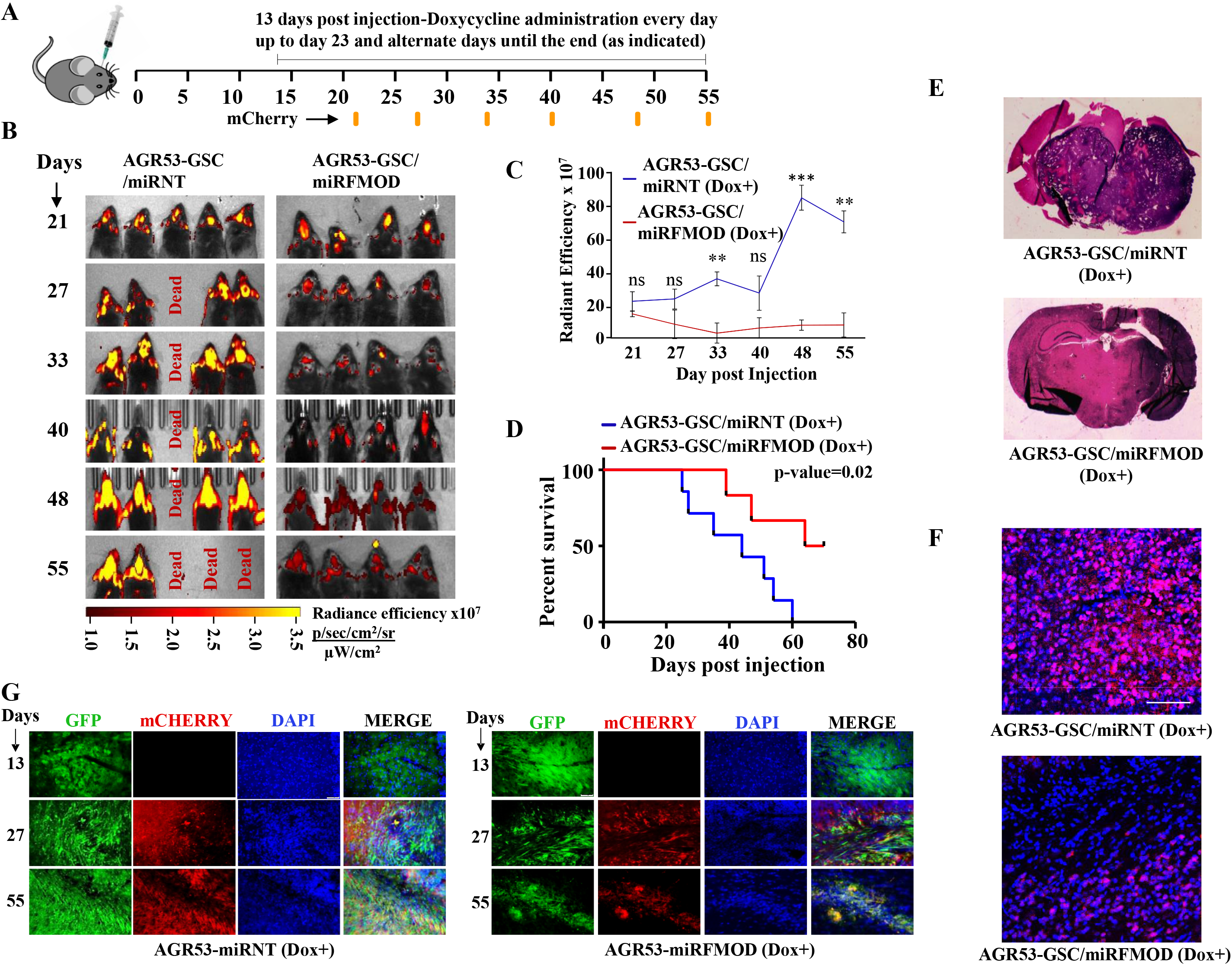
Conditional silencing of FMOD in DGCs formed *de novo* by GSC-initiated tumors inhibits tumor growth. **A**. Schema is showing the timeline of the *in vivo* experiments. AGR53-GSCs (stably expressing miRNT or miRFMOD) were injected (1 x 10^5^ cells per animal) on Day 0. Tumors were allowed to grow till day 13, and then doxycycline (100 μg/animal) administration was started. Please note, *in vitro* characterization show that the highest knockdown of FMOD was obtained on the 7^th^ day after doxycycline administration. First, *in vivo* imaging for mCherry expression depicting tumor size was done on day 21 post-injection, followed by imaging at regular intervals (as noted by the orange marks). **B.** *In vivo* fluorescence imaging of mice injected with either AGR53-GSC/miRNT or AGR53-GSC/miRFMOD cells. **C.** Radiance Efficiency for each time point in the two groups of animals was plotted as a measure of tumor size at the indicated days. **D**. Kaplan-Meier graphs showing the survival of mice bearing AGR53-GSC/miRNT (Dox+) and AGR53-GSC/miRFMOD (Dox+) cells. **E.** Haematoxylin and Eosin staining show a larger tumor (depicted by dark blue color due to extremely high cell density) in mice brain injected with AGR53-GSC/miRNT (Dox+) cells (top), compared to that of AGR53-GSC/miRFMOD (Dox+) cells (bottom). Magnification=0.8 X **F**. Immunohistochemical analysis showing FMOD expression in brains of mice injected with AGR53-GSC/miRNT (Dox+) and AGR53-GSC/miRFMOD (Dox+) cells. Red indicates FMOD, and Blue indicates H33342. The merged images have been shown for representation. Magnification = 20x, Scale= 50 μm **G.** Brain sections showing areas of fluorescence for both AGR53-GSC/miRNT (Dox+) (left panel) and AGR53-GSC/miRFMOD (Dox+) (right panel) groups of animals. Please note that AGR53 cell lines stably express GFP. mCherry expression is induced upon doxycycline addition. On day 13, prior to the administration of doxycycline, both miRNT(left) and miRFMOD (right) do not have any mCherry expression but have almost equal GFP expression. However, over time after the onset of doxycycline administration, both mCherry and GFP expression decreased in miRFMOD group but not in the miRNT group. Merged images show an overlap of GFP and mCherry-positive tumor areas. Magnification=20x, Scale= 50μm. For panels C and D, p-value less than 0.05 was considered significant with *, **, *** representing p-value less than 0.05, 0.01, and 0.001, respectively.

To determine the relevance of our findings for the human pathology, we investigated the importance of DGC-secreted FMOD in the growth of tumors initiated by MGG8 and U251 cells using a xenograft mouse glioma model. The enrichment of cancer stem cells in MGG8 GSC neurospheres was confirmed by the higher expression of CD133, compared to MGG8 DGCs (**Figure 6A**). We established orthotopic xenografts using MGG8-GSC/shNT and MGG8-GSC/shFMOD cells. Reminiscent of results obtained using transplantation of murine glioma cells, MGG8-GSC/shNT-transplanted mice readily developed intracranial tumors, whereas MGG8-GSC/shFMOD mice showed impaired tumor formation and increased mice survival (**Figures 6B, and C**). Immunostaining and confocal microscopy analysis showed high expression of FMOD in MGG8-GSC/shNT while FMOD was barely detectable in MGG8-GSC/shFMOD tumors (**Figure 6D**). Likewise, U251/shFMOD cells that showed reduced FMOD protein level compared to U251/shNT (**Supplementary Figure 11A)**, developed smaller tumors (**Figures 6E to 6H**), and mice bearing U251/shFMOD tumors had longer survival than those carrying U251/shNT tumors (**Figure 6G)**. As expected, the expression of FMOD was more elevated in U251/shNT tumors than in U251/shFMOD tumors (**Figure 6I**). Collectively, these findings indicate that DGC-secreted FMOD is essential for the growth of both human and mouse glioma.

**Figure 6:**
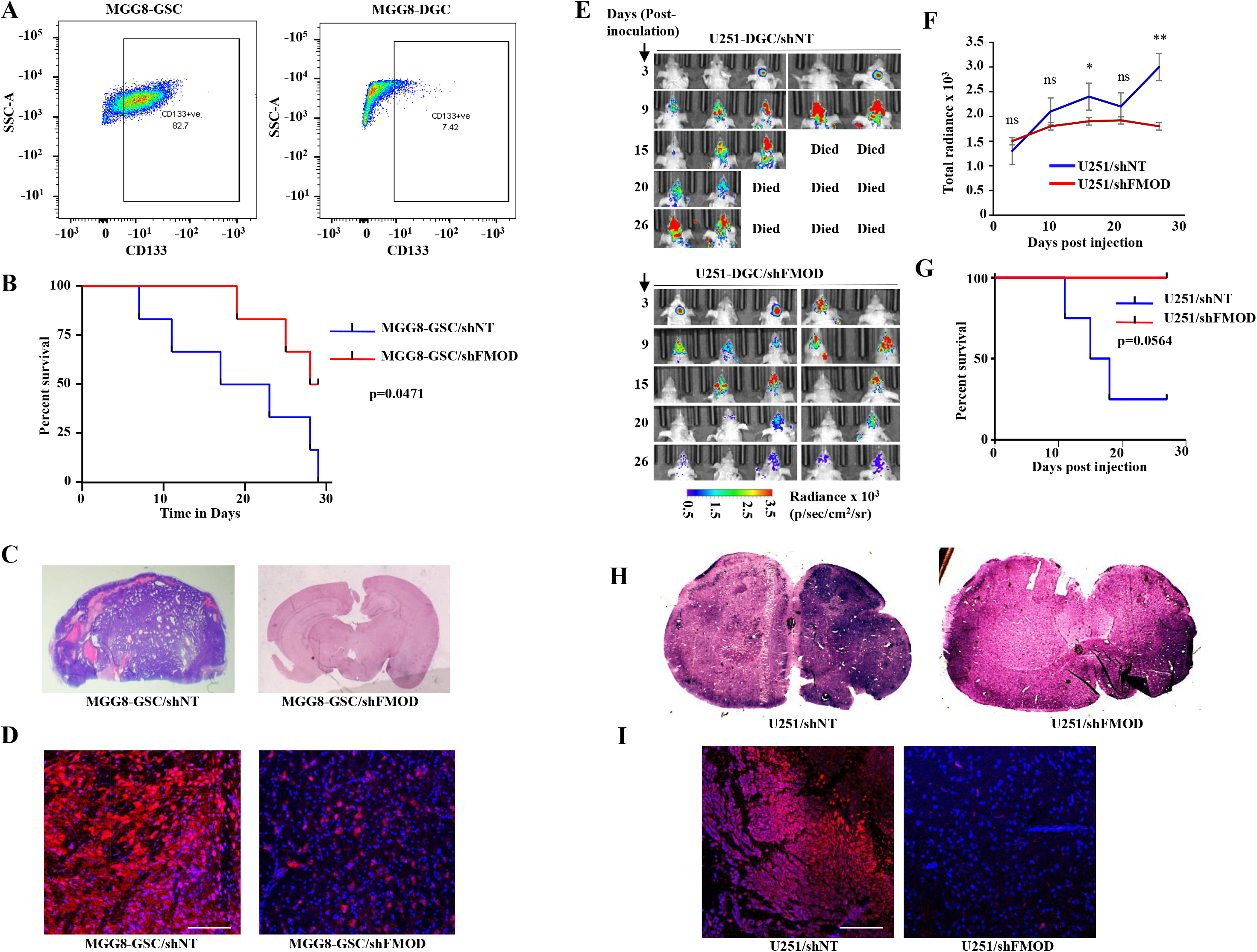
Growth of human GSC-initiated tumors require secreted FMOD. **A.** Flow-cytometry analysis showing enrichment of CD133-positive cells in MGG8-GSCs compared to MGG8-DGCs. **B.** Kaplan-Meier graphs showing the survival of mice injected with MGG8-GSC/shNT or MGG8-GSC/shFMOD cells. **C.** Haematoxylin and Eosin staining showing tumors in brains of mice injected with MGG8-GSC/shNT and MGG8-GSC/shFMOD cells. Magnification= 0.8 X **D.** Immunohistochemical labeling showing FMOD expression in brains of mice injected with MGG8-GSC/shNT or MGG8-GSC/shFMOD cells. Red indicates FMOD, and Blue indicates H33342. The merged images have been shown for representation. Magnification = 20x, Scale, 50 μm. **E.** *In vivo* bioluminescence imaging of two groups of animals injected with either U251/shNT or U251/shFMOD cells. The tumor formation was followed over 26 days. **F.** Total radiance for each time point was plotted as an index of tumor size for both animal groups. **G.** Kaplan-Meier graphs showing the survival of mice injected with U251-DGC/shNT or U251-DGC/shFMOD cells. **H.** Haematoxylin and Eosin staining showing tumors in brains of mice injected with U251-DGC/shNT or U251-DGC/shFMOD cells. Scale= 0.8 X. **I.** Immunohistochemical analysis showing FMOD expression in brains of mice injected with U251-DGC/shNT or U251-DGC/shFMOD cells. Red indicates FMOD, and Blue indicates H33342. The merged images have been shown for representation. Magnification = 20x, Scale = 50 μm. For panels B, F, and G, a p-value less than 0.05 was considered significant with *, **, *** representing p values less than 0.05, 0.01, and 0.001, respectively. ns stands for non-significant.

### Reduced angiogenesis in small tumors formed in FMOD-silenced conditions

Next, we investigated the mechanism underlying reduced tumor growth in FMOD-silenced conditions. First, we measured the expression of CD133 and GFAP markers as the representation of GSCs and DGCs in the tumors formed in the animal models. The CD133 positive cells are much less in proportion compared to GFAP positive cells in tumors formed by AGR53-GSC, DBT-Luc-GSCs, and MGG8-GSCs under FMOD non-targeting conditions (**Supplementary Figures 24A, and B, 25A and B, and 26A and B**), in good correlation to the low proportion of GSCs seen in brain tumors (**Singh et al., 2004, Galli et al., 2004, and Calabrese et al., 2007**). Further, most GFAP positive cells are also positive for FMOD expression compared to CD133 positive cells (**Supplementary Figures 24A and C, 25A and C, and 26A and C**), recapitulating the results we obtained *in vitro*, where FMOD is expressed specifically by DGCs (**Figure 1**). We also found a similar expression pattern of CD133 and GFAP markers in small tumors formed under FMOD-silenced conditions in all three tumor models (**Supplementary Figures 24D and E, 25D and E, and 26D and E).** The GFAP staining in these tumors confirms the occurrence of an efficient differentiation program even in FMOD-silenced conditions, which confirms our results obtained *in vitro*, where the absence of FMOD failed to affect the GSC differentiation to form DGCs **(Supplementary Figures 6 and 8)**. As expected, the small tumors formed in FMOD-silenced conditions showed substantially reduced FMOD staining (**Supplementary Figures 24D and F, 25D and F, and 26D and F).**

We next evaluated the extent of blood vessel formation by measuring the endothelial cell marker immunostaining in the tumors. The small size tumors formed by AGR53-GSC/miRFMOD cells after doxycycline treatment showed reduced staining for CD31 and vWF (von Willebrand factor) compared to AGR53-GSC/miRNT tumors (**Figures 7A to 7C, and Supplementary Figures 27A to 27C**). Tumors formed by DBT-Luc-GSC/miRFMOD cells in doxycycline-treated mice also showed significantly reduced CD31 staining compared to that measured in the absence of doxycycline treatment (**Supplementary Figures 27D to 27F**). Reminiscent of murine glioma tumors, tumors induced by MGG8-GSC/shFMOD cells also showed reduced CD31 staining compared to MGG8-GSC/shNT cells (**Figures 7D to 7F**). We then tested the extent of blood vessel formation by TDECs. In all three tumor models (AGR53-GSCs, DBT-Luc-GSCs, and MGG8-GSCs), blood vessels formed by TDECs were significantly reduced in tumors formed in FMOD-silenced conditions (**Figures 7G to 7L; Supplementary Figures 27G to 27I**). These findings confirm our previous results obtained *in vitro*, where the absence of FMOD decreased the ability of host-derived endothelial cells and TDECs to form angiogenic networks (**Figure 3; Supplementary Figures 13 to 15**). Next, we investigated the involvement of Integrin-FAK-JAG1-Notch-HES1 signaling in FMOD-induced angiogenesis in the context of glioma tumors. Confocal microscopy analysis in tumors formed by MGG8-GSCs, AGR53-GSCs, and DBT-Luc-GSCs revealed a significant colocalization of the endothelial cell marker CD31 with FMOD, pFAK, JAG1, and HES1 in blood vessels (**Supplementary Figures 28A to 28C**). From these results, we conclude that angiogenesis induced by DGC-secreted FMOD is essential for glioma tumor growth.

**Figure 7:**
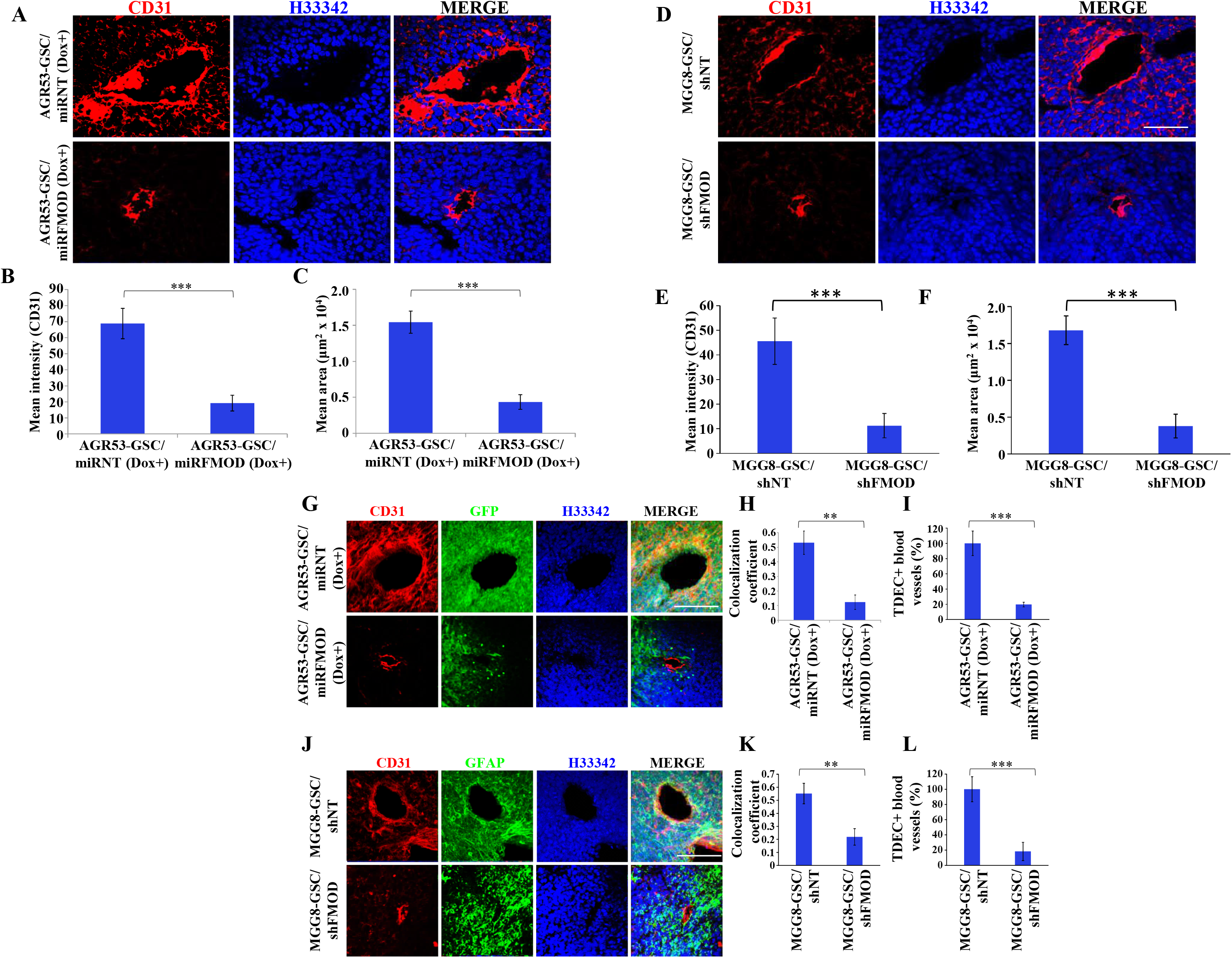
Reduced angiogenesis is characteristic of tumors initiated by FMOD-silenced glioma cells. **A.** Immunohistochemical analysis showing blood vessels lined by cells expressing CD31 in brains of mice injected with AGR53-GSC/miRNT or AGR53-GSC/miRFMOD cells after doxycycline administration. **B.** Quantification of the mean fluorescence intensity of CD31 in brains of mice injected with AGR53-GSC/miRNT or AGR53-GSC/miRFMOD cells after doxycycline administration. **C.** Quantification of the mean area of blood vessels in brains of mice injected with AGR53-GSC/miRNT or AGR53-GSC/miRFMOD cells after doxycycline administration. **D.** Immunohistochemical analysis showing CD31 expression in brains of mice injected with MGG8-GSC/shNT or MGG8-GSC/shFMOD. **E.** Quantification of the mean fluorescence intensity of CD31 in brains of mice injected with MGG8-GSC/shNT or MGG8-GSC/shFMOD cells. **F.** Quantification of the mean, mean area of blood vessels in brains of mice injected with MGG8-GSC/shNT or MGG8-GSC/shFMOD cells. **G.** Immunohistochemical analysis showing overlap of CD31 (red) and GFP (green) expression in brains of mice injected with AGR53-GSC/miRNT or AGR53-GSC/miRFMOD cells after doxycycline administration. The yellow arrow indicates the region exhibiting colocalization of both markers. **H.** Quantification of the colocalization coefficient in the AGR53-GSC/miRNT and AGR53-GSC/miRFMOD groups of animals after doxycycline injection. **I.** Quantification of the TDEC+ blood vessels, indicating the measure of vascular mimicry (VM) in the AGR53-GSC/miRNT and AGR53-GSC/miRFMOD groups. **J.** Immunohistochemical analysis showing overlap of CD31 (red) and GFAP (green) expression in brains of mice injected with MGG8-GSC/shNT or MGG8-GSC/shFMOD cells. The yellow arrow indicates the region exhibiting colocalization of both markers **K.** Quantification of the colocalization coefficient in the MGG8-GSC/shNT and MGG8-GSC/shFMOD groups. **L.** Quantification of the TDEC+ blood vessels, indicating the measure of vascular mimicry (VM) in the MGG8-GSC/shNT and MGG8-GSC/shFMOD groups. **M.** A model depicting how GBM tumors are made up of a small proportion of GSCs, and a massive number of DGCs that form the tumor bulk. FMOD, primarily secreted by DGCs, upregulates JAG1 through the activation of integrin signaling in ST1 cells. The higher expression of JAG1 causes the activation of the Notch signaling pathway, which results in the transcriptional upregulation of HES1 in endothelial cells. The integrin-dependent Notch pathway activation promotes angiogenesis and vascular mimicry, leading to glioma tumor growth. For panels A, D, G, and J, the magnification used is 20 X (Scale = 50 μm). For panels B, C, E, F, H, I, K, and L, a p-value less than 0.05 was considered significant with *, **, *** representing a p-value less than 0.05, 0.01, and 0.001, respectively.

## Discussion

Cellular hierarchy is well established in GBM. The importance of GSCs in tumor initiation, growth, immune escape, angiogenesis, invasion into the normal brain and resistance to therapy is also well established ***(Bao et al., 2006; Wakimoto et al., 2009)***. GSCs are known for their ability to self-renew and to differentiate to form DGCs, the bulk cells of tumors (***Galli et al., 2004; Ignatova et al., 2002; Singh et al., 2004; Suva et al., 2014; Yang et al., 2020a***). However, the role of DGCs in tumor growth remains poorly understood. The key requirement of tumor cells for self-maintenance in a novel tumor niche is the supply of nutrients. GSCs are known to promote the establishment of a highly vascularized microenvironment by being in close physical contact with endothelial cells ***(Calabrese et al., 2007)***. The massive proportion of DGCs in the tumor suggests that GSC-initiated angiogenesis might not be sufficient to meet the large nutrient requirement of the entire tumor. Wang *et al*. demonstrated that DGC-secreted BDNF is essential for GSC growth and maintenance through DGC-GSC paracrine signaling, which highlights the crucial role of DGC-secreted proteins in tumor formation (***Wang et al., 2018***). Further, a possible interaction between DGCs and stromal cells, such as endothelial cells, cannot be ruled out. We hypothesized that in addition to GSCs, DGCs might play an essential role in autocrine and paracrine signaling involving different types of cells to augment tumor growth. The present study demonstrates the existence of paracrine signaling between DGCs and endothelial cells, which promotes angiogenesis and glioma tumor growth.

Previously, we have demonstrated that FMOD is highly expressed in GBMs compared to normal brain tissues. The loss of FMOD expression hampers the migratory function of glioma cells but has no impact on glioma cell proliferation ***(Mondal et al., 2017).*** This study identifies that FMOD is expressed exclusively by DGCs and further showed that FMOD is not needed for GSC/DGC growth and their plasticity to form one from the other. Given these facts, the inability of DGCs silenced for FMOD to support the growth of tumors initiated by GSCs was unexpected. However, it enlightened us that DGC-secreted FMOD has some essential yet unidentified functions supporting GSC-initiated tumor growth.

Here, we explored the mechanisms underlying the role of DGCs acting as critical support for GSC-initiated tumor growth. We demonstrate that DGC-secreted FMOD promotes tumor growth by inducing angiogenesis through integrin-dependent Notch signaling in endothelial cells, thus highlighting the importance of DGCs in tumor-stroma interactions that contribute to a sustainable niche for tumor growth. We further investigated the mechanism by which FMOD activates integrin-dependent Notch signaling in endothelial cells. Based on RPPA data which showed upregulation of HES1 in endothelial cells upon FMOD treatment, we demonstrated that activation of Notch signaling is essential for FMOD-induced angiogenesis. The importance of Notch signaling in glioma, especially in GSC growth and angiogenesis, is well documented ***(Bazzoni and Bentivegna, 2019; Stockhausen et al., 2010; Teodorczyk and Schmidt, 2014)***. Similarly, activation of Notch signaling in endothelial cells has been involved in tumor development and angiogenesis ***(Gridley, 2007; Kofler et al., 2011; Phng and Gerhardt, 2009).*** RPPA data also showed an increase in pFAK in FMOD-treated glioma cells. Our previous study showed that FMOD acts on glioma cells via the Integrin-FAK-Src-Rho axis to promote migration ***(Mondal et al., 2017).*** The present study demonstrates that FMOD-activated Integrin signaling is essential for Notch pathway activation in ST1 cells. Integrin signaling in endothelial cells has been shown to play a crucial role in angiogenesis (***Short et al., 1998; Silva et al., 2008***). Based on our results indicating that FMOD-activated Notch signaling in endothelial cells is inhibited by GSI, we predicted that FMOD-elicited activation of the Notch pathway could involve Notch ligand-dependent process. Our experiments identified that JAG1 is the linker molecule that integrates Integrin signaling to the Notch pathway. We also found a significant colocalization of endothelial cell marker CD31 with FMOD, pFAK, JAG1, and HES1 in blood vessels of tumors formed in mouse models. Finally, endothelial cells stably expressing NICD showed enhanced angiogenic network formation in an FMOD-independent manner, suggesting that Notch activation is an essential step in FMOD-induced angiogenesis. Thus, our results show that Integrin-FAK-Src-KLF8-JAG1-dependent Notch signaling activation in endothelial cells mediates FMOD-induced angiogenesis.

Finally, in an orthotopic intracranial GBM mouse model, we show that conditional silencing of FMOD in newly generated DGCs during tumor growth leads to a significant reduction of tumor growth. Supporting our *in vitro* data indicating that FMOD is not required for GSC growth and their differentiation, we found that the small tumors formed in FMOD-silenced conditions show differentiation of GSCs. However, these tumors exhibited poorly developed blood vasculatures of host-derived endothelial cells and TDECs. The lower tumor burden in the absence of FMOD might be attributed to insufficient nutrient supply to sustain tumor growth due to the reduced blood vessel density. Thus, our study establishes an essential role of paracrine signaling between the DGCs and the stroma in the context of tumor growth in the natural tumor niche complexity. It also demonstrates the importance of FMOD secreted by DGCs in promoting human glioma tumor growth in a mouse model. We propose a tumor evolution model (**Figure 7M**), whereby GSCs, in addition to their self-renewal, continuously differentiate to form DGCs, which secrete protein factors like FMOD that mediate paracrine signaling in the different cell types of the tumor, thus creating a niche favorable to tumor growth.

In conclusion, the present study demonstrates that in addition to GSCs, DGCs have an essential role in tumor growth and maintenance. While the therapy-resistant and self-renewing GSCs trigger the early events of transformation and growth, DGCs, the proportion of which continues to increase during tumor growth, progressively become essential. Thus, targeting both CSCs and differentiated cancer bulk cells is vital to achieving a durable therapeutic response. The study also highlights the potential of GSC and DGC CM analysis to uncover novel targets in cancer therapy and the critical influence of DGC-secreted FMOD in glioma tumor growth.

## Author Contributions

SS carried out the experiments, MM carried out bioinformatic analyses, AM helped in some experiments related to cell culture, KRP and PM^2^ helped in animal experiments, SU carried out mass spectrometry experiments, DFM and PM^4^ corrected the manuscript and helped in planning experiments, KS wrote the manuscript and executed the whole study.

## Acknowledgments

The results published here are in whole or part based upon data generated by The Cancer Genome Atlas pilot project established by the NCI and NHGRI. Information about TCGA and the investigators and institutions that constitute the TCGA research network can be found at http://cancergenome.nih.gov/. We acknowledge the shRNA consortium (Dr. Subba Rao), IISc, India, for shRNA constructs. SS acknowledges IISc for fellowship. KS acknowledges CEFIPRA DBT, DST, and CSIR (Govt. of India) for research grants. Infrastructure supported by DST FIST, DBT-IISc partnership program, and UGC is acknowledged. KS is awarded J. C. Bose Fellowship from DST. This research was supported in part by grants to DFM from the Israel Science Foundation (Grant no.1315/15 and 1429/20). PM was supported by grants from Fondation pour la Recherche Médicale and CEFIPRA (n° IFC/5603-C/2016/503). Mass spectrometry experiments were carried out using the facilities of the Montpellier Proteomics Platform (PPM, BioCampus Montpellier).

## MATERIALS AND METHODS

### RESOURCE AVAILABILITY

Further information and requests for resources and reagents should be directed to and will be fulfilled by the lead contact Dr. Kumaravel Somasundaram.

### MATERIALS AVAILABILITY

This study did not generate any new unique reagents.

### DATA AND AVAILABILITY

Label-free mass spectrometry data between the GSC and DGC showing protein ratios in the GSC and DGC secretome and p values are shown on Supplementary Table S1 for proteins exhibiting significant differences in abundance in both conditions. The mass spectrometry data obtained the current study are available from the corresponding author on request.

### EXPERIMENTAL MODEL AND SUBJECT DETAILS

Experiments were performed in C57&BL/6J female mice and Athymic Nude female mice (6-8 week old) following the approved by the Institute Ethical Committee for Animal Experimentation. The mice were kept in a 12h light and dark cycle, fed *ad libitum* with a normal diet and the experiments were done in the light phase of the cycle.

#### Cell lines used

The GBM adherent cell lines LN229 and U251 were purchased from Sigma Aldrich, Saint Louis, Missouri, USA. Primary human tumour-derived GSCs MGG4, MGG6, and MGG8 were kindly gifted by Dr. Wakimoto (Massachusetts General Hospital, Boston, USA). AGR53 mouse-derived cell line is described before ***(Angel et al., 2020).*** DBT-Luc cells were a kind gift from Dr. Dinesh Thotala, Washington University in St. Louis St Louis, Missouri, United States. ST1 endothelial cells were a kind gift from Dr. Ron Unger, Johannes Gutenberg University, Germany. B.End3 cell line purchased from The American Type Culture Collection (ATCC, #CRL-2299). The HBMECs were purchased from Cell Biologics, USA (#H-6023).

#### Plasmids

The RAR3G vector (in which miRNT and miRFMOD are cloned for the inducible shRNA experiments) is previously described (***Friedmann-Morvinski et al., 2012***). shRNA for FMOD (miRFMOD) was cloned following the mir30 (miRNT) inducible backbone under the TRE (Tetracycline response element) promoter which is placed downstream of mCherry reporter gene. The m2RtTA transactivator is expressed from the same vector in the opposite direction under the EF1α promoter, and following a 2A peptide, the puromycin gene is placed for selection *in vitro*. The FMOD overexpression plasmid was bought from Origene, USA (#LY419579). NICD pCMV Neo/intracellular domain of human Notch1 (NIC-1) and CSL Luc were kind gifts from Prof. Thomas Kandesch, Department of Genetics, University of Pennsylvania School of Medicine. Hes-Luc plasmid was bought from Addgene, USA (#43806).

#### Antibodies used

##### Primary antibodies

FMOD (Abgent AP9243b, 1:2000), FAK (Cell Signaling Technology 3285S, 1:1000), pFAK (Cell Signaling Technology #3283S, 1:500), GAPDH (Sigma #G8795, 1:20,000), Actin (Sigma A3854, 1:20,000), HES1 (Cell Signaling Technology #D6P2U, 11988S, 1:1000), vWF (Abcam #6994, 1:2000), CD31 (Cell Signaling Technology #89C2, Mouse mAb 1:200 for IHC), JAG1 (Cell Signaling Technology, Jagged1 (D4Y1R) XP^®^ Rabbit mAb #70109, 1:200 for IHC, 1:1000 for WB), FMOD (Fibromodulin Polyclonal Antibody PA5-26250, Invitrogen, IHC 1:100, WB 1:1000), CD133 (Recombinant Anti-CD133 antibody-EPR20980-104 #ab216323 abcam, Flow Cyt. 1:100, Prom1 Monoclonal Antibody-2F8C5, #MA1-219, IHC 1:100), SOX2 (Cell Signaling Technology #3579 Rabbit mAb, IHC 1:100), GFAP ( Anti-GFAP antibody ab7260, abcam ICC 1:200, Recombinant Anti-GFAP antibody [EPR1034Y] - Mouse IgG2a (ab279290), IHC 1:200), NICD (NOTCH1 (Cleaved Val1744) Polyclonal Antibody PA5-99448, WB 1:200), SMAD2 (Smad2 (D43B4) XP^®^ Rabbit mAb #5339, Wb 1:1000), pSMAD2 (Phospho-SMAD2 (Ser465/Ser467) (E8F3R) Rabbit mAb #18338, WB 1:1000), Integrin beta-1 (Cell Signaling Technology #9699 Rabbit mAb, WB 1:1000), Integrin alphaV (Cell Signaling Technology #4711 Rabbit Ab, WB 1:1000), Integrin alpha-6 (Recombinant Anti-Integrin alpha 6 antibody [EPR18124] (ab181551), WB 1:500), KLF8 (Anti-KLF8 antibody (ab168527), WB 1:500)

##### Secondary antibodies

Goat Anti mouse HRP conjugate (Biorad #170-5047, WB 1:5000), Goat anti rabbit (H+L) secondary HRP conjugate (Invitrogen, #31460, WB 1:5000), Goat anti-Mouse IgG (H+L) Highly Cross-Adsorbed Secondary Antibody, Alexa Fluor 488) (Invitrogen, #A-11029), Goat anti-Rabbit IgG (H+L) Highly Cross-Adsorbed Secondary Antibody, Alexa Fluor 488 (Invitrogen, # Catalog # A-11034), Goat anti-Mouse IgG (H+L) Highly Cross-Adsorbed Secondary Antibody, Alexa Fluor 594 (Invitrogen, # Catalog # A-11032), Goat anti-Rabbit IgG (H+L) Highly Cross-Adsorbed Secondary Antibody, Alexa Fluor 594 (Invitrogen, # Catalog # A-11037), Goat anti-Rabbit IgG (H+L) Highly Cross-Adsorbed Secondary Antibody, Alexa Fluor 405 Plus (Invitrogen, # A48254), Goat anti-Mouse IgG (H+L) Highly Cross-Adsorbed Secondary Antibody, Alexa Fluor 405 Plus (Invitrogen, # Catalog # A48255). All Alexafluor conjugated antibodies were used at a dilution of 1:500 for IHC and ICC.

#### Resource Table

| REAGENT or RESOURCE | SOURCE | IDENTIFIER |
| --- | --- | --- |
| **Antibodies** |  |  |
| Rabbit polyclonal anti-FMOD | Abgent | Cat# AP9243b<br>RRID:AB_10612142 |
| Rabbit polyclonal anti-FAK | Cell Signaling<br>Technology | Cat#T9026; RRID:<br>AB_477593<br>RRID:AB_2269034 |
| Rabbit polyclonal anti-phospho FAK (Tyr397) | Cell Signaling<br>Technology | Cat# 3283,<br>RRID:AB_2173659 |
| Mouse Anti-GAPDH Monoclonal Antibody,<br>Unconjugated, Clone GAPDH-71.1 | Sigma-Aldrich | Cat# G8795,<br>RRID:AB_1078991 |
| Mouse Anti-Actin, beta Monoclonal Antibody,<br>Horseradish Peroxidase Conjugated, Clone AC-15 | Sigma-Aldrich | Cat# A3854,<br>RRID:AB_262011 |
| Rabbit Monoclonal HES1 (D6P2U) antibody | Cell Signaling<br>Technology | Cat# 11988,<br>RRID:AB_2728766 |
| Rabbit polyclonal Von Willebrand Factor antibody | Abcam | Cat# ab6994,<br>RRID:AB_305689 |
| Mouse monoclonal CD31 antibody | Cell Signaling<br>Technology | Cat#89C2<br>RRID:AB_2160882 |
| JAG1 Rabbit monoclonal antibody | Cell Signaling<br>Technology | Cat#70109<br>RRID: AB_2799774 |
| FMOD Rabbit polyclonal antibody | Invitrogen | Cat#PA5-26250<br>RRID: AB_2543750 |
| CD133 Recombinant antibody | Abcam | Cat#ab216323 RRID: AB_2847920 |
| Prom1 Monoclonal antibody | Invitrogen | Cat#MA1-219<br>RRID: AB_2725113 |
| SOX2 Rabbit monoclonal antibody | Cell Signaling Technology | Cat#3579<br>RRID: AB_2195767 |
| GFAP Mouse antibody | Abcam | Cat#ab279290<br>RRID: AB_1209224 |
| GFAP Recombinant antibody | Abcam | Cat#ab7260<br>RRID: AB_305808 |
| NOTCH1 Polyclonal antibody | Invitrogen | Cat#PA5-99448<br>RRID: AB_2818381 |
| SMAD2 Rabbit monoclonal antibody | Cell Signaling Technology | Cat#5339<br>RRID: AB_10626777 |
| pSMAD2 Rabbit monoclonal antibody | Cell Signaling Technology | Cat#18338<br>RRID: AB_2798798 |
| Integrin beta-1 Rabbit monoclonal antibody | Cell Signaling Technology | Cat#9699<br>RRID: AB_11178800 |
| Integrin alpha-V Rabbit monoclonal antibody | Cell Signaling Technology | Cat#4711<br>RRID: AB_2128178 |
| Integrin Alpha 6 antibody | Abcam | Cat#ab181551<br>RRID: |
| KLF8 antibody | Abcam | Cat#ab168527<br>RRID: |
| Goat Anti mouse HRP conjugate | Biorad | Cat#170-5047<br>RRID: AB_11125753 |
| Goat anti rabbit (H+L) secondary HRP conjugate | Invitrogen | Cat#31460<br>RRID: AB_228341 |
| Goat anti-Mouse IgG (H+L) Highly Cross-Adsorbed Secondary Antibody, Alexa Fluor 488 | Invitrogen | Cat#A-11029<br>RRID: AB_138404 |
| Goat anti-Rabbit IgG (H+L) Highly Cross-Adsorbed Secondary Antibody, Alexa Fluor 488 | Invitrogen | Cat#A-11034<br>RRID: AB_2576217 |
| Goat anti-Mouse IgG (H+L) Highly Cross-Adsorbed Secondary Antibody, Alexa Fluor 594 | Invitrogen | Cat#A-11032<br>RRID: AB_2534091 |
| Goat anti-Rabbit IgG (H+L) Highly Cross-Adsorbed Secondary Antibody, Alexa Fluor 594 | Invitrogen | Cat#A-11037<br>RRID: AB_2534095 |
| Goat anti-Rabbit IgG (H+L) Highly Cross-Adsorbed Secondary Antibody, Alexa Fluor 405 Plus | Invitrogen | Cat#A48254<br>RRID: AB_2890548 |
| Goat anti-Mouse IgG (H+L) Highly Cross-Adsorbed Secondary Antibody, Alexa Fluor 405 Plus | Invitrogen | Cat#A48255<br>RRID: AB_2890536 |
| **Biological samples** |  |  |
| Healthy adult C57BL/6 brain tissue | In this study |  |
| GBM adult C57BL/6 brain tissue | In this study |  |
| Healthy adult nude mice brain tissue | In this study |  |
| GBM adult nude mice brain tissue | In this study |  |
| GBM adult C57BL/6 subcutaneous tumor tissue | In this study |  |
| **Chemicals, peptides, and recombinant proteins** |  |  |
| γ-Secretase inhibitor (GSI) | Merck | 565750 |
| RGD peptide (Integrin inhibitor) | Sigma-Aldrich | **A8052** CAS Number: <u>99896-85-2</u> |
| FAK inhibitor (PF-573228) | Sigma-Aldrich | PZ0117 CAS Number: <u>869288-64-2</u> |
| PP2 (Src inhibitor) | Sigma-Aldrich | **P0042**; CAS Number: 172889-27-9 |
| PP3 (negative control, structural analogue of PP2) | Sigma-Aldrich | 529574 CAS 5334-30-5 |
| ROCK1 inhibitor | Sigma-Aldrich | 555550 CAS 871543-07-6 |
| RAC1 inhibitor | Sigma-Aldrich | 553502 CAS 1177865-17-6 |
| TGFB R1 inhibitor SB431542 | Sigma-Aldrich | 616464 CAS 301836-41-9 |
| Fibromodulin (FMOD) (NM_002023) Human Recombinant Protein | Origene | **CAT#: TP306534** |
| **Deposited data** |  |  |
| Label-free mass spectrometry performed with conditioned media from MGG4, MGG6 and MGG8 GSC vs DGC. | This paper | Supplementary Table S1 |
| **Experimental models: cell lines** |  |  |
| Human cell line: LN229 | Sigma Aldrich |  |
| Human cell line: U251-MG | Sigma Aldrich | 09063001 |
| Human cell line: MGG4, MGG6. MGG8 | Gift from Dr. Wakimoto (Massachusetts General Hospital, Boston, USA). |  |
| Mouse Cell line: AGR-53 | Lab of Dr. Dinorah Friedmann-Morvinski, Tel Aviv University, Israel |  |
| Mouse Cell line: DBT-Luc | Lab of Dr. Dinesh Thotala, Washington University in St. Louis St Louis, Missouri, United States |  |
| Human Cell line: ST1 (HPMECs) | Lab of Dr. Ron Unger, Johannes Gutenberg University, Germany. |  |
| Human brain-derived microvascular endothelial cells (HBMECs) | Cell Biologics, USA | #H-6023 |
| B.End3 Mouse brain-derived immortalized endothelial cells | The American Type Culture Collection | #CRL-2299 |
| **Experimental models: organisms/strains** |  |  |
| Mouse: C57BL/6J, females, 6-8 weeks old | The Jackson Laboratory | JAX:000664 |
| Mouse: Athymic nude, Nude, 6-8 weeks old, females | The Jackson Laboratory | JAX 002019 |
| Oligonucleotides |  |  |
| All shRNA sequences used in the study are provided in the methods section. |  |  |
| Primers for gene expression, detail given in the methods section |  |  |
| Recombinant DNA | Gene Tools |  |
| Plasmid: RAR3G (TetOn) | Lab of Dr. Dinorah Friedmann-Morvinski |  |
| Plasmid: FMOD overexpression construct in pcMV-entry backbone | Origene | #LY419579 |
| Plasmid: CSL Luc | Lab of Prof. Thomas Kandesch, Department of Genetics, University of Pennsylvania School of Medicine. |  |
| Plasmid: pHes1(2.5k)-luc | Addgene | Cat # 43806 |
| NICD pCMV Neo/intracellular domain of human Notch1 (NIC-1) | Lab of Prof. Thomas Kandesch, Department of Genetics, University of Pennsylvania School of Medicine. |  |
| Software and algorithms |  |  |
| ImageJ | Chen et al., 2013 |  |
| MaxQuant software package | Cox et al., 2008 |  |
| Zen Black from Zeiss | https://www.zeiss.com/microscopy/int/products/microscope-software/zen.html |  |
| Other |  |  |
| Reverse Phase Protein Array (RPPA) analysis | https://www.mdanderson.org/research/research-resources/core-facilities/functional-proteomics-rppa-core.html | M.D. Anderson Cancer Center, University of Texas, USA. |

#### Neurosphere culturing

The GSCs are obtained by dissecting GBM tumor tissue and then treating with Tryspin, followed by Trypsin inhibitor. The chopped tissue is then passed through a cell strainer to remove the debris and the obtained filtrate is plated in ultra-low attachment plates using the stem cell for neurosphere formation media described in the following sentence. Then neurospheres were grown in Neurobasal medium (#21103049, Gibco) supplemented with 1X L-Glutamine (# 25030081, stock 200 mM i.e.,100X, Gibco), Heparin 2μg/ml (#H3393 Sigma), 1X B27 supplement (#17504044, Gibco, stock concentration 50X), 0.5X N2 supplement (#17502048, Gibco, stock concentration 100X), 20ng/ml rhEGF (#g5021, Promega), 2ng/ml rhFGF-basic (#100-18B-100 μg, Peprotech) and Penicillin and Streptomycin. To make single-cell suspensions for re-plating, the spheres are chemically dissociated after 7 days of plating, using NeuroCult™ Chemical Dissociation Kit (Mouse, #05707) from Stem Cell Technologies according to the manufacturer’s instructions.

#### Differentiation of GSCs to DGCs

For differentiation of GSCs to DGCs, fully grown neurospheres were collected from the ultra-low attachment plates (Corning, USA) and plated on normal cell culture dishes in DMEM media supplemented with 10% FBS and antibiotics as mentioned earlier, for 15-20 days (***Suva et al., 2014***).

#### Reprogramming of DGCs

The differentiated counterparts of GSCs as well as differentiated GBM cell lines were removed from 10% FBS containing DMEM, spun down twice in PBS to remove any trace of FBS and plated on ultra-low attachment plates in Neurobasal medium containing all the supplements mentioned earlier and antibiotics for 7-10 days.

#### Neurosphere assay

Viral infection was done in the GSCs using lentivirus for non-targeting shRNA (shNT) or shRNA for the gene of interest. The small pellets of cells were collected 48 hours after viral infection, dissociated to form single cells that were counted and re-plated in 6-well plates (30,000 cells/well) in complete Neurobasal medium. At the same time, cells were harvested and checked for specific gene manipulation (like knockdown verification). Media was replenished every 2-3 days and sphere formation was monitored till the 6^th^ or 7^th^ day after re-plating. The number of spheres, sphere diameter and size were analyzed using Image J software. Spheres having area less than 50 µm^2^ were excluded from the analysis.

#### Limiting Dilution assay

For each condition, 1, 10, 50, 100 and 200 GSCs (single cells) were plated in 10 wells each, respectively, of a 96-well plate and sphere formation was assessed over the next 5-7days. The number of wells not forming spheres were counted and plotted against the number of cells per well. Extreme Limiting Dilution Assay using the ELDA software available online (https://www.elda.at/cdscontt/?contentid=10007.854970&portal=elaportal).

#### CM collection, concentration and sample preparation for mass spectrometry

GSC cell lines growing as neurospheres were grown in complete Neural Stem Cell medium containing Glutamine, Heparin, N2, B27, EGF and FGF for 6 days. They were thoroughly washed using PBS and re-plated without disrupting them in Neural Stem Cell medium devoid of supplements and growth factors. Thirty-six hours after plating in incomplete medium, the conditioned medium was collected, spun at 1,500 rpm for 15 min, filtered using 0.45 micron syringe filters and stored at −80°C. The DGCs were grown in complete DMEM supplemented with 10% FBS, then washed with PBS and CM was collected after 36 hours in incomplete medium and similarly stored.

The conditioned media were concentrated using Centricons (3 kDa cut off; Merck) from 8 ml to 100 μl, followed by precipitation of proteins with trichloroacetic acid (TCA 10% for 30 min at 4°C). Equal amounts of proteins from each condition were run on gradient (4-20 %) SDS gels and stained with Colloidal Coomassie blue. The gels were then destained and each lane cut into 5 equal pieces. Proteins were digested in-gel using trypsin (Gold Promega, 1 μg per band, overnight at 30°C,) as previously described (***Thouvenot et al., 2008***).

#### Mass Spectrometry, protein identification and relative quantification

Trypsic peptides were analysed online by nano-flow liquid chromatography coupled to tandem-mass spectrometry using a Q-Exactive+ mass spectrometer (Thermo Fisher Scientific, Waltham USA) coupled to a RSLC-U3000 HPLC (Thermo Fisher Scientific). Desalting and pre-concentration of samples were performed on-line on a Pepmap^®^ pre-column (0.3 mm × 10 mm, Dionex). A gradient consisting of 2–25% B for 80 min, 25-40 % B for 20 min and finally 40-90% B for 5 min (A = 0.1% formic acid; B = 0.1% formic acid in 80% acetonitrile) at 300 nL/min was used to elute peptides from the capillary reverse-phase column (0.075 mm × 150 mm, Acclaim Pepmap 100^®^ C18, Thermo Fisher Scientific). Eluted peptides were electro-sprayed online at a voltage of 1.5 kV into the Q-Exactive+ mass spectrometer. A cycle of one full-scan mass spectrum (MS1, 375–1,500 m/z) at a resolution of 70,000, followed by 12 data-dependent tandem-mass (MS2) spectra was repeated continuously throughout the nano-LC separation. Parameters used for MS2 spectra were: resolution of 17,500, AGC target of 1e5, a normalized collision energy of 28 and an isolation window of 1.2 m/z. Mass spectra were processed using the MaxQuant software package (v 1.5.5.1, ***Cox and Mann, 2008***) against the UniProtKB Reference proteome UP000005640 database for Homo sapiens (release 2018_11) and contaminant database. The following parameters were used: enzyme specificity set as Trypsin/P with a maximum of two missed cleavages, Oxidation (M) and Acetyl (Protein N-term) set as variable modifications and carbamidomethyl (C) as fixed modification, and a mass tolerance of 0.5 Da for fragment ions. The maximum false peptide and protein discovery rate was specified as 0.01. Match between runs was used to transfer identification from run to run. Relative protein quantification was performed using LFQ intensity in MaxQuant. For statistical analysis, missing values were defined using the imputation tool of the Perseus software (v. 1.6.1.1, ***Tyanova et al., 2016***).

#### Lentivirus preparation and transduction of cells

HEK-293T cells were seeded in 60 mm or 90 mm Poly-L-Lysine coated cell culture dishes. The cells were transfected with shRNA plasmid and helper plasmids psPAX2 and pMD2.G using Opti-MEM (Invitrogen #22600-050) medium and lipofectamine (https://www.thermofis.com/order/catalog/product/12566014Invitrogen), when the cells were 60-70% confluent. Six hours after transfection, the Opti-MEM medium was replaced by fresh DMEM medium supplemented with 10% FBS. Sixty hours post-transfection, the supernatant from the transfected cells were collected in 15 ml falcon tubes, centrifuged at 5000 rpm for 10 min, filtered through 0.45 μm, and stored in −80°C for future use.

For virus used in endothelial cells, DMEM was removed after 24 hours of transfection and changed to complete M199 and the virus was collected after sixty hours as mentioned previously. The same method was used for virus to be used for GSCs (DMEM was changed to NBM).The shRNA construct number TRCN0000152163 from Sigma human TRC shRNA library was used for knockdown studies of human FMOD. The shRNA construct number TRCN000094248 from Sigma mouse TRC shRNA library was used for knockdown studies of mouse FMOD. A pooled lentivirus using the construct numbers TRCN000024441, TRCN000024441, TRCN000024441, and TRCN000024420 were used for knock-down studies of JAG1 in the ST1 cells.

#### Endothelial cell culture

ST1 cells were grown in Medium 199 (Sigma #M4530) supplemented with 20% FBS and ECGS (Sigma #E2759), Heparin, Glutamine (1X) and 1X Antibiotic-Antimycotic Solution (Sigma, # A5955, stock concentration 100X).

#### Transdifferentiation of DGCs

The MGG8-DGCs were grown in Medium 199 having the same composition as mentioned above, for three days and subjected to hypoxia (1% O_2_) for 8 hours. The cells then formed transdifferentiated endothelial cells (TDECs), showed endothelial morphology (data not shown), were harvested for checking markers levels, and were also plated for the *in vitro* angiogenesis assay.

#### NICD stable generation

For NICD stable cell-line generation, the ST1 cells were transfected with NICD pCMV Neo/intracellular domain of human Notch1 (NIC-1) plasmid using Lipofectamine 200 (Invitrogen), and then were harvested 48 hours after transfection for western blot and *in vitro* angiogenesis assay.

#### RNA isolation, cDNA conversion and Real-Time Quantitative RT-PCR analysis

Total RNA from the cells was isolated from cells using the TRI reagent (#T9424 Sigma) according to the manufacturer’s instructions. Next, the integrity of the RNA was checked by running it on 2% MOPS-formaldehyde gel. RNAs were quantified by Nanodrop. Two µg of total RNA were used for cDNA conversion using the High capacity cDNA reverse transcription kit (Applied Biosystems, USA) according to the manufacturer’s protocol. The cDNAs were diluted in a ratio of 1:10 with nuclease-free water to make its final concentration 10 ng/µl. Subsequently, real-time quantitative PCR was done using the ABI Quant Studio 5 and 6 (Life technologies, USA). The cDNA was used as template and DyNAmo Flash SYBR Green qPCR kit (#F-416L) was used. Gene-specific primer sets were used for the reaction (Table at the end of the Methods section0) under the following conditions: 95°C for 15 min, 40 cycles of 95°C for 20 sec, 60°C for 25 sec and 72°C for 30 sec followed by the dissociation cycle for melt curve generation. Each sample was run either in duplicate or triplicate. GAPDH (Glyceraldehyde 3-phosphate dehydrogenase), ACTB (beta actin), 18S rRNA, RPL35a (ribosomal protein L35a) and *ATP5G1* [ATP synthase, H+ transporting, mitochondrial F0 complex, subunit C1 (subunit 9)] were used as reference genes for human gene expression analysis. For mouse gene expression analysis, Cyclophilin was used as a housekeeping gene. ΔΔCT method was used for the calculation of gene expression, which was transformed to log2 ratio and then to absolute scale for plotting.

#### Western Blotting

For Western blot analysis, cell pellets were lysed in RIPA lysis buffer (containing 1 mM sodium orthovanadate, 5mM sodium fluoride, 1 mM phenylmethanesulfonyl fluoride and 1X protease inhibitor cocktail, Sigma, USA), and proteins were isolated from the cells by spinning at 13,000 rpm for 30 mins. The supernatant, containing the proteins was collected. Protein concentrations were measured using the Bradford’s reagent and a standard BSA curve was used to determine the protein concentrations. Equal amounts of proteins from all conditions were mixed with protein loading dye (1X), denatured at 95°C for 15 mins, loaded in each well of an SDS polyacrylamide gel and the gel was run for around 8 hours. For preparation of SDS-polyacrylamide gel, resolving and stacking gels were prepared at a concentration of 10-12 concentrations.. The gel was run at 70V-100V and then the proteins were transferred onto a polyvinyl fluoride (PVDF) membrane using the semi-dry transfer method. After the transfer, the membrane was blocked using 5% skimmed milk in 1X Tris-buffered saline-Tween (TBST) for 1 h. Subsequently, the membrane was washed in TBST for 30 min and probed initially with primary antibodies in 5% BSA-TBST for 14-16 hours at 4°C. Then the membrane was washed in TBST for 30 min and secondary antibody, diluted in 5% skimmed milk in TBST was added and incubated at room temperature for 2-3 h. Finally, the blot was washed and developed using Perkin-Elmer ECL Plus lightning and Biorad Clarity and Clarity Plus ECL chemiluminescent reagent using GE Image Quant machine.

#### Chromatin immunoprecipitation (ChIP)

ChIP assay was conducted with chromatin isolated from ST1 cells treated with 10 μM TGFβ RI inhibitor (SB431542) and DMSO for 6 hrs. Briefly, after cross-linking, the nuclei were prepared and sonicated to generate chromatin fragments between 100 and 10,000 bp following the manufacturer’s protocol using Simple Chromatin Immunoprecipitation kit (CST; Cat no. 9003). The sheared chromatin was collected by centrifugation (10000g for 10 min at 4°C) and a 10 μL aliquot was removed to serve as a positive input sample. Aliquots of 100 μL sheared chromatin were incubated with 2 μg of the required antibody/antibodies followed by Protein G magnetic beads for the stipulated time. An equal amount of IgG and H3 antibodies were used as controls. The eluted DNA was analyzed by quantitative PCR using the FMOD promoter-specific primer set ChIP-F/ChIP-R (in list of primers) to amplify the desired region in FMOD promoter. Conditions of linear amplification were determined empirically for this primer. The PCR conditions were as follows: 95°C for 5min; 95°C for 30 sec, 56°C for 30 s and 72°C for 30 s for 35 cycles. PCR products were resolved by electrophoresis on a 2% agarose gel and visualized after ethidium bromide staining. Real time qPCR was performed with the same eluted DNA. The conditions were as follows: 95°C for 3 min; 95°C for 10 sec, 56°C for 30 sec, 72°C for 30 s for 40 cycles and 72°C for 5 mins. The Ct values of different conditions were normalized to Ct values in IgG control.

#### Boyden chamber assay for cell migration

Trans-well assay was done in 24-well Boyden chambers with 8-μm pore size polycarbonate membranes (BD Biosciences, San Diego, USA). ST1 cells (5 × 10^4^) were re-suspended in 500 μl serum-free Medium 199 and placed in the upper chamber, and the lower chamber was filled with serum-free Medium 199 with 750 μl conditioned medium or 400nm final concentration of recombinant protein dissolved in incomplete medium (serving as a chemo-attractant). After 24 hours of incubation, cells remaining on the upper surface of the membrane were removed with a wet cotton bud. The cells that have migrated to the lower surface of the membrane were fixed in ice-cold methanol and stained with crystal violet and imaged in a light microscope.

#### Boyden chamber assay for cell invasion

Trans-well assay was done in 24-well Boyden chambers with 8-μm pore size polycarbonate membranes coated with Matrigel (BD Biosciences, San Diego, USA). ST1 cells (5 × 10^4^) were re-suspended in 500 μl serum-free Medium 199 and placed in the upper chamber, and the lower chamber was filled with serum-free Medium 199 with 750 μl conditioned medium or 400nm final concentration of recombinant protein dissolved in an incomplete medium (serving as a chemo-attractant). After 24 hours of incubation, cells remaining on the upper surface of the membrane were removed with a wet cotton bud. The cells that have invaded to the lower surface of the membrane were fixed in ice-cold methanol and stained with crystal violet and imaged in a light microscope.

#### Cell proliferation assay (MTT assay)

1.5 x 10^3^ ST1 cells were plated 2% FBS-containing M199 Medium in each well of a 96-well plate. MTT assay was performed as per the established protocol. MTT was added to each well and Formazan crystals formed after 3 hours of incubation were dissolved in DMSO and the absorbance was measured at 420 nm. The first reading served as the untreated condition (0^th^ time-point). After this reading, the cells were treated with 400nm rhFMOD or an equivalent concentration of BSA, and readings were taken every 24 hrs, till 96 hours. The cell viability was then plotted as a line graph.

#### *In vitro* angiogenesis assay

ST1 endothelial cells were seeded in 96-well plates (10,000-15,000 cells per well), coated with Geltrex (Invitrogen) and grown in Medium 199 (Sigma) without growth factors. Equal protein amounts (50-100 µg) from serum-free conditioned media from different conditions were added on top of the cells. After 10-12 hours of incubation, endothelial cells form tube-like structures. Each complete circular structure was considered as one complete network and the total number of networks for each condition was counted in a double-blind manner. For positive control, cells are plated in complete endothelial cell media (Medium 199) supplemented with Endothelial Cell Growth Factors (ECGS) and 20% FBS, and in the negative control, cells are plated in incomplete Medium 199 (without serum and ECGS).

#### Immunofluorescence staining of fixed cells

Cells were plated on coverslips in 12-well plates and allowed to attach. The cells were then fixed with 4% paraformaldehyde and permeabilized using PBS supplemented with 0.25% Triton-X100. Cells were then washed with PBS and blocked using PBS supplemented with 1% BSA, 0.3% Triton X-100 and 5% goat serum for 2 h at room temperature. After blocking, primary antibody, diluted in the blocking buffer, was added to the coverslips overnight at 4°C. Foor dual staining, the primary antibodies (one anti-mouse and the other anti-rabbit) were together added to the samples in the required dilutions. After removal of the primary antibody, cells were thoroughly washed with PBS 3 times for 5 mins each. Fluorescence-conjugated secondary antibody was dissolved in the blocking buffer and added to the cells for 3 h, after which the cells were washed three times with PBS for 5 min. The cells were stained with DAPI (1 µl/ml) for 5 min at room temperature. Coverslips were mounted on glass slides using glycerol as mounting agent and imaged using a Zeiss LSM 880 confocal microscope.

#### Luciferase reporter assay

Cells were seeded in a 6-well plates and co-transfected with the reporter luciferase construct and pCMV-beta gal (as control), using lipofectamine according to the manufacturer’s instructions. 24-48 h after transfection, cell extracts were made in reporter lysis buffer (Promega). Protein concentrations of the cell lysates were measured by Bradford assay reagent (Bio-Rad). Ten µg protein were mixed with 30 µl of luciferase assay reagent (LAR) to determine the luciferase activity and values were normalized to beta-galactosidase activity units.

#### Flow cytometry

The GSCs and DGCs are pelleted down and stained with live dead fluorescent stain (L34955 LIVE/DEAD™ Fixable Violet Dead Cell Stain Kit, for 405 nm excitation) for 15 mins at 37 °C. Then, the cells are blocked using 10% FCS and 1% Sodium Azide in PBS. After blocking, the cells are washed with PBS thrice, incubated with primary antibody for 2 hrs at room temperature, washed again thrice, and incubated with the secondary antibody for 2 hrs in the dark. The final pellet is washed thrice with PBS, dissolved in 200 μl 3% BSA in PBS and analysed using the BD FACS Verse flow cytometer.

#### RPPA analysis

Untreated ST1 cells and cells treated for 10, 30, and 60 mins with 400 nm rhFMOD were lysed using RPPA lysis buffer. Lysates were serially diluted in 5 two-fold dilutions using lysis buffer and printed on nitrocellulose-coated slides using an Aushon Biosystem 2470 arrayer. Slides were probed with 304 validated primary antibodies followed by detection with appropriate biotinylated secondary antibodies. Slides were scanned, analyzed, and quantified using Array-pro Analyzer software (Media Cybernetics) to generate spot intensity (level 1 data). Signals were visualized by a secondary streptavidin-conjugated HRP antibody and DAB colorimetric reaction. The list of 304 antibodies can be found at the link provided (http://www.mdanderson.org/research/research-resources/core-facilities/functional-proteomics-rppa-core/antibody-information-and-protocol.html).

#### Cryo-sectioning of fixed mouse brain

Mice were perfused intracardiacally using 4% paraformaldehyde solution and the brains were harvested and stored in PFA for 12 h and subsequently in 30% sucrose solution. The brains were then embedded in Poly-freeze solution (Sigma, #35059990) and sectioned into 20-μm thick sections using a Leica Cryostat. The sections were stored in −80°C in Tissue Cutting Solution (TCS).

#### IHC of free-floating sections

For immunofluorescence, the brain sections were removed from TCS, put in 96-well plates and washed thoroughly with PBS. Following that, the same protocol was followed for immunofluorescence staining of tissue sections, as that followed for monolayer cells grown on coverslips. At the final step, after adding the secondary antibody, DAPI and PBS washes, the sections were individually mounted on glass slides using the ProLong™ Glass Antifade Mountant (Invitrogen, # <u>P36980</u>) and covered by coverslips. Images were taken using the Zeiss LSM 880 confocal microscope using 10x, 20x, and 40x objectives, for various conditions.

For immunofluorescence of FFPE sections, an extra step of antigen retrieval was performed by de-paraffinizing the sections in xylene, followed by boiling in distilled water twice, for 5 mins each. After this, the rest of the steps (from permeabilization to mounting, the same steps were followed as that of immunofluorescence of free-floating sections.

#### Scoring methods for confocal images

The areas of the blood vessels and the fluorescence intensities for all the fluorophores used in the tissue sections were measured using the Zeiss black software (https://www.zeiss.com/microscopy/int/products/microscopesoftware.html), converted to percentages and plotted as bar diagrams. For determining the extent of vascular mimicry in the blood vessels of the tissues, the overlap of green (coming from GFP of the tumor cells) and red (measure of CD31 expression of the endothelial cells) forming yellow color was calculated as a measure of co-localization coefficients in the Zeiss black software. The absolute values of the co-localization coefficients were plotted for each condition.

#### Subcutaneous injection of DBT-Luc cells for the co-implantation mouse model

All animal procedures followed were approved by the Institute Ethical Committee for Animal Experimentation. Mouse cell line DBT-Luc-DGC, which grows as a monolayer cell line (stable for luciferase expression), were reprogrammed to form DBT-Luc-GSCs. A combination of 10^5^ DBT-Luc-GSCs (without any shRNA) and 10^6^ DBT-Luc-DGCs that carried either the miRNT or miRFMOD constructs (referred to as DBT-Luc-DGC/miRNT or DBT-Luc-DGC/miRFMOD respectively) were injected subcutaneously in the mice. Two groups of mice (n=5 each) received a combination of 10^5^ DBT-Luc-GSCs + 10^6^ DBT-Luc-DGC/miRNT and another two groups (n=5 each) received a combination of 10^5^ DBT-Luc/GSCs + 10^6^ DBT-Luc-DGC/miRFMOD (only one of the two groups for both miRNT and miRFMOD received Doxycycline). Doxycycline injection (intraperitoneal, 100 μg per animal) began on 9^th^ day post the injection and was given on every day up to the 16^th^ day, after which it was given on every alternate day, till the end of the experiment. Since the Doxycycline administration induced the mCherry expression along with the shRNA expression, the animals were imaged for both bioluminescence (marked by red on the timeline) and fluorescence (marked by orange on the timeline) on the days indicated in the timeline. Two other groups (n=5 each) received either only 10^5^ DBT-Luc-GSCs or 10^6^ DBT-Luc-DGCs (not stables for any shRNA), as controls.

#### Intra-cranial injection of GBM cells

Cells were harvested in incomplete DMEM or NBM depending on the cell type to be injected. 250,000 DGCs or 100,000 GSCs were injected intracranially (in the hippocampus, 3 mm deep) in each animal using a stereotaxic apparatus. The animals were imaged on the 3^rd^ day after injection and subsequently, every 5 to 6 days until the end of the experiment. In experiments using inducible shRNAs, doxycycline was injected every day for 10 days (from 13^th^ day after injection) and on every alternate day until the end of the experiment. MGG8-GSC (MGG8-GSC/shNT vs. MGG8-GSC/shFMOD), AGR53-GSCs (AGR53-GSC/miRNT vs. AGR53-GSC/miRFMOD), and DBT-Luc-GSCs (DBT-Luc-GSC/miRFMOD) cells were intracranially injected in this study.

#### In vivo imaging

*In vivo* imaging was done for bioluminescence or fluorescence with the Perkin Elmer IVIS Spectrum by using mild gas anesthesia (using isoflurane) for the animals.

#### Hematoxylin and Eosin staining

Brains from perfused mice were paraffin embedded and sectioned using a microtome (5μ sections). Sections were mounted on glass slides and removed from paraffin, rehydrated and stained with Harris Hematoxylin for nuclear staining and Eosin Y solution for cytoplasmic staining. The sections were then mounted using DPX mounting medium and imaged at 0.8X using a Lawrence and Mayo digital microscope.

#### Gene Set Enrichment Analysis (GSEA)

The differentially expressed genes between the GSC and DGC (as identified in GSE54792) were pre-ranked based on fold change and used as an input to perform GSEA. All the gene sets available in the Molecular Signature Database (MSigDB, roughly 18,000 gene sets) were used to run the GSEA. We filtered out the TGF-beta pathway related gene sets to identify that most of them were significantly enriched in the DGCs over the GSCs. Similarly, the same analysis was carried out in multiple publicly available GBM vs. normal samples datasets to show the significant enrichment of TGF-beta gene sets in GBM over normal. We acknowledge our use of the GSEA software and MSigDB (Subramanian et al., 2005) (http://www.broad.mit.edu/gsea/). Single

#### Single Sample GSEA (ssGSEA)

Gene set variation analysis (gsva) was performed using ssGSEA to determine the enrichment of the different molecular subtypes of GBM in GSE54792 and also the enrichment of the TGF-beta Hallmark gene set from MSigDb. The higher gsva score indicates the highest enrichment which gradually decreases.

#### Survival analysis

Kaplan-Meier survival analysis was done using GraphPad Prism 5.0 (GraphPad Software, San Diego, California, USA).

#### Heatmap generation

The heatmaps were generated using the Multiple Experiment Viewer (MEV) software (http://www.tm4.org/mev.html) version 4.8.1. LFQ values for protein expression from mass-spec data, mean pixel density for the RPPA were used as inputs for Heatmap generation. A non-parametric t-test was performed with a false discovery rate (FDR) and a p-value cut-off of 0.05.

#### Quantification and Statistical analysis

Bar diagrams are generated using Microsoft excel. The box plots are generated using GraphPad Prism 5.0 (GraphPad Software, San Diego, California, USA). p-value is calculated by unpaired t test with Welch’s correction are indicated or student t-test was done using Microsoft Excel. ANOVA p value was calculated using GraphPad Prism 5 software. p value less than 0.05 is considered significant with *, **, *** representing p value less than 0.05, 0.01 and 0.001 respectively. ns stands for non-significant.

#### List of primer used in this study

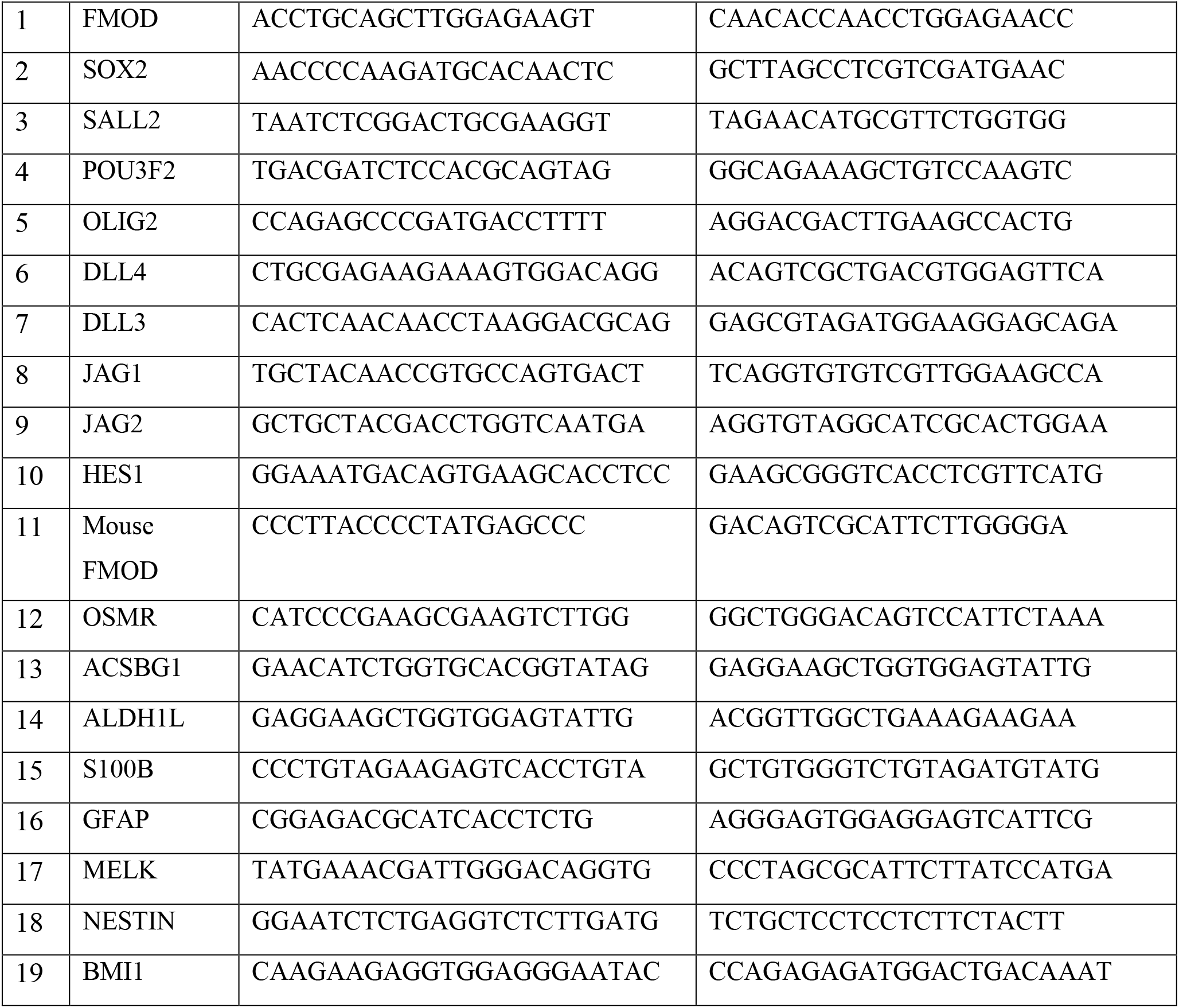
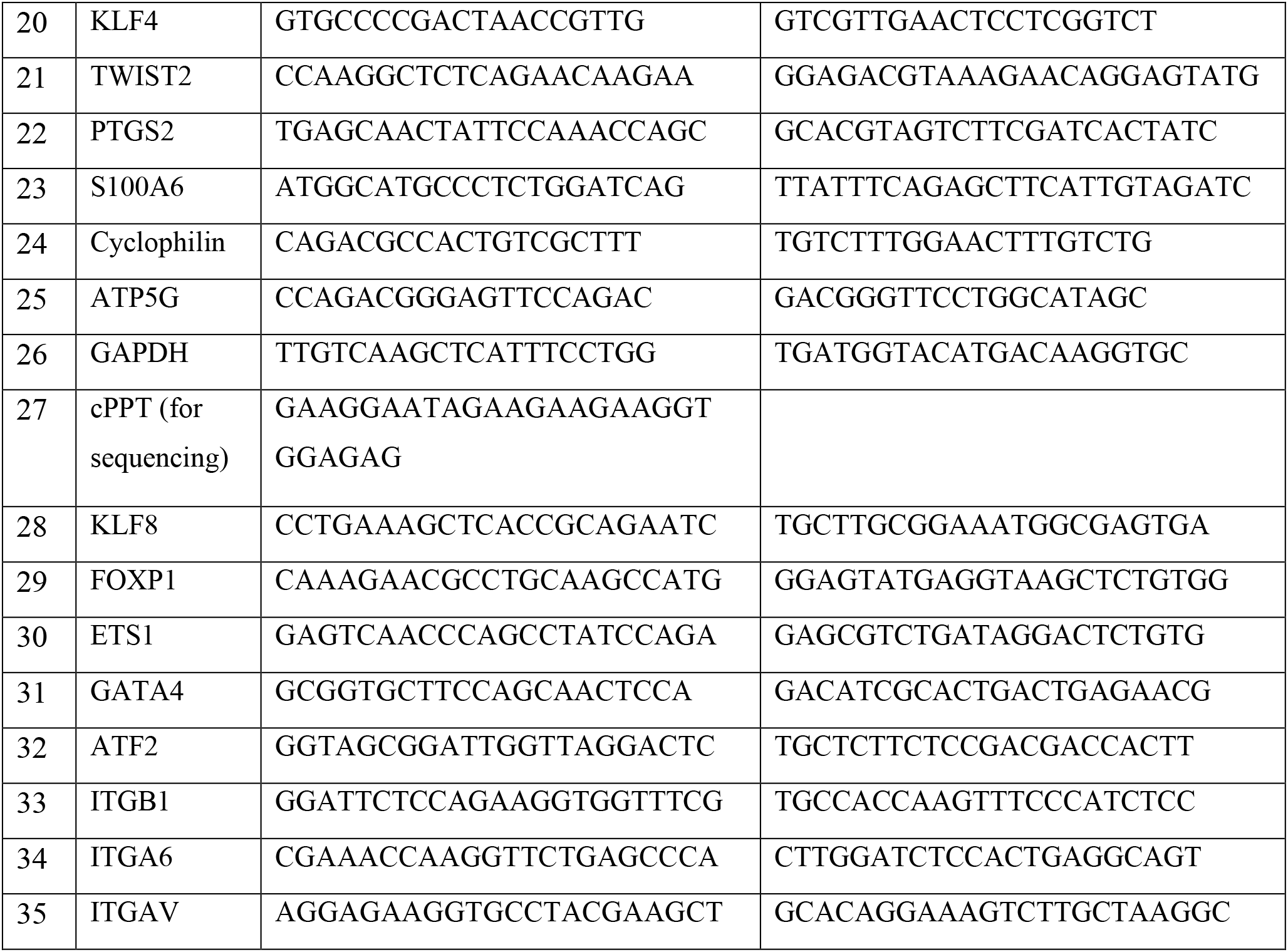

#### List of shRNAs used in the study (all IDs are of Sigma TRC whole-genome shRNA library)

| Sl.no | Gene Name | Clone 1 | Clone 2 | Clone 3 | Clone 4 | Clone 5 |
| --- | --- | --- | --- | --- | --- | --- |
| 16 | FMOD | TRCN0000153650 | TRCN0000156734 | TRCN0000152163 | TRCN0000153199 | TRCN0000151908 |
| 17 | Mouse FMOD | TRCN0000094246 | TRCN0000094245 | TRCN0000094248 |  |  |
| 18 | KLF8 | TRCN0000015878 | TRCN0000015879 | TRCN0000015880 | TRCN0000015881 | TRCN0000015882 |
| 19 | JAG1 | TRCN0000033439 | TRCN0000033440 | TRCN0000033441 | TRCN0000033442 | TRCN0000033443 |
| 20 | ITGA6 | TRCN0000057773 | TRCN0000057774 | TRCN0000057775 | TRCN0000057776 | TRCN0000057777 |
| 21 | ITGB1 | TRCN0000275134 | TRCN0000275133 | TRCN0000275083 | TRCN0000275135 | TRCN0000275082 |
| 22 | ITGAV | TRCN0000003238 | TRCN0000003239 | TRCN0000003240 | TRCN0000003241 |  |

#### List of inhibitors used

**List of inhibitors used:**
| S.No. | Inhibitor Name | Target Molecule | Catalog Number |
| --- | --- | --- | --- |
| 1 | Gamma- secretase inhibitor (GSI) | Gamma Secretase | 565750 (Merck) |
| 2 | RGD Peptide | Integrins | A5082 (Sigma Aldrich) |
| 3 | PP2 | Src | P0042 (Sigma Aldrich) |
| 4 | PP3 | Structural analog to PP2 | 529574 (Calbiochem) |
| 5 | H1152 | ROCK1 | 555550 (Calbiochem) |
| 6 | RAC1 inhibitor | RAC1 | 553502 (Sigma Aldrich) |
| 7 | PF573228 | FAK | PZ0117 (Sigma Aldrich) |
| 8 | SB431542 | TGFB RI | 616464 (Sigma Aldrich) |

## Supplemental item titles

1. Supplementary information
2. Supplementary figure legends
3. Supplementary Figures 1-28
4. Supplementary Table S1
5. Supplementary Table S2
6. Supplementary Table S3

## Supplementary information

### The role of FMOD on GSC, DGC growth, and their plasticity *in vitro*

Before studying the importance of DGC secreted FMOD on tumor growth, we decided to investigate its requirement for the GSC, DGC growth, and their plasticity to form the other type. We have earlier shown that FMOD is not required for the proliferation of established glioma cell lines **(Mondal et al., 2017).** We used two human glioma cell lines-MGG8 and U251, two murine glioma cell lines-AGR53 (***Angel et al., 2020***), and DBT-Luc (**Yun et al., 2007**).

MGG8 GSCs transduced with a small hairpin RNA targeting FMOD (MGG8 GSC/shFMOD) grew as neurospheres with equal efficiency as measured by neurosphere formation and limiting dilution assays and also differentiated to form DGCs as efficiently as control MGG8 GSCs transduced with non-targeting shRNA (MGG8 GSC/shNT) (**Supplementary figure 6A, B, C and D).** As expected, the differentiation was accompanied by the downregulation of glioma reprogramming factors in both MGG8 GSC/shNT and MGG8 GSC/shFMOD cells (**Supplementary figure 6E)**. Concordance with human GSCs, we found a higher expression of FMOD in AGR53-DGCs than AGR53-GSCs. (**Supplementary figures 7A, B, and C**). To silence the expression of FMOD in AGR53-DGC, we used a doxycycline-inducible FMOD shRNA (miRFMOD) construct that contains an inducible mCherry-shRNA cassette downstream of the Tet-responsive element (***Angel et al., 2020***; **Figure 2A**). Efficient silencing of FMOD in doxycycline-treated AGR53-DGC/miRFMOD cells was observed compared to AGR53-DGC/miRNT cells (**Supplementary figure 7D and E).** Next, we investigated the impact of FMOD silencing on AGR53-GSC growth and differentiation to DGCs *in vitro*. Like human GSCs, AGR53-GSC/miRNT and AGR53-GSC/miRFMOD grew as neurospheres with equal efficiency both in the absence and presence of doxycycline (**Supplementary figure 8A).** Further, both AGR53-GSC/miRNT and AGR53-GSC/miRFMOD differentiated efficiently to grow as a monolayer (**Supplementary figure 8C**). The differentiation resulted in the significant upregulation of astrocytic markers (**Supplementary figures 8D**). These results confirm that FMOD is overexpressed in both human and murine DGCs but is not required for GSC growth and their differentiation to DGCs.

To study the role of FMOD on reprogramming of DGCs to form GSCs, AGR53-DGC and DBT-Luc DGC were tested for their ability to reprogram. DBT-Luc is a luciferase-expressing glioblastoma-derived cell line, which grows as a differentiated monolayer (referred here onward as DBT-Luc-DGC) in FBS containing medium (**37**). AGR53-DGC/miRNT and AGR53-DGC/miRFMOD (both in the absence and presence of doxycycline) cells grow as a monolayer efficiently and reprogrammed to form neurospheres (**Supplementary figure 9A**). The reprogramming resulted in the significant upregulation of stem cell markers (**Supplementary figures 9B**). Similarly, DBT-Luc-DGC could readily reprogram to form DBT-Luc neurospheres (DBT-Luc-GSC; data not shown) with concomitant upregulation of stem cell markers and downregulation of astrocyte markers (**Supplementary figure 10A and B**). DBT-Luc-DGCs expressed higher levels of FMOD transcript and protein compared to DBT-Luc-GSCs (**Supplementary figure 10C and D, respectively**). To silence the expression of FMOD in DBT-Luc-DGC, we used the doxycycline-inducible shFMOD (miRFMOD) construct as explained above (***Angel et al., 2020***; **Figure 2A**). The addition of doxycycline resulted in downregulation of FMOD transcript and protein in DBT-Luc-DGC/miRFMOD but not in DBT-Luc-DGC/miRNT cells (**Supplementary figure 10E and F).** As expected, both DBT-Luc-DGC/miRFMOD and DBT-Luc-DGC/miRNT cells showed mCherry expression after doxycycline treatment (**Supplementary figure 10G).** Both DBT-Luc-DGC/miRNT and DBT-Luc-DGC/miRFMOD (both in the absence and presence of doxycycline) cells reprogrammed to form neurospheres with equal efficiency (data not shown). We next explored the impact of FMOD silencing on the ability of U251 cells, an established human glioma cell line, to form neurospheres through reprogramming. U251 cells, which grow as differentiated monolayer cells (referred here as U251-DGC) in an FBS-containing medium, can reprogram and form neurospheres enriched in CD133 expression (***Tao et al., 2018***). Both U251-DGC/shNT and U251-DGC/shFMOD cells could reprogram with equal efficiency, as measured by neurosphere formation and limiting dilution assays (**Supplementary figure 11A, B, and C**). The neurospheres formed through reprogramming showed an upregulation of glioma reprogramming factors in U251-GSC/shNT and U251-GSC/shFMOD cells (**Supplementary figure 11D**). Collectively, these observations indicate that FMOD is not required for GSC or DGC growth and differentiation or reprogramming processes.

### Bioinformatics analysis of FMOD, HES1, and JAG1 transcript levels in GBM– classification, coorelation, and prognosis

The transcript data of FMOD, HES1, and JAG1 in different data sets were used for deriving 1) transcriptional upregulation in GBM over control brain, 2) survival prediction by Kaplan meier analysis, and 3) coorelation between transcripts (**see Table below more detail**). We used a total of 1885 samples that included GBMs (n=1833) and control brain samples (n=52). Whereever, the data is not available (NA), the specific analysis is not carried out. A p value less than 0.05 is considered significant with **** as less than 0.0001, *** as less than 0.001, ** as less than 0.01 and * as less than 0.05. The non significant data is denoted as “ns”. The data is shown in Supplementary Figure 21 and 22.

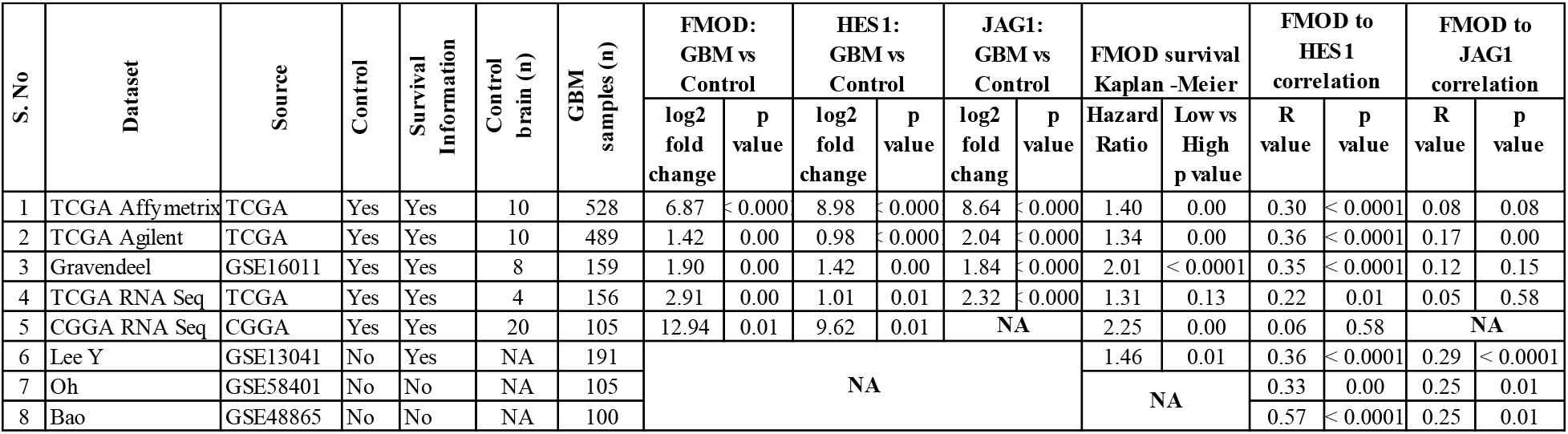

| S.No | Dataset | Source | Control | Survival Information | Control brain (n) | GBM samples (n) | FMOD: GBM vs Control |  | HES1: GBM vs Control |  | JAG1: GBM vs Control |  | FMOD survival Kaplan -Meier |  | FMOD to HES1 correlation |  | FMOD to JAG1 correlation |  |
| --- | --- | --- | --- | --- | --- | --- | --- | --- | --- | --- | --- | --- | --- | --- | --- | --- | --- | --- |
|  |  |  |  |  |  |  | log2 fold change | p value | log2 fold change | p value | log2 fold change | p value | Hazard Ratio | Low vs High p value | R value | p value | R value | p value |
| 1 | TCGA Affymetrix | TCGA | Yes | Yes | 10 | 528 | 6.87 | <0.000 | 8.98 | <0.000 | 8.64 | <0.000 | 1.40 | 0.00 | 0.30 | <0.0001 | 0.08 | 0.08 |
| 2 | TCGA Agilent | TCGA | Yes | Yes | 10 | 489 | 1.42 | 0.00 | 0.98 | <0.000 | 2.04 | <0.000 | 1.34 | 0.00 | 0.36 | <0.0001 | 0.17 | 0.00 |
| 3 | Gravendeel | GSE16011 | Yes | Yes | 8 | 159 | 1.90 | 0.00 | 1.42 | 0.00 | 1.84 | <0.000 | 2.01 | <0.0001 | 0.35 | <0.0001 | 0.12 | 0.15 |
| 4 | TCGA RNA Seq | TCGA | Yes | Yes | 4 | 156 | 2.91 | 0.00 | 1.01 | 0.01 | 2.32 | <0.000 | 1.31 | 0.13 | 0.22 | 0.01 | 0.05 | 0.58 |
| 5 | CGGA RNA Seq | CGGA | Yes | Yes | 20 | 105 | 12.94 | 0.01 | 9.62 | 0.01 | NA |  | 2.25 | 0.00 | 0.06 | 0.58 | NA |  |
| 6 | Lee Y | GSE13041 | No | Yes | NA | 191 | NA |  |  |  |  |  | 1.46 | 0.01 | 0.36 | <0.0001 | 0.29 | <0.0001 |
| 7 | Oh | GSE58401 | No | No | NA | 105 |  |  |  |  |  |  | NA |  | 0.33 | 0.00 | 0.25 | 0.01 |
| 8 | Bao | GSE48865 | No | No | NA | 100 |  |  |  |  |  |  | NA |  | 0.57 | <0.0001 | 0.25 | 0.01 |

For survival prediction based on the methylation status of FMOD promoter, the samples were divided into methylation high (above median β value) and low (below median β value) for two CpG IDs-cg03764585 and cg04704856, derived from TCGA and GSE48461 data sets. The data is presented in Supplementary Figure 22.

### Investigating Integrin heterodimeric subunits that are required in rhFMOD treated endothelial cells

Integrins are a family of α/β heterodimeric transmembrane adhesion receptors. To identify the integrin α and β subunits that are involved in rhFMOD activation of integrin signaling in endothelial cells, we resorted to an unbiased approach where we analyzed the transcriptome data of microvessels isolated through laser capture microdissection (LCM) from human brain samples (**Song et al., 2020, 10: 12358, Scientific Reports**). The transcript abundance of all integrins subunits (α subunits (n=19) and β subunits (n=13) are shown in the table below. Three integrin subunits with maximum transcript abundance – ITGA6, ITGB1, and ITGAV, were chosen for testing. The ability of rhFMOD to induce pFAK levels in ST1 cells silenced for any of the above three integrins was investigated. The results show that all three integrins are essential for rhFMOD activation of integrin signaling in endothelial cells (**Supplementary figure 18**).

| **High abundant integrins in endothelial cells<sup>\$</sup>** | | | | | |
| --- | --- | --- | --- | --- | --- |
| **S. No.** | **Integrins** | **Microvessels 1** | **Microvessels 2** | **Microvessels 3** | **Average** |
|  |  | **FPKM (log2)** |  |  | **FPKM (log2)** |
| 1 | **ITGA6** | 191 | 68 | 249 | 169 |
| 2 | **ITGB1** | 78 | 26 | 151 | 85 |
| 3 | **ITGAV** | 57 | 91 | 61 | 70 |
| 4 | ITGA7 | 23 | 28 | 76 | 42 |
| 5 | ITGA1 | 30 | 4 | 79 | 38 |
| 6 | ITGB8 | 19 | 41 | 15 | 25 |
| 7 | ITGB5 | 39 | 9 | 21 | 23 |
| 8 | ITGA5 | 13 | 11 | 38 | 20 |
| 9 | ITGA10 | 11 | 9 | 40 | 20 |
| 10 | ITGB1BP1 | 16 | 7 | 26 | 16 |
| 11 | ITGA3 | 1 | 15 | 12 | 9 |
| 12 | ITGA2 | 11 | 4 | 8 | 8 |
| 13 | ITGA9 | 0 | 18 | 4 | 7 |
| 14 | ITGB2 | 12 | 6 | 2 | 7 |
| 15 | ITGB4 | 0 | 6 | 13 | 7 |
| 16 | ITGB3BP | 2 | 0 | 16 | 6 |
| 17 | ITGAE | 5 | 1 | 6 | 4 |
| 18 | ITGB7 | 0 | 0 | 11 | 4 |
| 19 | ITGB3 | 6 | 0 | 3 | 3 |
| 20 | ITGA4 | 7 | 0 | 0 | 2 |
| 21 | ITGAL | 1 | 0 | 5 | 2 |
| 22 | ITGA8 | 0 | 1 | 4 | 2 |
| 23 | ITGAX | 4 | 0 | 0 | 1 |
| 24 | ITGAM | 2 | 0 | 0 | 1 |
| 25 | ITGA11 | 1 | 0 | 0 | 0 |
| 26 | ITGB1BP2 | 0 | 0 | 1 | 0 |
| 27 | ITGA2B | 0 | 0 | 0 | 0 |
| 28 | ITGBL1 | 0 | 0 | 0 | 0 |
| 29 | ITGA9-AS1 | 0 | 0 | 0 | 0 |
| 30 | ITGAD | 0 | 0 | 0 | 0 |
| 31 | ITGB2-AS1 | 0 | 0 | 0 | 0 |
| 32 | ITGB6 | 0 | 0 | 0 | 0 |

## Supplementary figure legends

**Supplementary Figure 1:**
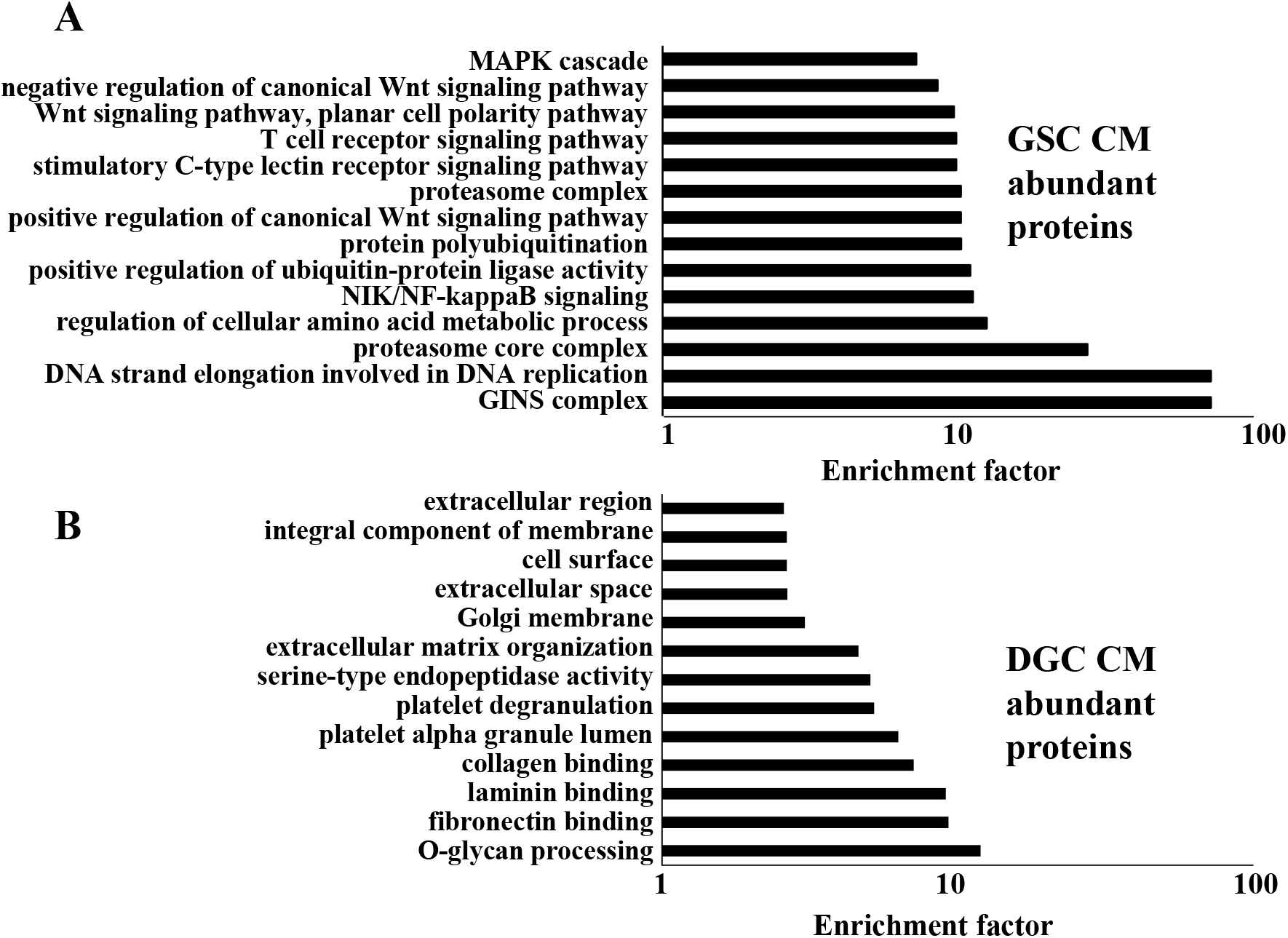
Gene ontology analysis of differentially abundant proteins in GSC and DGC CMs. **A.** Gene Ontology (GO) analysis of proteins exhibiting higher abundance in the GSC CM. **B.** Gene Ontology (GO) analysis of proteins exhibiting higher abundance in DGC CM. Enrichment factor is calculated by-log (q-value). The q-value is a modified Fisher exact p-value provided by the Uniprot Database for annotation, visualization, and integrated discovery enrichment analysis.

**Supplementary Figure 2:**
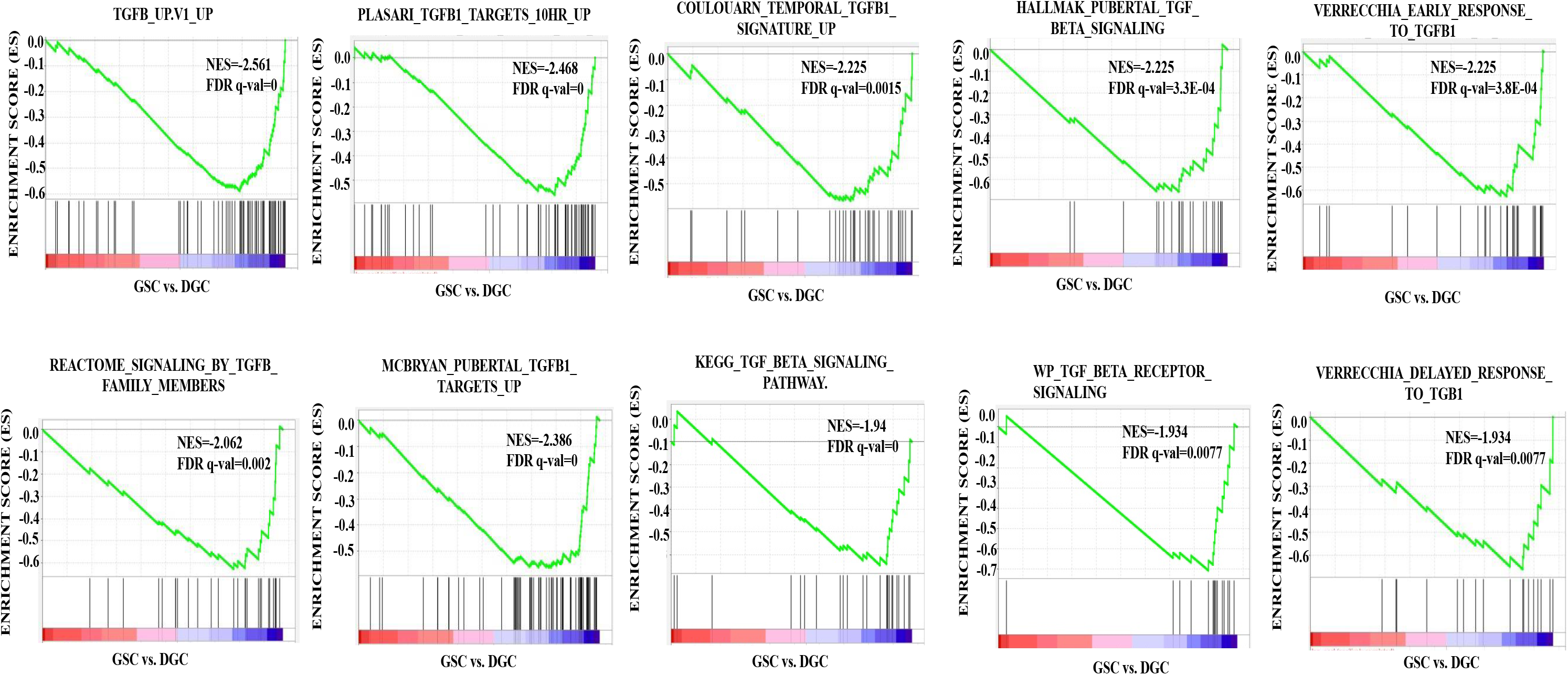
TGF-β pathway is activated in DGCs over GSCs. **A.** Gene Set Enrichment Analysis (GSEA) shows a negative enrichment of multiple TGF-β-related gene sets in GSCs over DGCs, suggesting an activated TGF-β signaling in the DGCs. All these gene sets have positive normalized enrichment score (NES) and significant FDR q-value and p value.

**Supplementary Figure 3:**
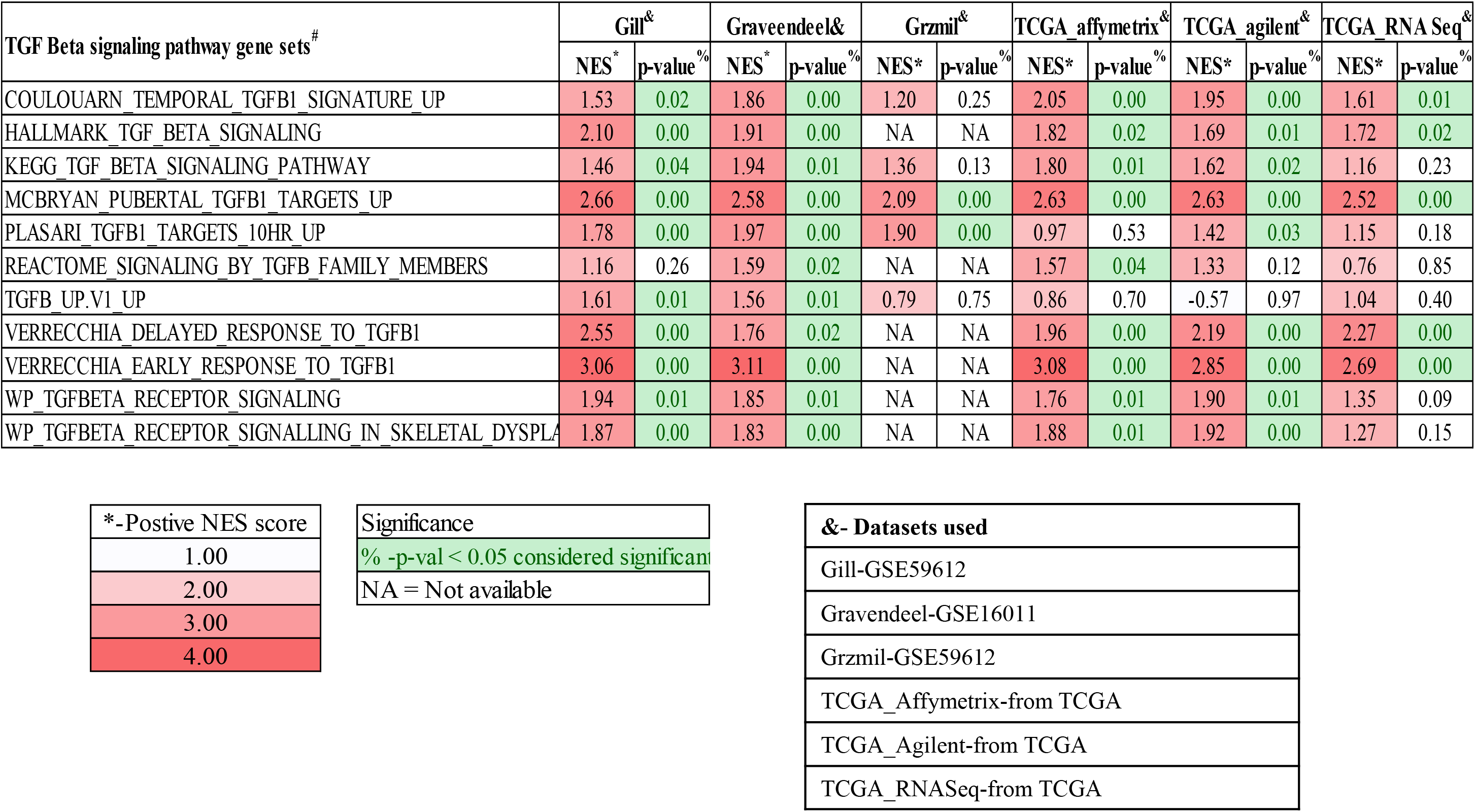
TGF-β is activated in GBM over normal samples in multiple datasets. Gene Set Enrichment Analysis (GSEA) shows the positive enrichment of multiple TGF-β-related gene sets in GBM over normal in multiple publicly available datasets and darker to lighter red indicates highest to lowest normalized enrichment score (NES), while % and green indicates the significant gene sets. p value less than 0.05 is considered significant. NA indicates Not Available.

**Supplementary Figure 4:**
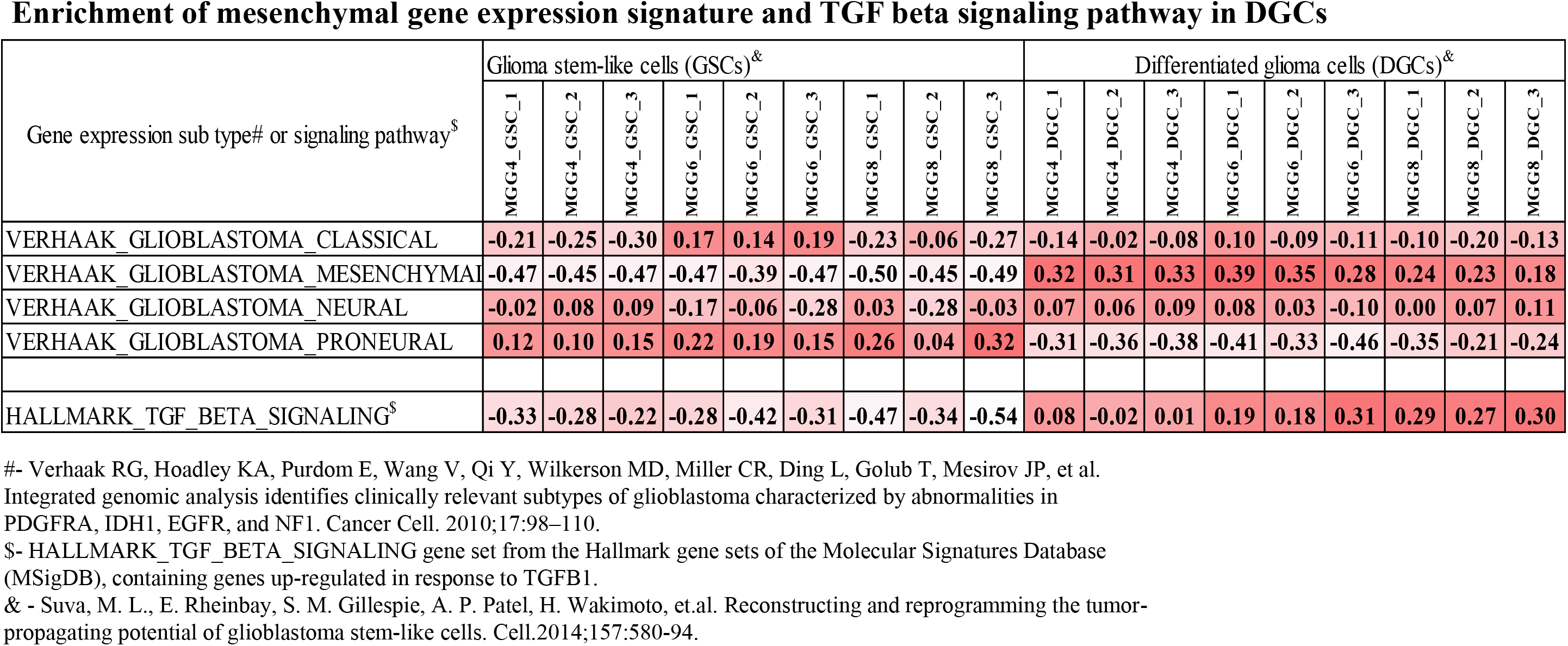
Mesenchymal gene expression signature and TGF-β signaling pathway are enriched in DGCs. Table indicating the subtypes of MGG4, MGG6, and MGG8 GSCs vs. DGCs (each in triplicates). The darkest red indicates the highest value for a subtype that is most enriched in that particular sample, with decreasing color intensity indicating the other lesser enriched subtypes in a gradual manner. The Table also indicates the enrichment of the TGF-β hallmark gene set from MSigDb. The intensity of the red color indicates the varying enrichment scores, with darkest red depicting highest enrichment and gradual lighter colors indicating gradually decreasing enrichment scores.

**Supplementary Figure 5:**
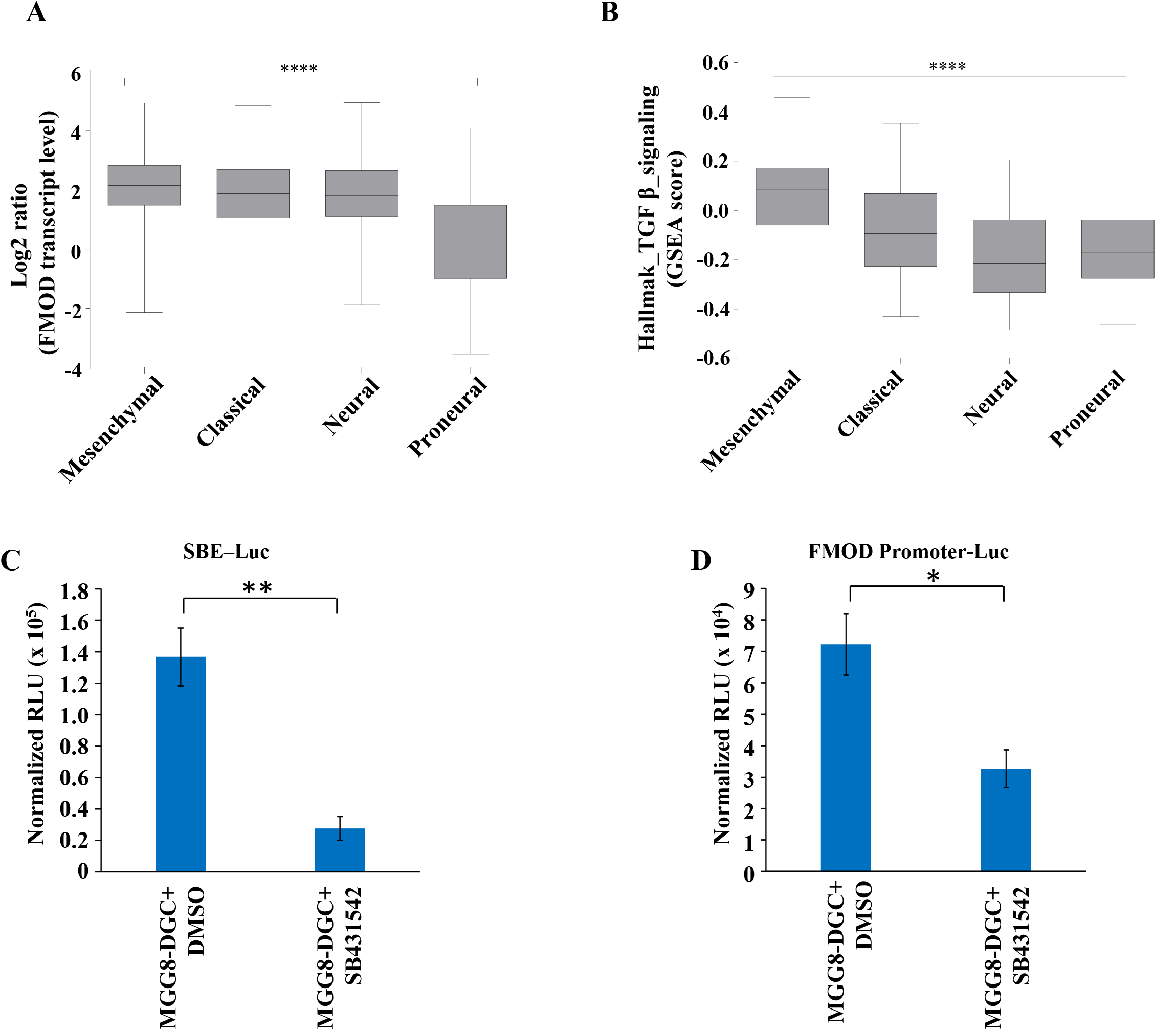
FMOD expression and TGF-β signaling activation are significantly higher in mesenchymal GBM over the other subtypes. **A.** Box plots depicting significantly higher expression of FMOD in mesenchymal GBM samples over the other subtypes in the TCGA Agilent dataset. **B.** Box plots depicting significantly higher enrichment of the TGF-β hallmark gene set from MSigDb in mesenchymal GBM samples over the other subtypes in the TCGA Agilent dataset. **** indicates the ANOVA p value for **A** and **B**. **C.** Bar diagram showing increased SBE-Luc activity indicating activated TGF-β pathway in MGG8-DGCs, that is inhibited upon TGF-β inhibitor treatment. **D.** Bar diagram showing increased FMOD promoter Luc activity in MGG8-DGCs, that is inhibited upon TGF-β inhibitor treatment. p-value is calculated by unpaired t-test with Welch’s correction are indicated. P-value less than 0.05 is considered significant with *, **, *** representing p-value less than 0.05, 0.01 and 0.001 respectively.

**Supplementary Figure 6:**
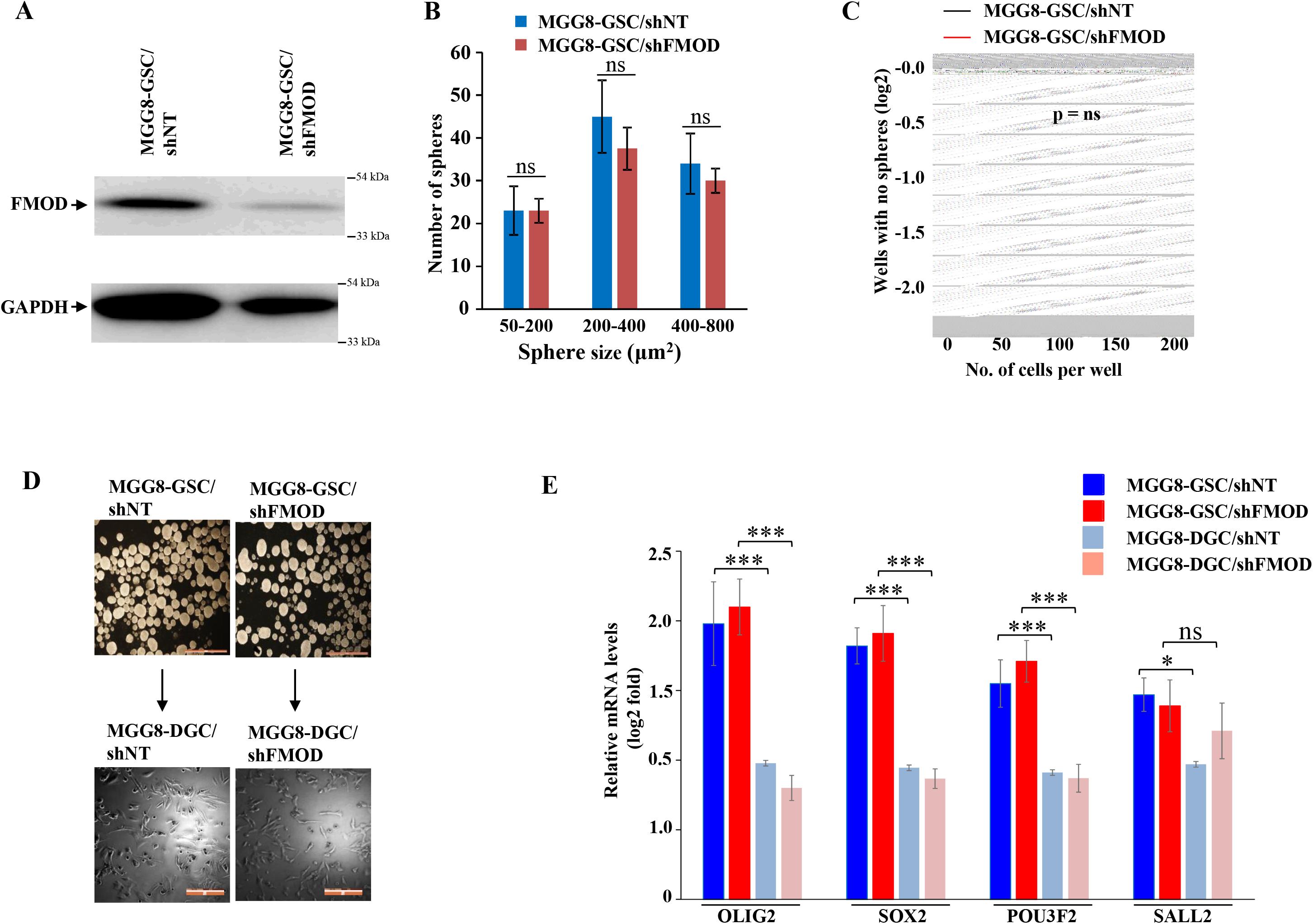
FMOD does not play a role in GSC neurosphere formation/maintenance and differentiation in human GSCs. **A.** Western blotting showing silencing of FMOD in MGG8-GSCs. **B.** Bar diagram, quantifying number of spheres, shows no significant difference in sphere formation of MGG8 GSCs between shNT and shFMOD conditions (spheres are divided into different sizes, 50-200μm2, 200-440μm2, and 400-800μm2). **C.** Limiting dilution assay shows no significant decrease (p=0.92) in the sphere-forming capacity of MGG8 GSCs between MGG8-GSC/shNT (black line) and MGG8-GSC/shFMOD (red line) conditions. **D.** Representative images showing MGG8-GSCs showing no difference in sphere formation and differentiation between shNT and shFMOD conditions. Magnification,=4X, Scale=200 μm **E.** Real-time qPCR analysis shows that the four GSC reprogramming factors (OLIG2, SOX2, POU3F2, and SALL2) have similar less expression in both MGG8-DGC/shNT and MGG8-DGC/shFMOD (depicted by solid blue and solid red respectively), but undergo an expected similar significant increase in MGG8-GSC/shNT and MGG8-GSC/shFMOD cells (depicted by striped blue and striped red bars respectively). p values calculated by unpaired t test with Welch’s correction are indicated. p value less than 0.05 is considered significant with *, **, *** representing p value less than 0.05, 0.01 and 0.001 respectively. ns stands for non-significant.

**Supplementary Figure 7:**
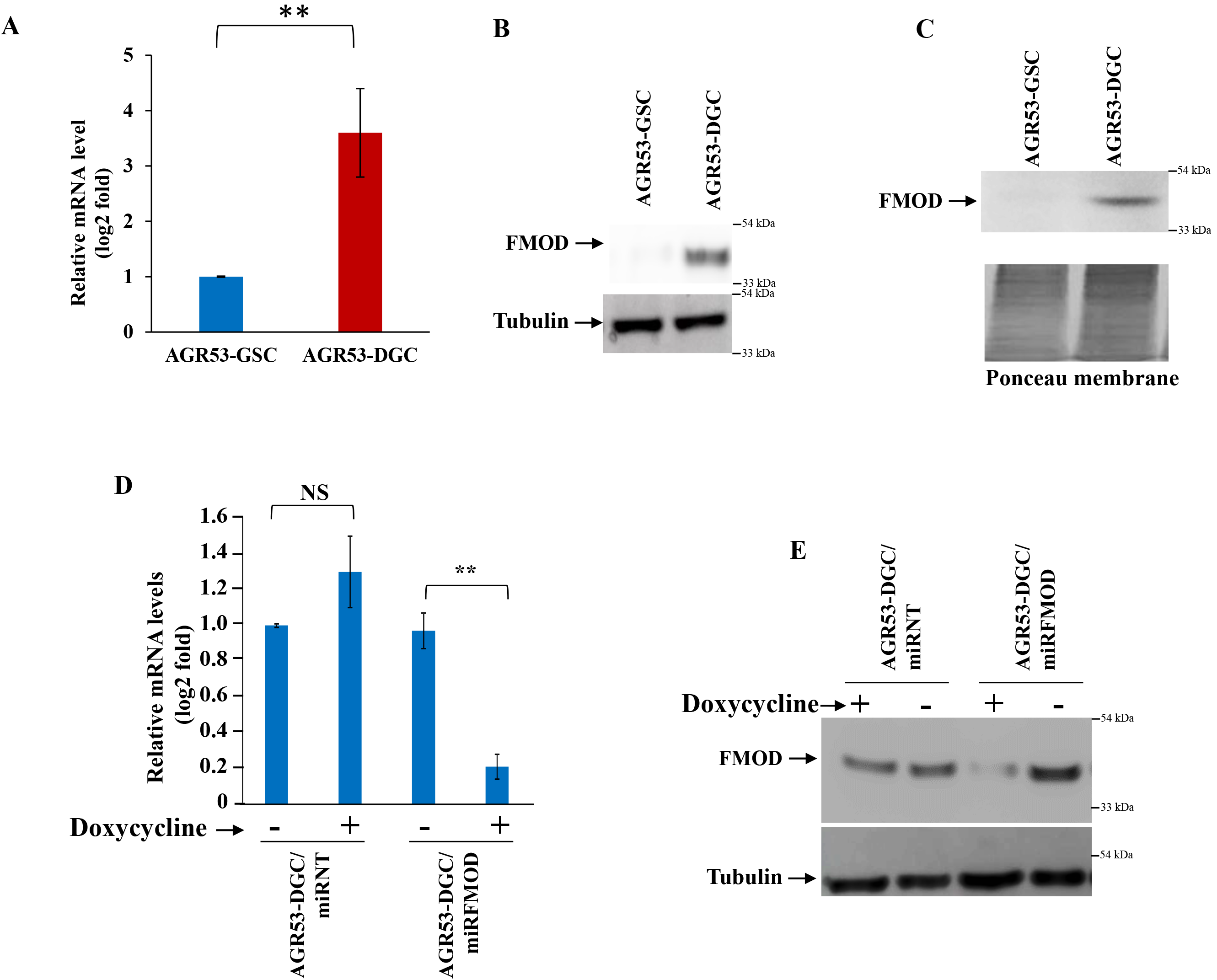
Difference in the level of FMOD expression between GSC and DGC of mouse glioma cell line AGR53 and confirmation of efficient conditional knockdown. **A.** Real time qRT-PCR analysis showing significantly higher FMOD expression in AGR53-DGCs, compared with AGR53-GSCs. **B.** Western blotting showing FMOD is expressed more at protein level intra-cellularly, in AGR53-DGCs, compared with AGR53-GSCs **C.** Western blotting showing FMOD is secreted more by AGR53-DGCs compared to AGR53-GSCs. Ponceau stained blot is used to ensure equal loading. **D.** Doxycycline addition induced FMOD knockdown at the mRNA level as well as at the **E.** protein level of AGR53-DGC/miRFMOD cells but not in the AGR53-DGC/miRNT cells. p value less than 0.05 is considered significant with *, **, *** representing p value less than 0.05, 0.01 and 0.001 respectively. ns stands for non-significant.

**Supplementary Figure 8:**
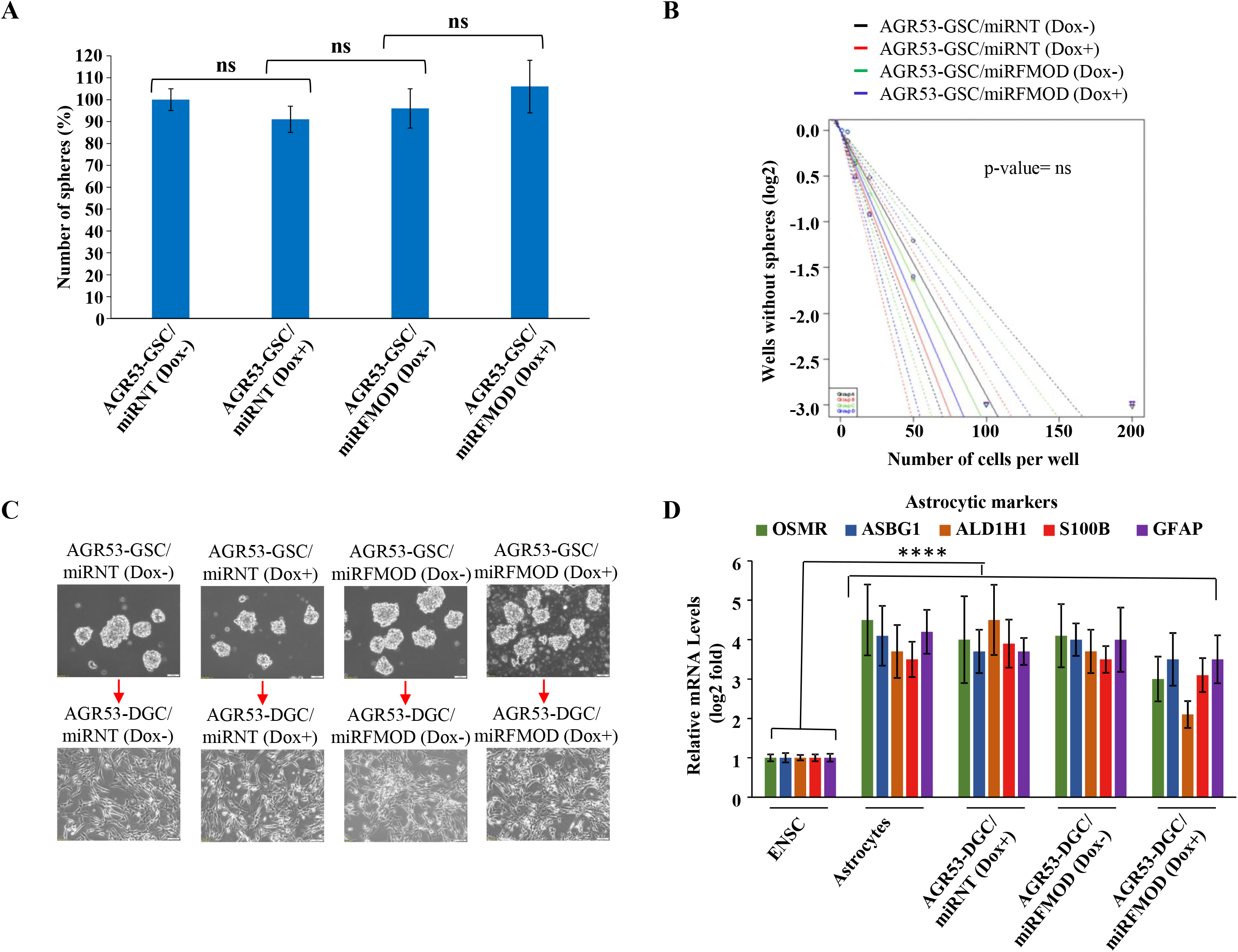
FMOD does not play a role in GSC neurosphere formation and differentiation in murine GSCs. **A.** Bar diagram, quantifying number of spheres, shows no significant difference in sphere formation of AGR53-GSC/miRNT and AGR53-GSC/miRFMOD cells with or without doxycycline treatment. **B.** Limiting dilution assay shows no significant decrease (p=0.749) in the sphere-forming capacity of AGR53-GSC/miRNT and AGR53-GSC/miRFMOD cells with or without doxycycline treatment. **C.** Representative images showing no difference in neurosphere formation and subsequent differentiation of AGR53-GSC/miRNT and AGR53-GSC/miRFMOD cells with or without doxycycline treatment. Scale= 10X, Magnification= 200 μm. **D.** Real time qRT-PCR analysis showing that the addition of doxycycline did not hamper the differentiation potential of AGR53-GSC/miRNT and AGR53-GSC/miRFMOD cells with or without doxycycline treatment, as indicated by an upregulation of the astrocytic markers in all groups of cells. Embryonic Neuronal Stem Cells (ENSCs) were used as a negative control and astrocytes were used as a positive control. p values calculated by unpaired t test with Welch’s correction are indicated. p value less than 0.05 is considered significant with *, **, *** representing p value less than 0.05, 0.01 and 0.001 respectively. ns stands for non-significant.

**Supplementary Figure 9:**
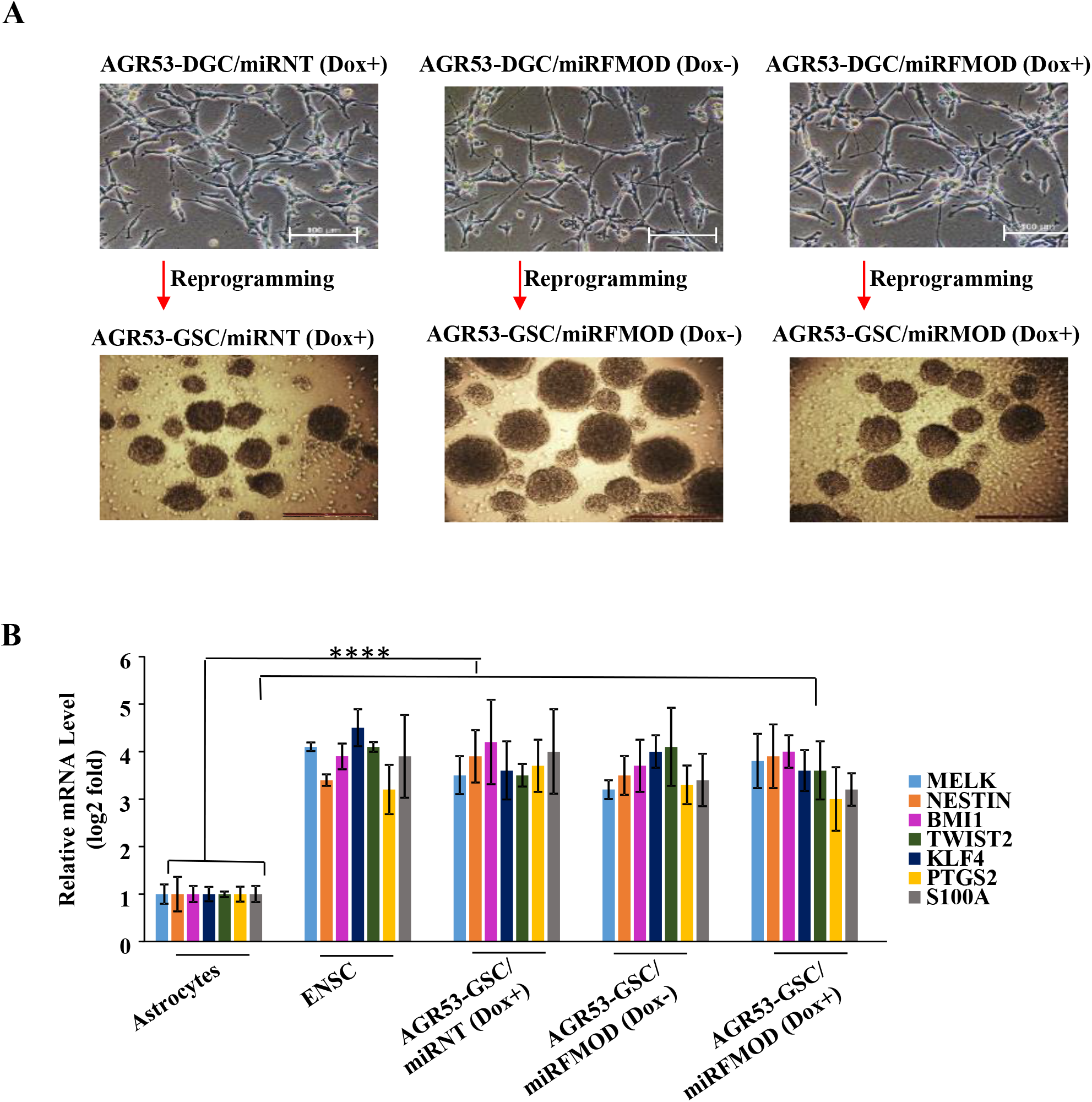
FMOD does not play a role in GSC differentiation and reprogramming of murine GSCs. **A.** Representative images showing no difference in differentiation and subsequent neurosphere formation AGR53-DGC/miRNT and AGR53-DGC/miRFMOD cells with or without doxycycline treatment. Magnification=4X, Scale= 100 μm **B. Real-time** qRT-PCR analysis showing that the addition of doxycycline did not hamper the neurosphere formation potential of AGR53-DGC/miRNT and AGR53-DGC/miRFMOD cells with or without doxycycline treatment, as indicated by an upregulation of the stem cell markers in all groups of cells. Embryonic Neuronal Stem Cells (ENSCs) were used as a positive control and astrocytes were used as a negative control. p values calculated by unpaired t-test with Welch’s correction are indicated. P-value less than 0.05 is considered significant with *, **, *** representing p-value less than 0.05, 0.01 and 0.001 respectively. ns stands for non-significant.

**Supplementary Figure 10:**
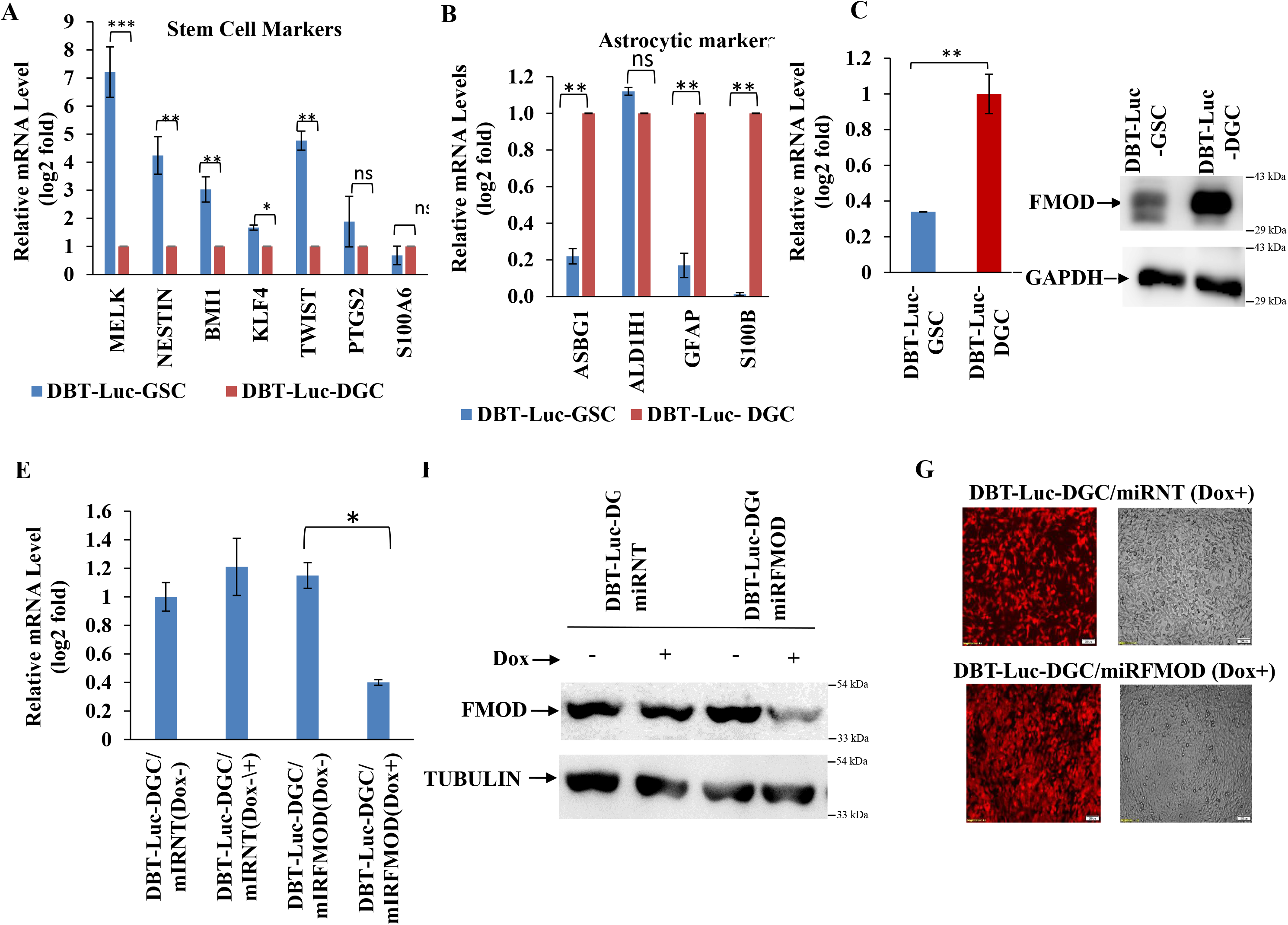
Validation of FMOD levels and knockdown in DBT-Luc mouse glioma cell-line. **A.** DBT-Luc cells were reprogrammed from DGCs to GSCs, and it was found that the mouse-stem-cell markers (MELK, NESTIN, BMI1, KLF4, TWIST2, PTGS2, and S100A6) were significantly upregulated in the DBT-Luc-GSCs, compared with the DBT-Luc-DGCs, at the mRNA level. **B.** It was also observed that the astrocytic markers (ASBG1, ALD1H1, GFAP, S100B) were significantly higher the mRNA level, in the DBT-Luc-DGCs compared with the DBT-Luc-GSCs. **C.** FMOD mRNA level was significantly higher in the DBT-Luc-DGCs compared with DBT-Luc-GSCs. **D.** FMOD protein level was also significantly higher in the DBT-Luc-DGCs compared to DBT-Luc-GSCs. **E.** Upon Doxycycline addition, a significant reduction in FMOD mRNA level was seen in DBT-Luc-DGC/miRFMOD (Dox+) group compared with DBT-Luc-DGC/miRFMOD (Dox-) group. No difference was seen in FMOD mRNA level between DBT Luc-DGC/miRNT (Dox-) and DBT-Luc-DGC/miRNT (Dox+) group of cells. **F.** Upon Doxycycline addition, a significant reduction in FMOD protein level was seen in DBT-Luc-DGC/miRFMOD(Dox+) group compared to DBT-Luc-DGC/miRFMOD(Dox-) group. No difference was seen in FMOD protein level between DBT-Luc-DGC/miRNT(Dox-) and DBT-Luc-DGC/miRNT(Dox+) group of cells. **G.** Upon Doxycycline addition, mCherry expression was seen in both DBT-Luc-DGC/miRNT and DBT-Luc-DGC/miRFMOD group of cells. Magnification-4X, Scale- 200μm. p values calculated by unpaired t-test with Welch’s correction are indicated. p-value less than 0.05 is considered significant with *, **, *** representing p-value less than 0.05, 0.01 and 0.001 respectively. ns stands for non-significant.

**Supplementary Figure 11:**
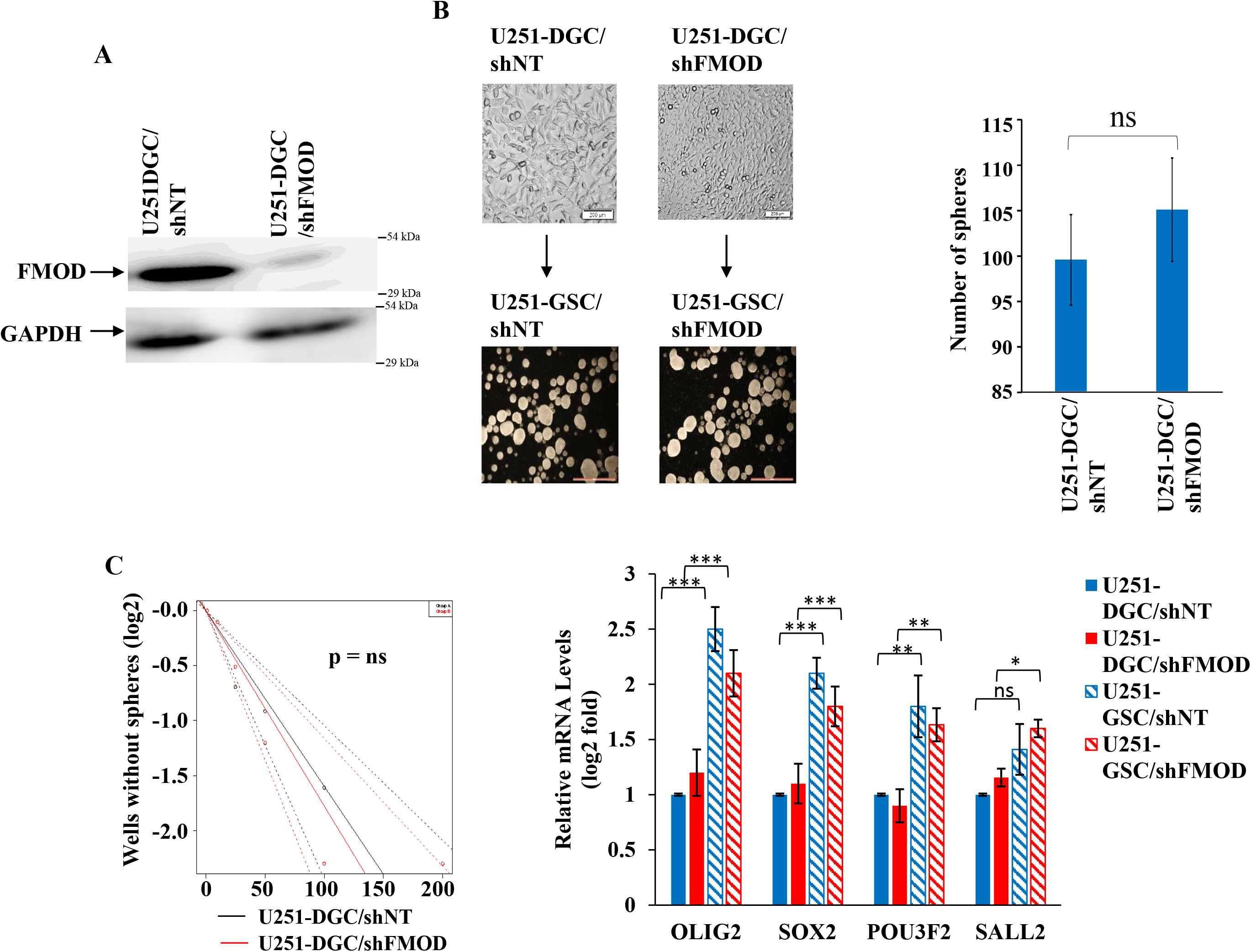
FMOD does not have a role in de-differentiation of DGCs to GSCs. **A.** U251-DGC/shNT and U251-DGC/shFMOD cells show no difference in growth in DMEM, or in neurosphere formation, when plated in stem-cell media in ultra-low attachment plates. Magnification = 10X, Scale: 100μm. Bar diagram, quantified from **A,** shows that there is no significant decrease in sphere formation of U251 de-differentiated cells (U251/GSCs) between U251-GSC/shNT and U251-GSC/shFMOD conditions. **B.** Limiting dilution assay shows no significant decrease (p=0.89) in neurosphere forming capacity of U251 neurospheres between U251-GSC/shNT (black line) and U251-GSC/shFMOD (red line) conditions. **C.** Real-time qPCR analysis shows that the four GSC reprogramming factors (OLIG2, SOX2, POU3F2, and SALL2) have similar less expression in both U251-DGC/shNT and U251-DGC/shFMOD (depicted by solid blue and solid red respectively), but undergo an expected similar significant increase in U251-GSC/shNT and U251-GSC/shFMOD cells (depicted by striped blue and striped, red bars respectively). p values calculated by unpaired t-test with Welch’s correction are indicated. P-value less than 0.05 is considered significant with *, **, *** representing p-value less than 0.05, 0.01 and 0.001 respectively. ns stands for non-significant.

**Supplementary Figure 12:**
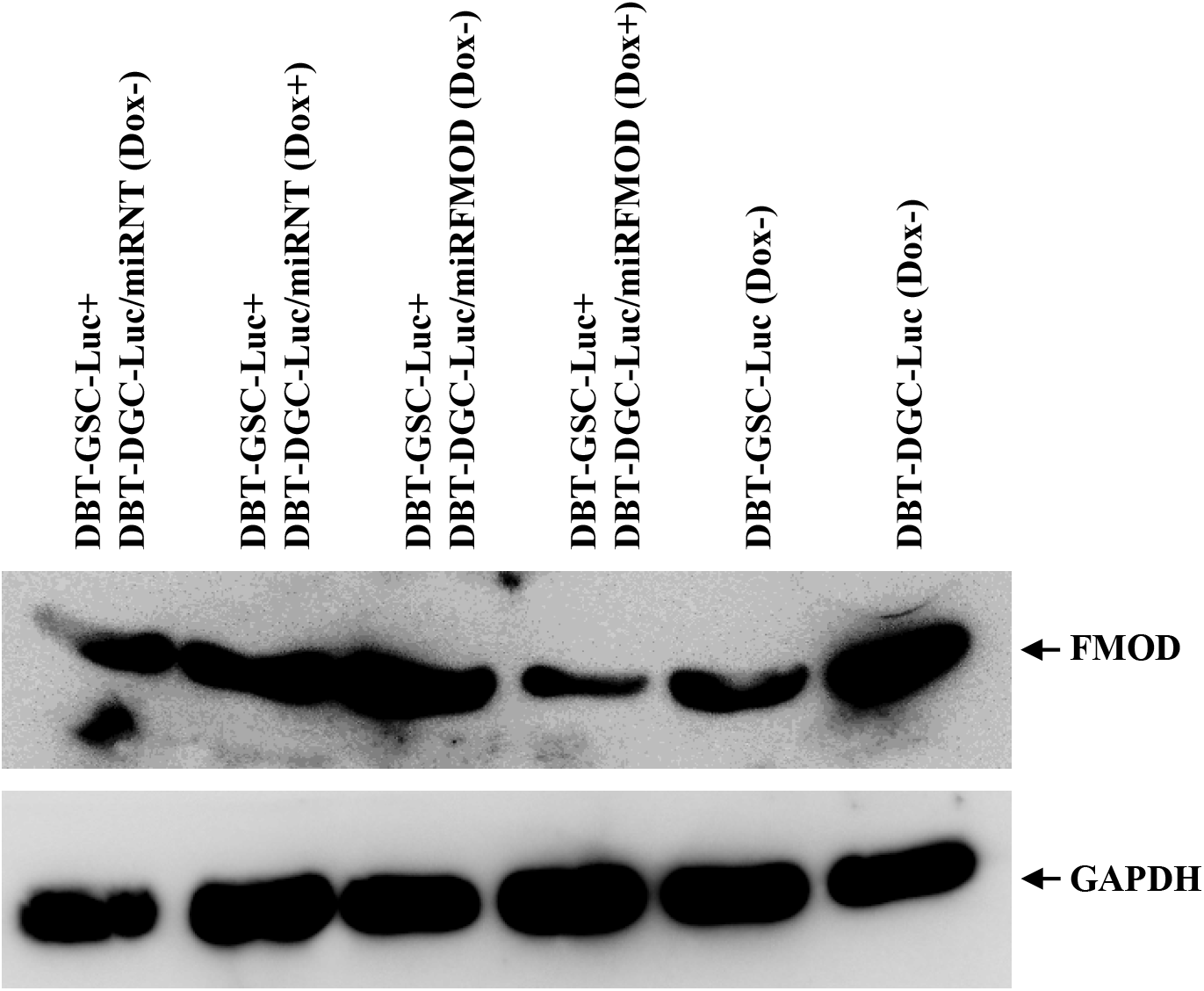
Induction with doxycycline reduces FMOD level in DBT-GSC-Luc+DBT-DGC-Luc/miRFMOD(Dox+) tumors. Western blot validating a decreased FMOD expression in tumors formed by co-injecting DBT-GSC-Luc cells with DBT-DGC-Luc/miRFMOD, upon doxycycline induction. Expression of FMOD was higher in all the other co-injection tumor groups ( DBT-GSC-Luc+DBT-DGC-Luc/miRNT(Dox+), DBT-GSC-Luc+DBT-DGC-Luc/miRNT(Dox-) DBT-GSC-Luc+DBT-DGC-Luc/miRFMOD(Dox-)) and the DBT-Luc-DGC(Dox-) tumors, while it was low in the DBT-Luc-GSC(Dox -)tumor.

**Supplementary Figure 13:**
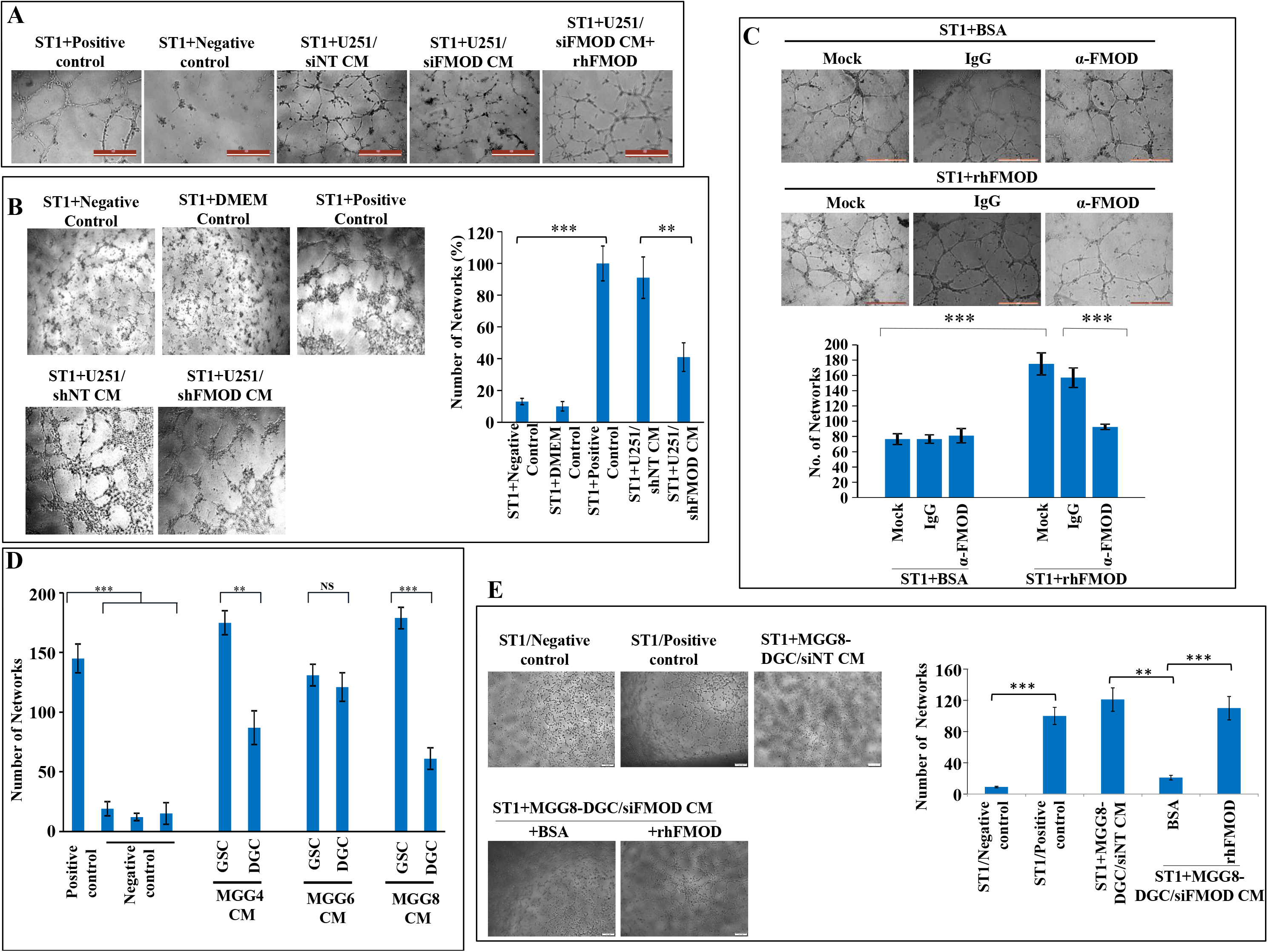
DGC-secreted FMOD induces angiogenesis by endothelial cells of various origins. **A.** Representative images of the decrease in the number of networks formed by ST1 cells when treated with U251/siFMOD CM, compared with U251/siNT CM, which is rescued by the exogenous addition of rhFMOD. **B.** Representative images (left) and quantification (right) of the decrease in the number of networks formed by ST1 cells when treated with U251/shFMOD CM, compared with U251/shFMOD CM. **C.** Representative images (top) and quantification (bottom) of the number of networks formed by the ST1 cells **show** that the cells form significantly less networks when an FMOD neutralizing antibody is added, as compared to control antibody (IgG). BSA is used as a control for rhFMOD. **D.** Quantification of the number of networks formed in *in vitro* angiogenesis assay upon treatment of ST1 cells with the CM from MGG4, MGG6 and MGG8 GSCs and their corresponding DGCs. **E.** Representative images of the decrease in the number of networks formed by ST1 cells when treated with MGG8-DGC/siFMOD CM, compared with MGG8-DGC/siFMOD CM, which is rescued by the exogenous addition of rhFMOD(left). Quantification of the number of networks formed (right). p values calculated by unpaired t-test with Welch’s correction are indicated. P-value less than 0.05 is considered significant with *, **, *** representing p-value less than 0.05, 0.01 and 0.001 respectively.

**Supplementary Figure 14:**
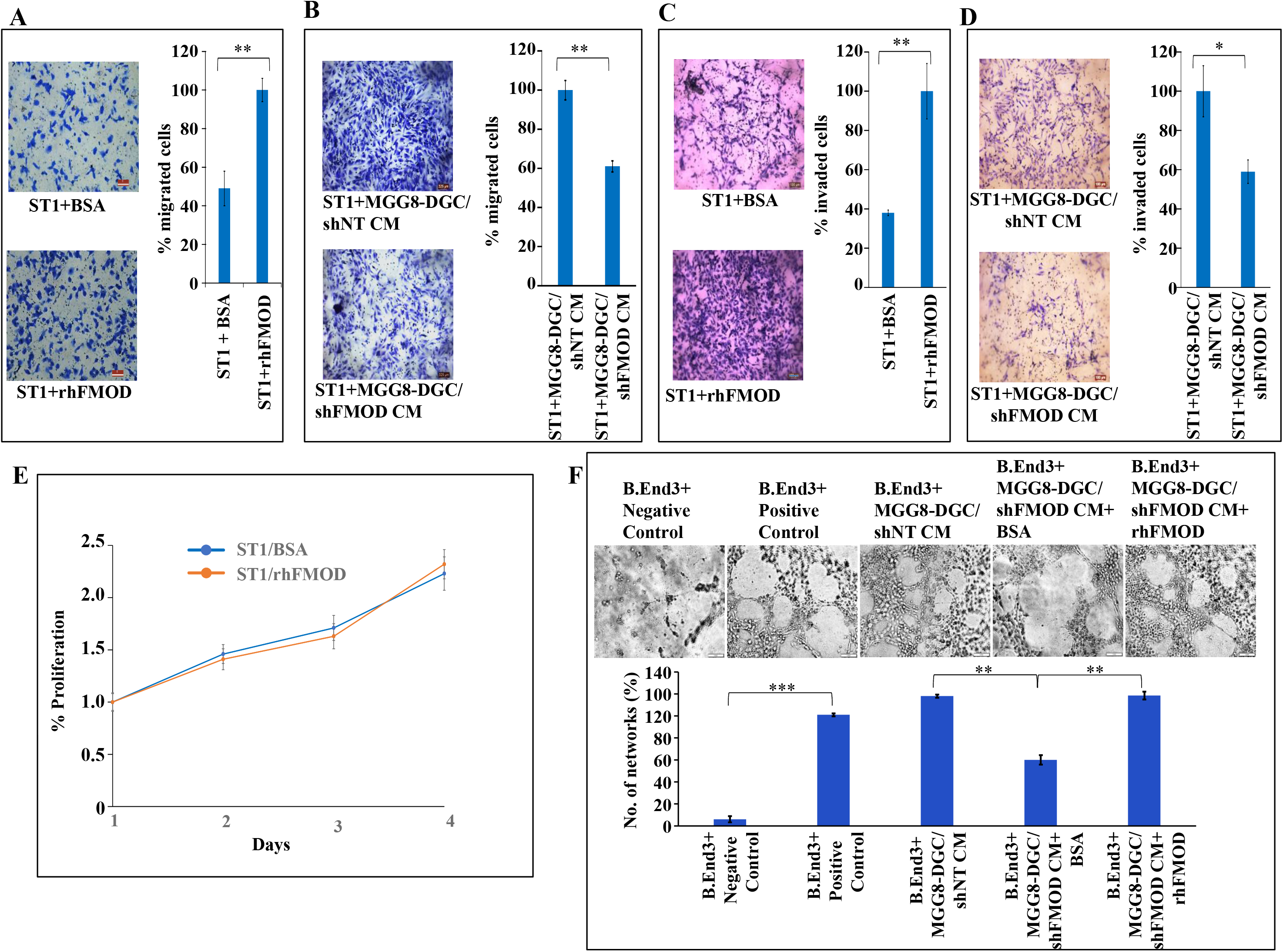
FMOD enhances angiogenesis as well as the migration and invasion, but not the proliferation of endothelial cells. **A.** Representative images (left) and quantification (right) from Boyden chamber assay showing increased migration of ST1 cells upon rhFMOD treatment over BSA. Magnification=10x scale=100μm.**B.** Representative images (left) and quantification (right) from Boyden chamber assay showing reduced migration of ST1 cells upon MGG8-DGC/shFMOD CM treatment compared with MGG8-DGC/shNT CM treatment. Magnification=10x scale=100μm.**C.** Representative images (left) and quantification (right) from Boyden chamber assay showing increased invasion of ST1 cells upon rhFMOD treatment over BSA. Magnification=10x scale=100μm.**D.** Representative images (left) and quantification (right) from Boyden chamber assay showing reduced invasion of ST1 cells upon MGG8-DGC/shFMOD CM treatment compared with MGG8-DGC/shNT CM treatment. Magnification=10x scale=100μm. **E.** MTT cell-proliferation assay shows no difference in the proliferation of ST1 cells treated with either BSA or rfFMOD, over a period of 96 hours. **F.** Representative images (top) and quantification (bottom) of the decrease in the number of networks formed by immortalized mouse brain-derived endothelial cells, B.End3, upon treatment with MGG8-DGC/shFMOD CM, compared with MGG8-DGC/shNT CM, which is rescued upon exogenous addition of rhFMOD. p values calculated by unpaired t-test with Welch’s correction are indicated. P-value less than 0.05 is considered significant with *, **, *** representing p-value less than 0.05, 0.01 and 0.001 respectively. ns stands for non-significant.

**Supplementary Figure 15:**
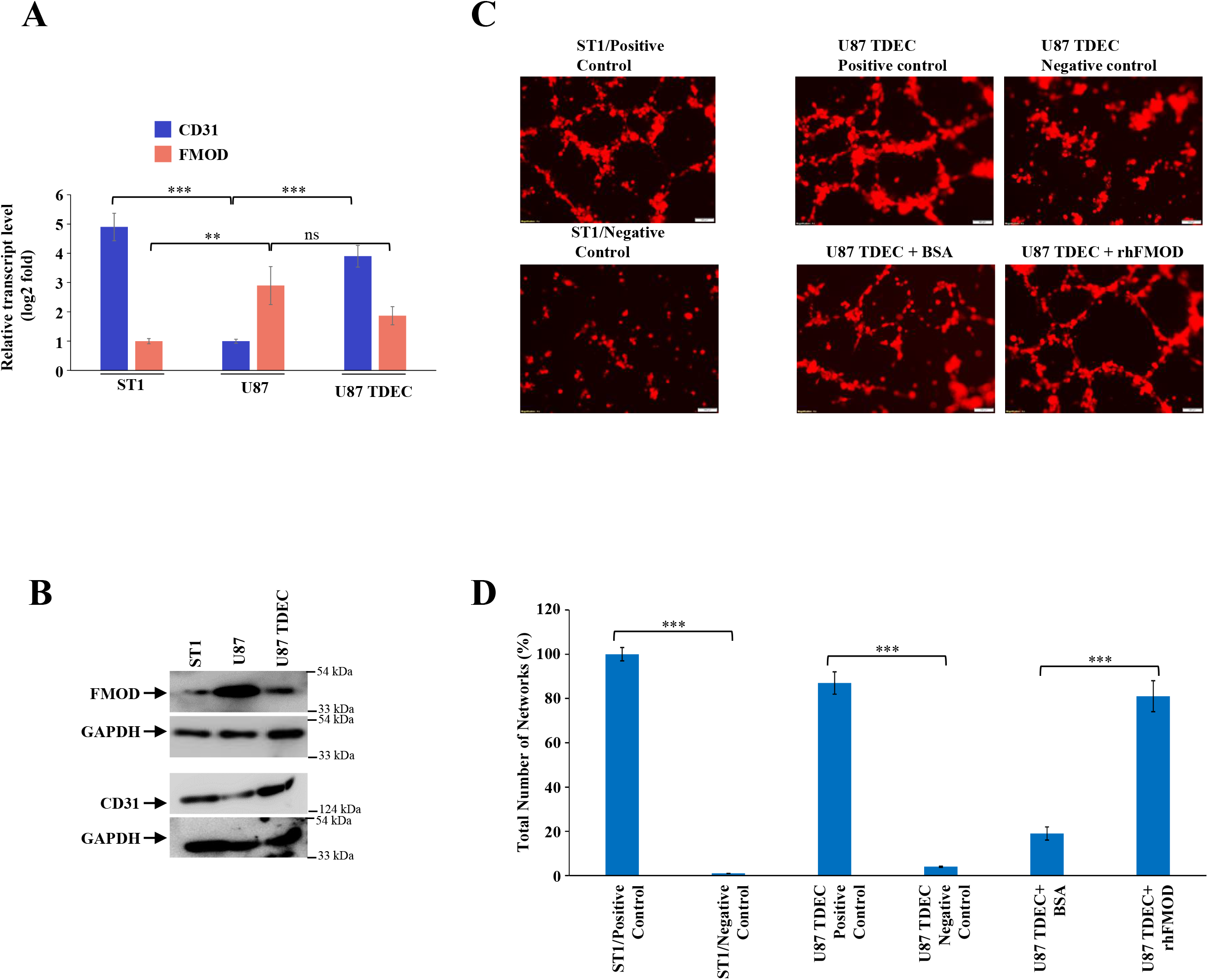
FMOD induces GBM cells to undergo transdifferentiation. **A.** Real-time qRT-PCR analysis showing the expression of CD31(blue bars) and FMOD (orange bars) in ST1 cells and U87 cells, before and after transdifferentiation (TDECs represent the transdifferentiated cells).). **B.** Western blotting showing the expression of FMOD (top) and CD31 (bottom) in U87 cells, before and after transdifferentiation (TDECs represent the transdifferentiated cells). **C**. Representative images of *in vitro* network formation by U87 and U87 TDECs upon BSA and rhFMOD treatments. Magnification 10X, Scale bar = 100 μm. **D.** Quantification of the number of complete networks formed in **C.** p-value is calculated by unpaired t-test with Welch’s correction are indicated. P-value less than 0.05 is considered significant with *, **, *** representing p value less than 0.05, 0.01 and 0.001 respectively. ns stands for non-significant.

**Supplementary Figure 16:**
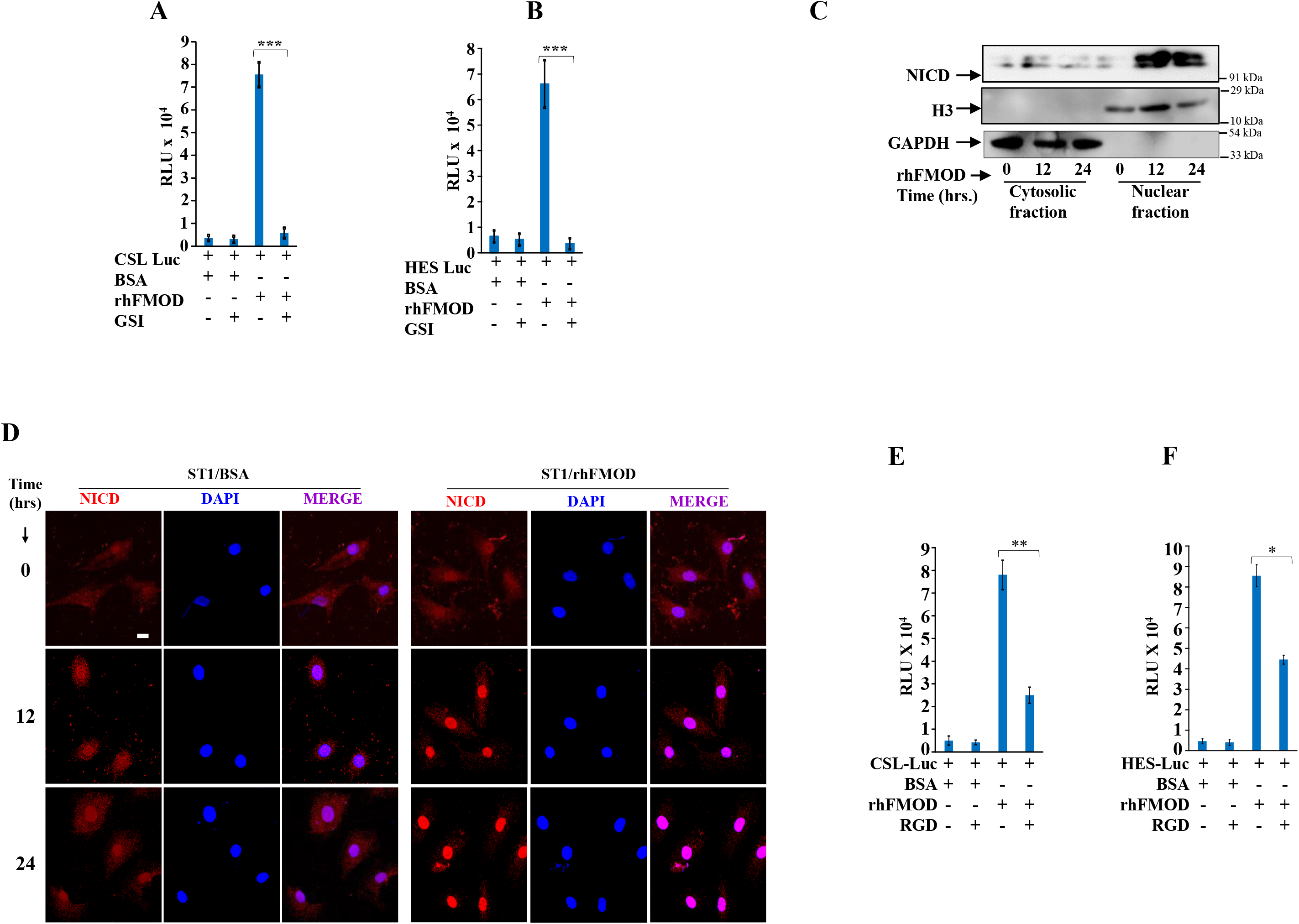
FMOD induces activation of Notch signaling in endothelial cells. **A.** rhFMOD induces a significant increase in CSL Luc (Notch pathway-dependent reporter luciferase construct) activity in ST1 cells, an effect reduced by cell pretreatment with γ-secretase inhibitor (GSI, 10μM), a Notch pathway inhibitor. **B**. rhFMOD induces a significant increase in HES Luc activity (Notch pathway-dependent reporter (HES1 promoter) luciferase construct), which is reduced when ST1 cells are pre-treated with GSI. **C.** Western blotting showing a time-dependent translocation of the Notch intracellular domain (NICD) from the cytoplasm to the nucleus in ST1 cells, upon treatment with rhFMOD. Histone H3 was used as a nuclear loading control, GAPDH as a cytoplasmic loading control. **D.** Immunocytochemical analysis showing that rhFMOD treatment of endothelial cells causes translocation of NICD from the cytoplasm to the nucleus in a time-dependent manner. BSA is used as a control. Magnification = 40X, Scale bar= 50μm. **E.** rhFMOD-mediated increase in the reporter-luciferase activity of CSL Luc significantly decreases in ST1 endothelial cells pre-treated with the RGD peptide. **F.** rhFMOD-mediated increase in the reporter-luciferase activity of HES Luc significantly decreases in ST1 cells pre-treated with the RGD peptide. p-value is calculated by unpaired t-test with Welch’s correction are indicated. P-value less than 0.05 is considered significant with *, **, *** representing p-value less than 0.05, 0.01 and 0.001 respectively. ns stands for non-significant.

**Supplementary Figure 17:**
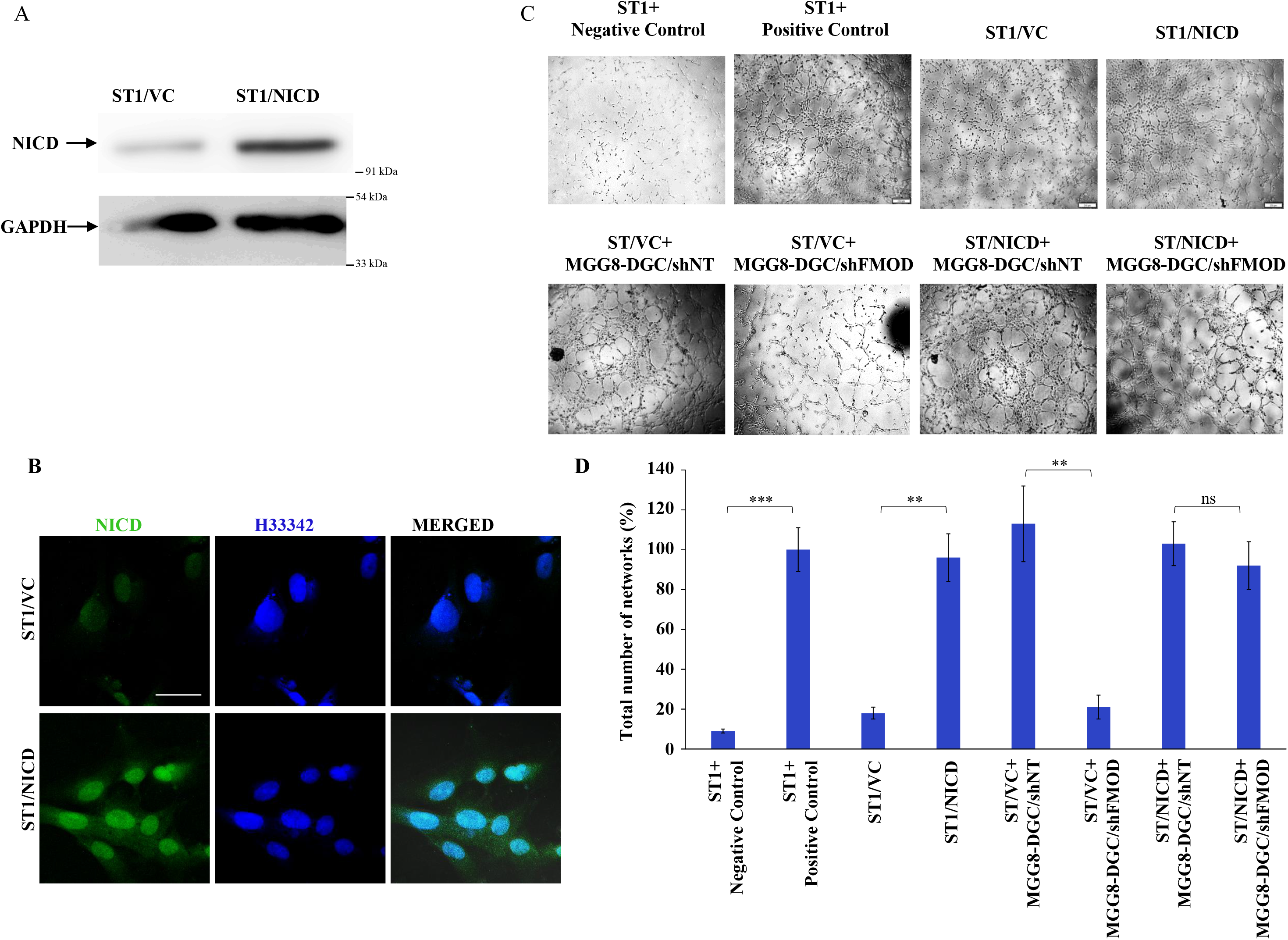
ST1 cells stably expressing NICD are independent of FMOD in forming angiogenic networks. **A.** Western blotting confirming the overexpression of NICD in ST1 cells. **B.** Immunocytochemistry analysis confirming the overexpression of NICD in ST1 cells. **C.** Representative images of network formation by control and NICD overexpressed ST1 cells in the presence of BSA and rhFMOD. Magnification = 63x, Scale=20 μm. **D**. Quantification of the number of networks formed in **C.** p-values calculated by unpaired t-test with Welch’s correction are indicated. P-value less than 0.05 is considered significant with *, **, *** representing p-value less than 0.05, 0.01 and 0.001, respectively. ns stands for non-significant.

**Supplementary Figure 18:**
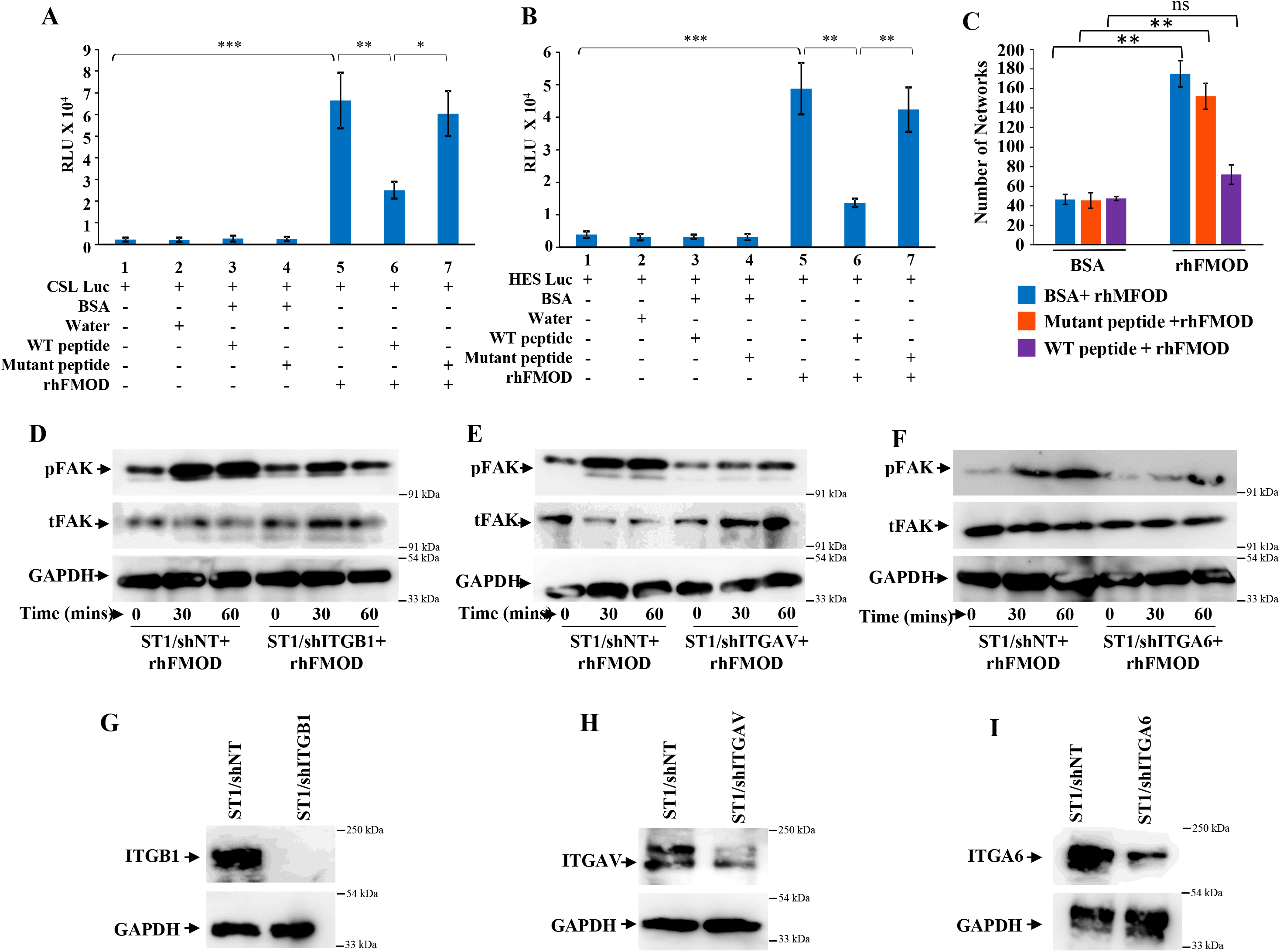
FMOD-Type I collagen interaction is crucial for FMOD-mediated activation of downstream signaling pathways. **A.** Bar diagram showing that rhFMOD cannot activate Notch-dependent reporter luciferase activity of CSL Luc in ST1 cells in the presence of WT peptide but can do so in the presence of mutant peptide. **B.** Bar diagram showing that rhFMOD cannot activate Notch-dependent reporter luciferase activity of HES Luc in ST1 cells in the presence of WT peptide but can do so in the presence of mutant peptide. **C.** Bar diagram quantifying the number of networks formed in *in vitro* angiogenesis assay shows that the WT (wild-type) peptide competes with rhFMOD to bind to Type I Collagen, whereas the mutant peptide cannot. Hence, rhFMOD induces more angiogenesis in the presence of the mutant peptide, as Type I Collagen is free for it to bind. However, because of the competition with the WT peptide, rhFMOD less efficiently induces angiogenesis in endothelial cells. **D.** Western blotting shows reduced induction in pFAk levels in rhFMOD-treated ST1/shITGB1 cells compared with the rhFMOD-treated ST1/shNT cells, where pFAK levels increae at 30 and 60 mins, while total FAK remains constant in both the cases. **E.** Western blotting shows reduced induction in pFAk levels in rhFMOD-treated ST1/shITGAV cells compared with the rhFMOD-treated ST1/shNT cells, where pFAK levels increae at 30 and 60 mins, while total FAK remains constant in both the cases. **F.** Western blotting shows reduced induction in pFAk levels in rhFMOD-treated ST1/shITGA6 cells compared with the rhFMOD-treated ST1/shNT cells, where pFAK levels increae at 30 and 60 mins, while total FAK remains constant in both the cases. **G.** Western blotting showing knockdown of ITGB1 in shITGB1 transduced ST1 cells compared with ST1/shNT cells. **H.** Western blotting showing knockdown of ITGAV in shITGAV transduced ST1 cells compared with ST1/shNT cells. **I.** Western blotting showing knockdown of ITGA6 in shITGA6 transduced ST1 cells compared with ST1/shNT cells. p values calculated by unpaired t test with Welch’s correction are indicated. p value less than 0.05 is considered significant with *, **, *** representing p value less than 0.05, 0.01 and 0.001 respectively. ns stands for non-significant.

**Supplementary Figure 19:**
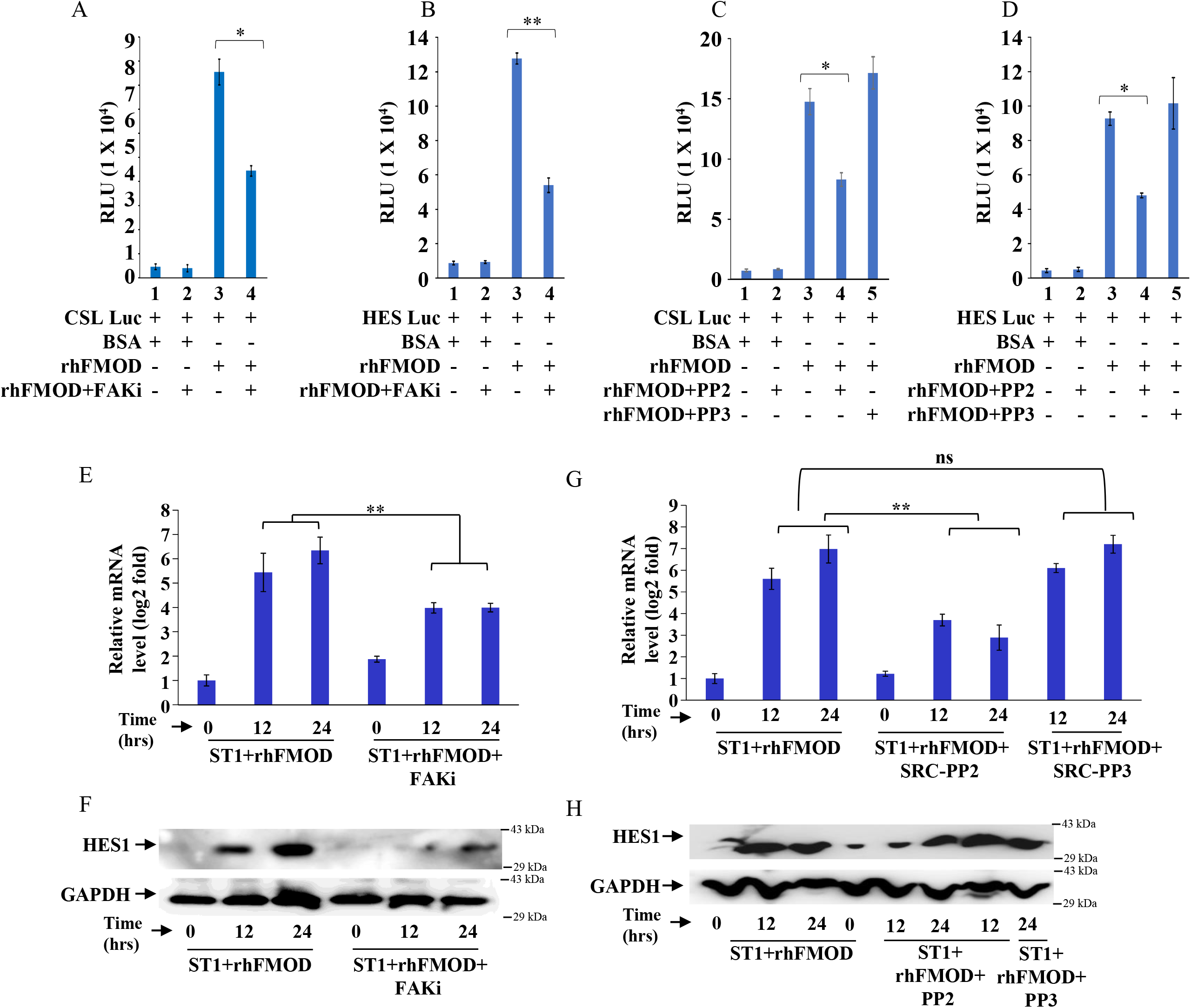
FMOD-mediated activation of Integrin-dependent Notch signaling in endothelial cells involve the integrin pathway downstream molecules FAK and Src. **A.** Bar diagram showing that rhFMOD cannot activate Notch dependent reporter luciferase activity of CSL Luc in ST1 cells pre-treated with FAK inhibitor. **B.** Bar diagram showing that rhFMOD cannot activate Notch dependent reporter luciferase activity of HES Luc in ST1 cells pre-treated with FAK inhibitor. **C.** Bar diagram showing that rhFMOD cannot activate Notch dependent reporter luciferase activity of CSL Luc in ST1 cells pre-treated with the specific Src inhibitor PP2, but not when treated with the inactive structural analog, PP3. **D.** Bar diagram showing that rhFMOD cannot activate Notch dependent reporter luciferase activity of HES Luc in ST1 cells pre-treated with the specific Src inhibitor PP2, but not when treated with the inactive structural analog, PP3. **E.** Real-time qRT-PCR analysis shows that rhFMOD treatment of ST1 cells cause a time-dependent increase in HES1 mRNA which is inhibited in cells pre-treated with FAK inhibitor. **F.** Western blotting showing that rhFMOD treatment of ST1 cells cause a time-dependent increase in HES1 protein which is inhibited in cells pre-treated with FAK inhibitor. **G.** Real-time qRT-PCR analysis shows that rhFMOD treatment of ST1 cells cause a time-dependent increase in HES1 mRNA which is inhibited in cells pre-treated with the specific Src inhibitor PP2, but not when treated with the inactive structural analog, PP3. **H.** Western blotting showing that rhFMOD treatment of ST1 cells cause a time-dependent increase in HES1 protein which is inhibited in cells pre-treated with the specific Src inhibitor PP2, but not when treated with the inactive structural analog, PP3. p value less than 0.05 is considered significant with *, **, *** representing p value less than 0.05, 0.01 and 0.001 respectively.

**Supplementary Figure 20:**
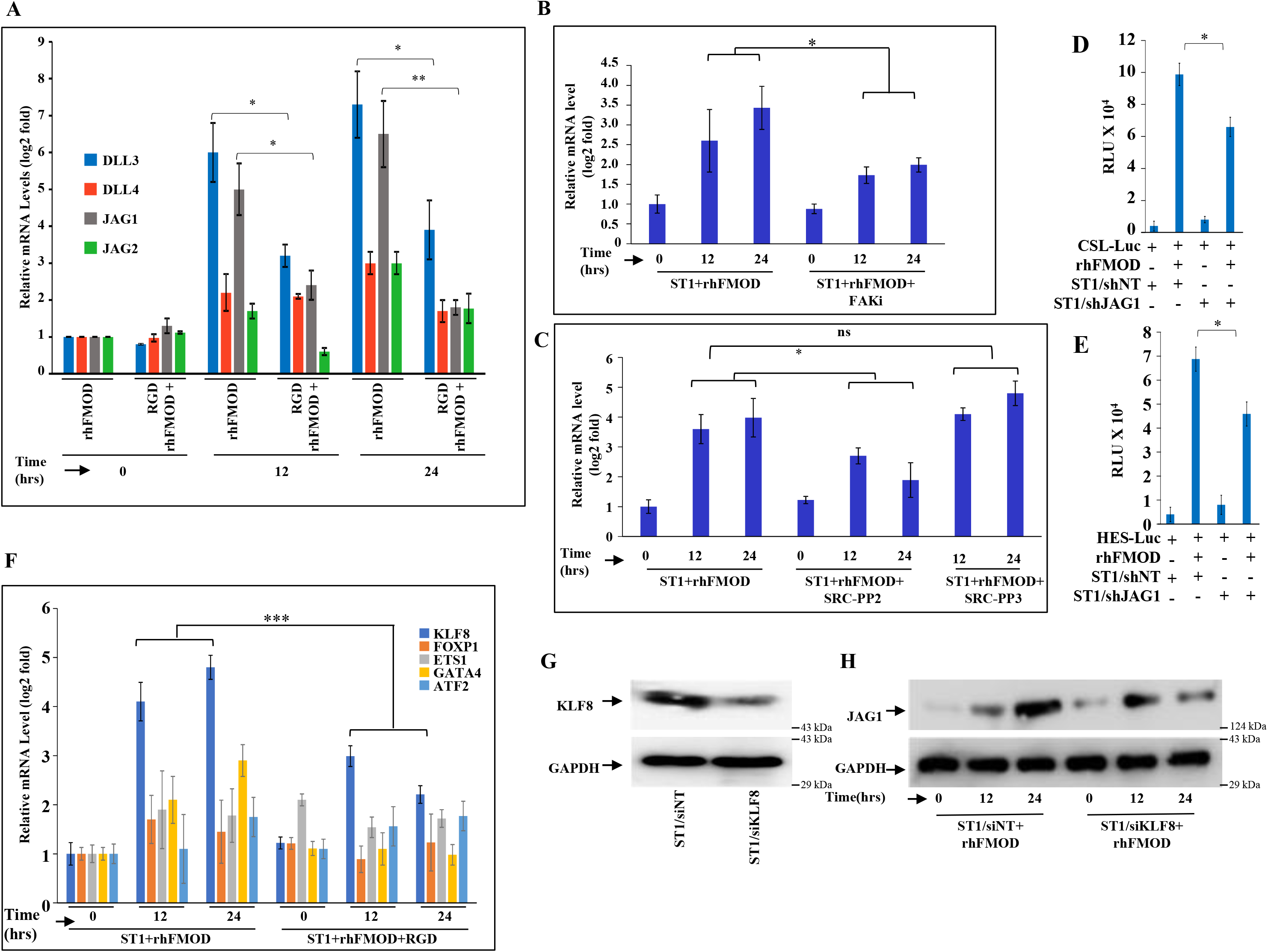
FMOD-mediated crosstalk of the integrin and Notch signaling pathways occurs via JAG1 upregulation. **A.** Real-time qRT-PCR analysis shows the rhFMOD-mediated increase in the mRNA levels of Notch ligands in a time-dependent manner. In cells that are pre-treated with RGD (10 μM), the rhFMOD-mediated increase of JAG1 and DLL3 are significantly reduced. **B.** Real-time qRT-PCR analysis shows that rhFMOD treatment of ST1 cells causes a time-dependent increase in JAG1 mRNA which is inhibited in cells pre-treated with FAK inhibitor. **C.** Real-time qRT-PCR analysis shows that rhFMOD treatment of ST1 cells cause a time-dependent increase in JAG1 mRNA which is inhibited in cells pre-treated with the specific Src inhibitor PP2, but not when treated with the inactive structural analog, PP3. **D.** rhFMOD-mediated CSL-Luc reporter-luciferase activity is significantly decreased in ST1/shJAG1 cells compared with ST1/shNT cells. **E.** rhFMOD-mediated HES-Luc reporter-luciferase activity is significantly decreased in ST1/shJAG1 cells compared with ST1/shNT cells. **F.** Real-time qRT-PCR analysis shows that rhFMOD treatment of ST1 upregulates 2 out of a set of 5 TFs in ST1, of which the expression of KLF8 (blue bar) shows a significant inhibition when the cells are pre-treated with RDG. **G.** Western blotting showing knockdown of KLF8 in siKLF8 transfected ST1 cells compared with ST1/siNT cells. **H.** Western blotting showing that rhFMOD treatment of ST1 cells cause a time-dependent increase in JAG1 protein which is inhibited in cells are silenced for KLF8. p value less than 0.05 is considered significant with *, **, *** representing p value less than 0.05, 0.01 and 0.001 respectively. ns stands for non-significant.

**Supplementary Figure 21:**
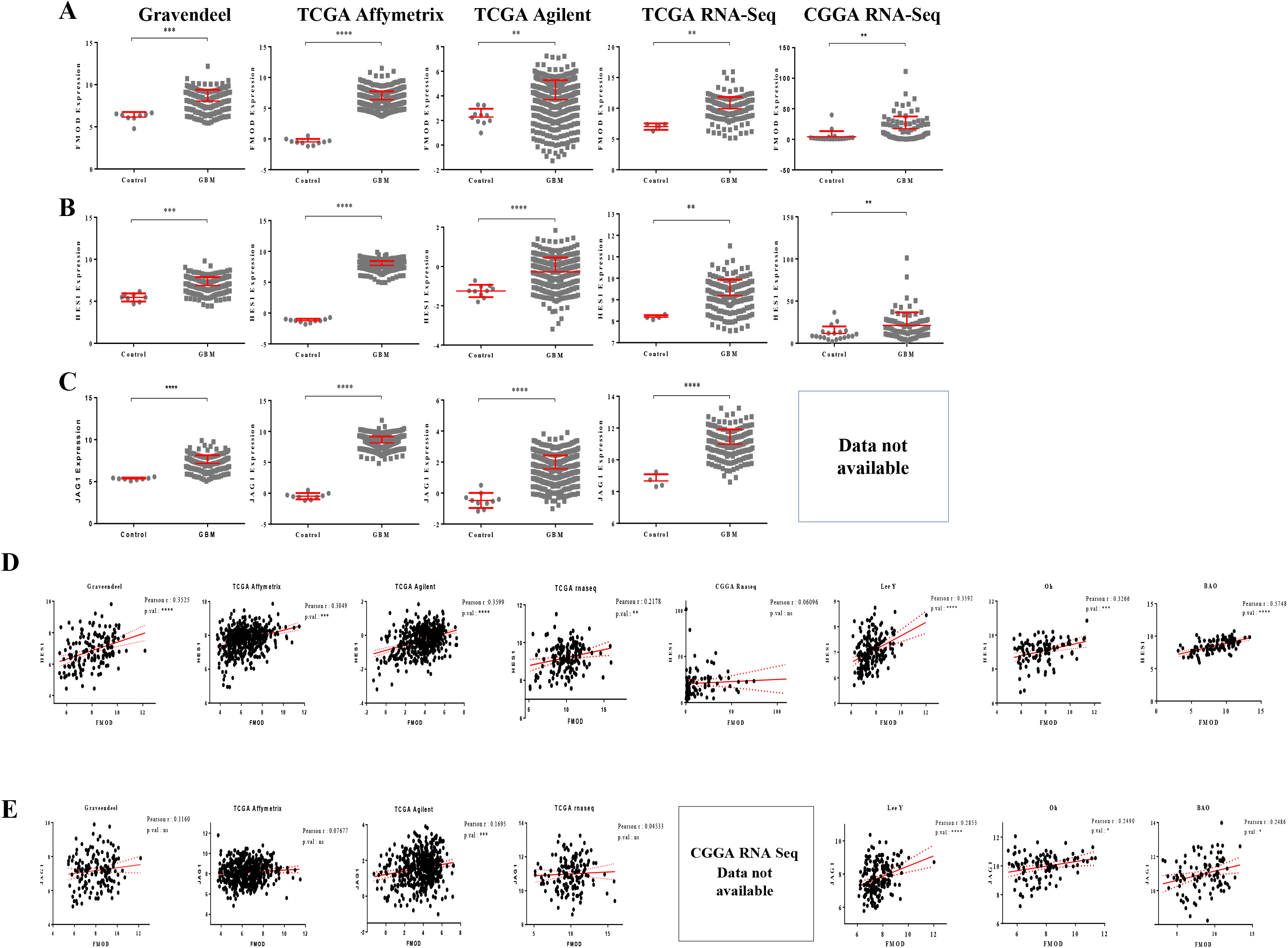
Clinical relevance of the FMOD-JAG1-HES1 signaling. **A.** FMOD is upregulated at the transcript level in GBM samples over normal samples across multiple publicly available datasets. **B.** HES1 is upregulated at the transcript level in GBM samples over normal samples across multiple publicly available datasets. **C.** JAG1 is upregulated at the transcript level in GBM samples over normal samples across multiple publicly available datasets. **D.** FMOD and HES1 mRNAs are significantly positively correlated across multiple publicly available datasets. **E.** FMOD and JAG1 mRNAs are significantly positively correlated across multiple publicly available datasets.

**Supplementary Figure 22:**
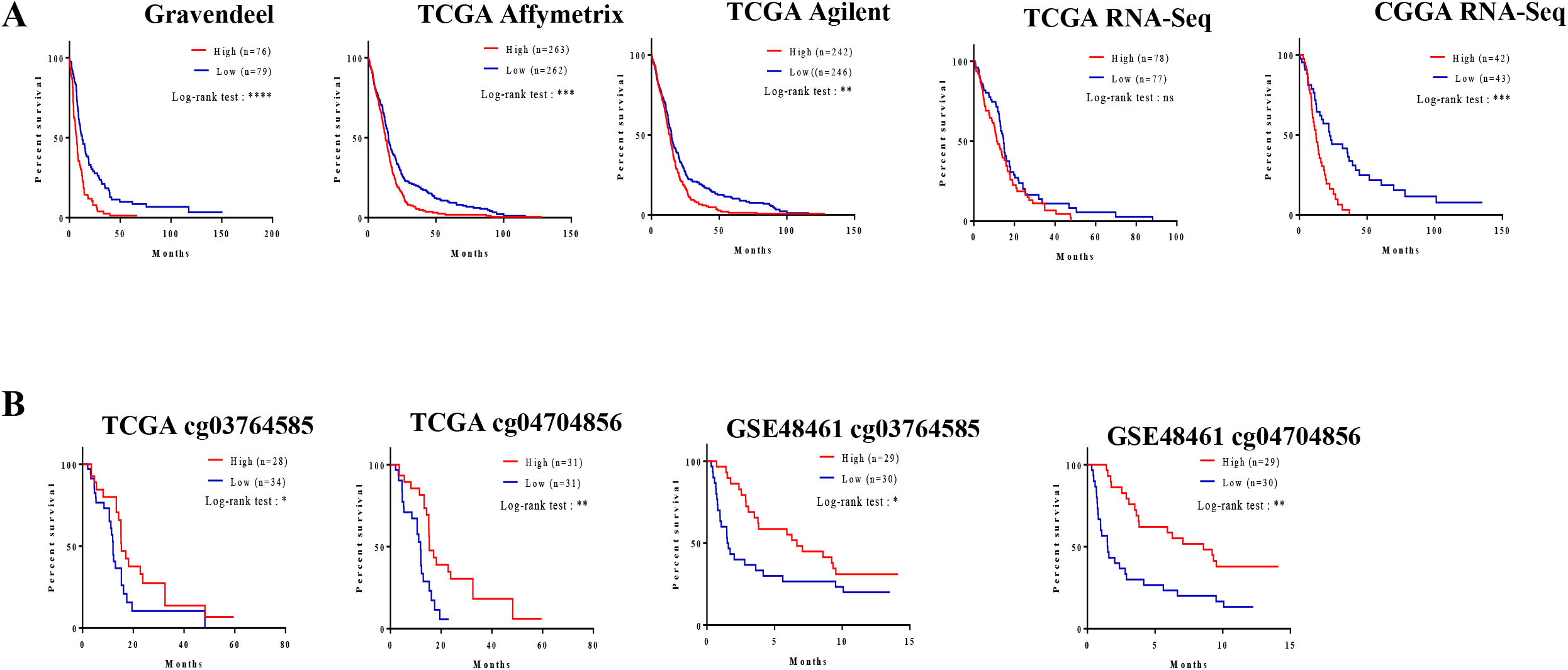
Clinical relevance of the FMOD-JAG1-HES1 signaling. **A.** High FMOD mRNA levels in patients show poorer survival than low FMOD patients, in multiple publicly available datasets. **B.** FMOD promoter hypomethylation (in CpGs cg03764585 and cg04704856) in patients show poorer survival than low patients with FMOD promoter hypermethylation, in multiple publicly available datasets.

**Supplementary Figure 23:**
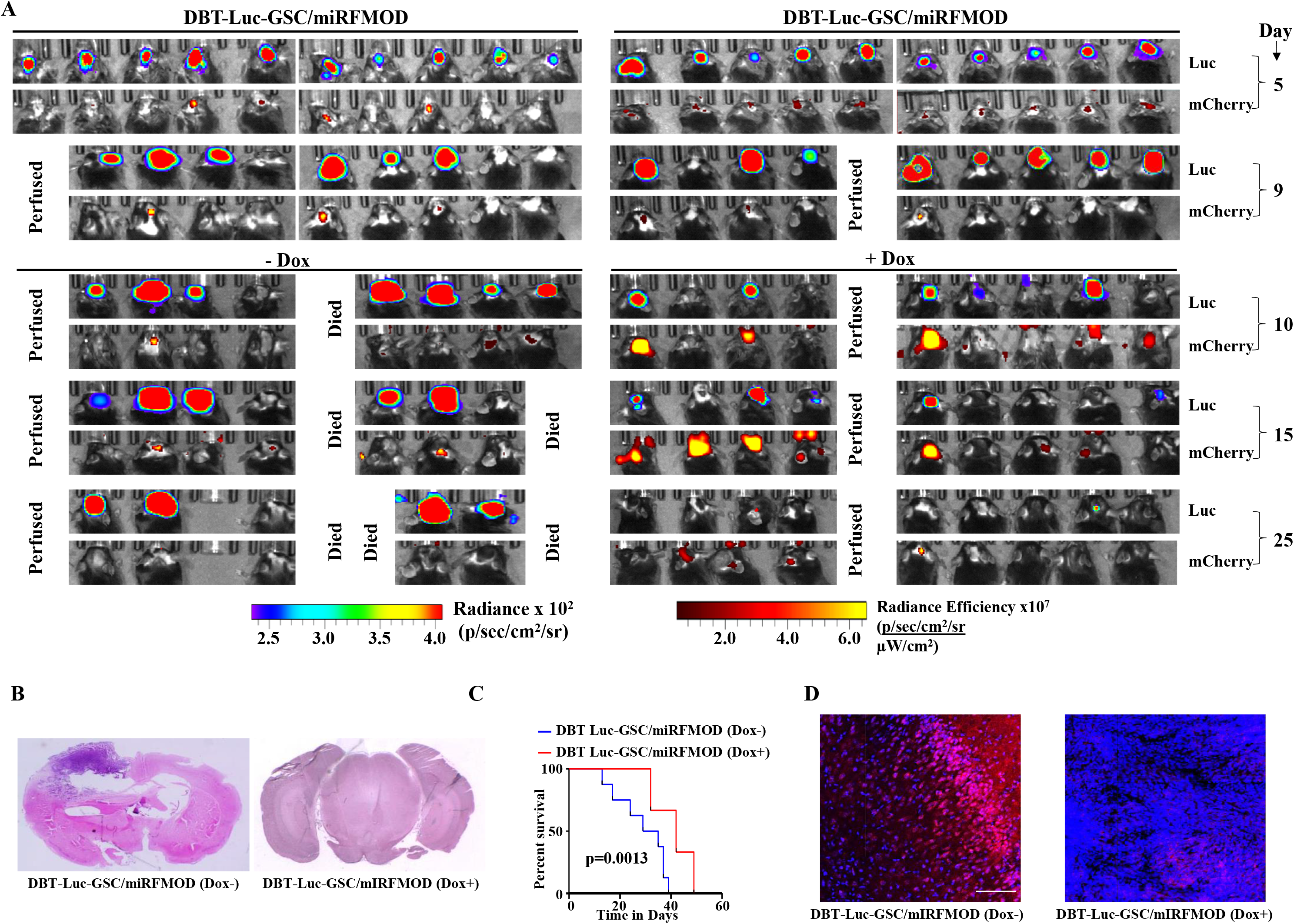
Conditional silencing of FMOD in DGCs formed *de novo* by GSC-initiated tumors inhibits tumor growth. **A.** *In vivo* fluorescent imaging of mice injected with either DBT-Luc-GSC/miRNT or DBT-Luc-GSC/miRFMOD cells. **B.** Haematoxylin and Eosin staining shows a larger tumor (depicted by dark blue color due to extremely high cellular density) in mice brain injected with DBT-Luc-GSC/miRFMOD (Dox-), but not in DBT-Luc-GSC/miRFMOD (Dox+) cells. **C.** Kaplan-Meier survival graphs showing the survival of mice injected with DBT-Luc-GSC/miRFMOD(Dox-) or DBT-Luc-GSC/miRFMOD (Dox+) cells. **D.** Immunohistochemical analysis showing FMOD expression in brains of mice injected with DBT-Luc-GSC/miRFMOD(Dox-) or DBT-Luc-GSC/miRFMOD(Dox+) cells. Red represents FMOD and blue represents H33342. The merged images have been shown for representation. Magnification = 20x, Scale=100 μm.

**Supplementary Figure 24:**
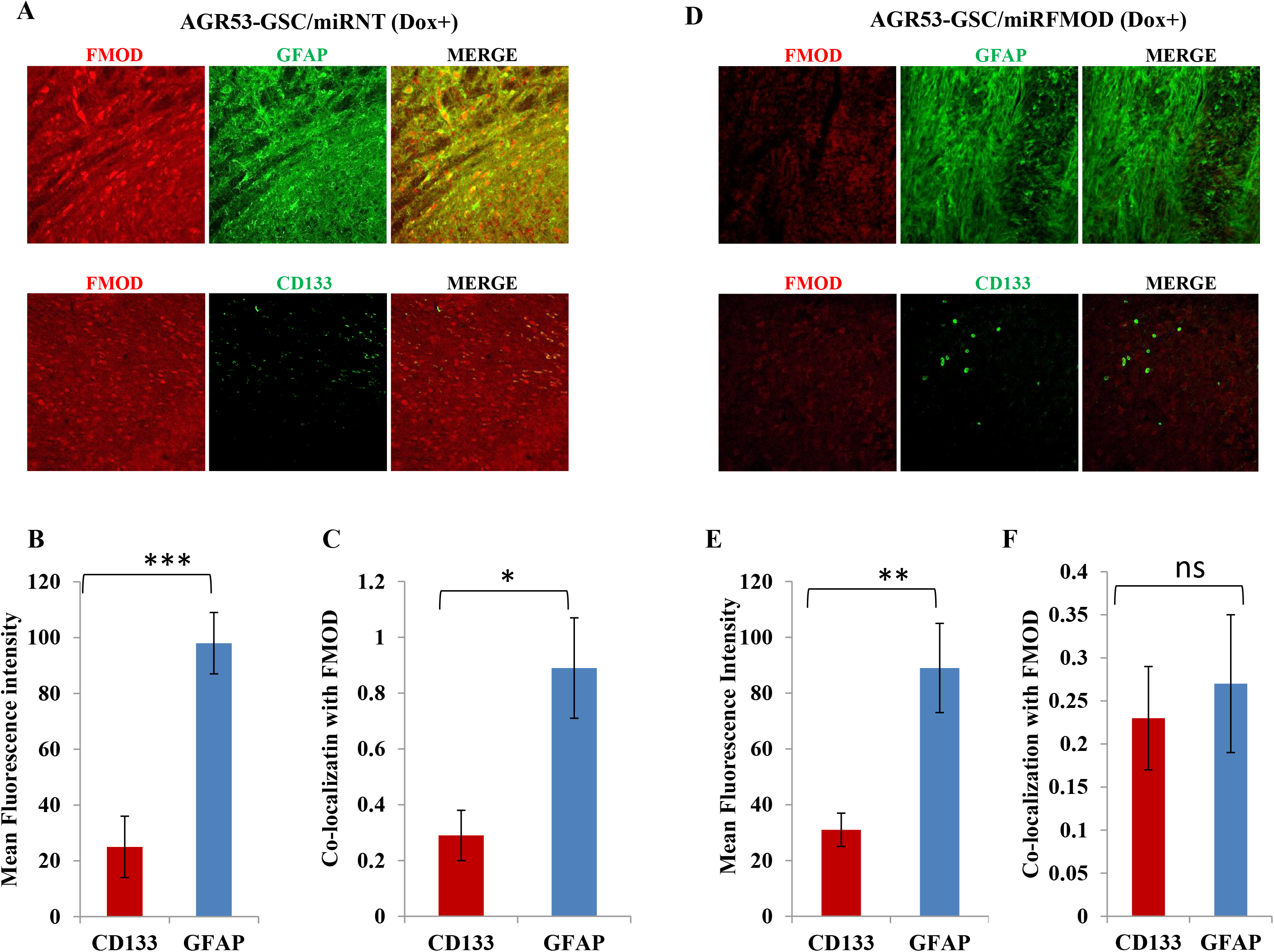
FMOD-silencing does not hamper the differentiation potential of tumor cells in murine glioma models. **A.** Immunohistochemical analysis of brain-derived sections obtained from mice injected with AGR53-GSC/miRNT cell show FMOD, GFAP, and CD133 expression, and co-localization of FMOD with GFAP, and not with CD133 (left, after doxycycline injection). **B.** Bar diagram quantifying mean fluorescence intensity of CD133 and GFAP in AGR53-GSC/miRNT cells after doxycycline injection. **C.** Bar diagram quantifying co-localization of FMOD with CD133 and GFAP, respectively, in AGR53-GSC/miRNT cells after doxycycline injection. **D.** Immunohistochemical analysis of brain-derived sections obtained from mice injected with AGR53-GSC/miFMOD cell show low FMOD, and hifg GFAP, and CD133 expression, and loss of co-localization of FMOD with GFAP (right, after doxycycline injection). **E.** Bar diagram quantifying mean fluorescence intensity of CD133 and GFAP in AGR53-GSC/miRFMOD cells after doxycycline injection. **F.** Bar diagram quantifying co-localization of FMOD with CD133 and GFAP, respectively, in AGR53-GSC/miRNT cells after doxycycline injection. Magnification for panels **A** and **D** =20 X, Scale= 100 μm. p-values for panels **B, C, E**, and **F** are calculated by Student’s t-test, p-value are represented by *, **, *** indicate values <0.05, <0.01, and <0.001 respectively.

**Supplementary Figure 25:**
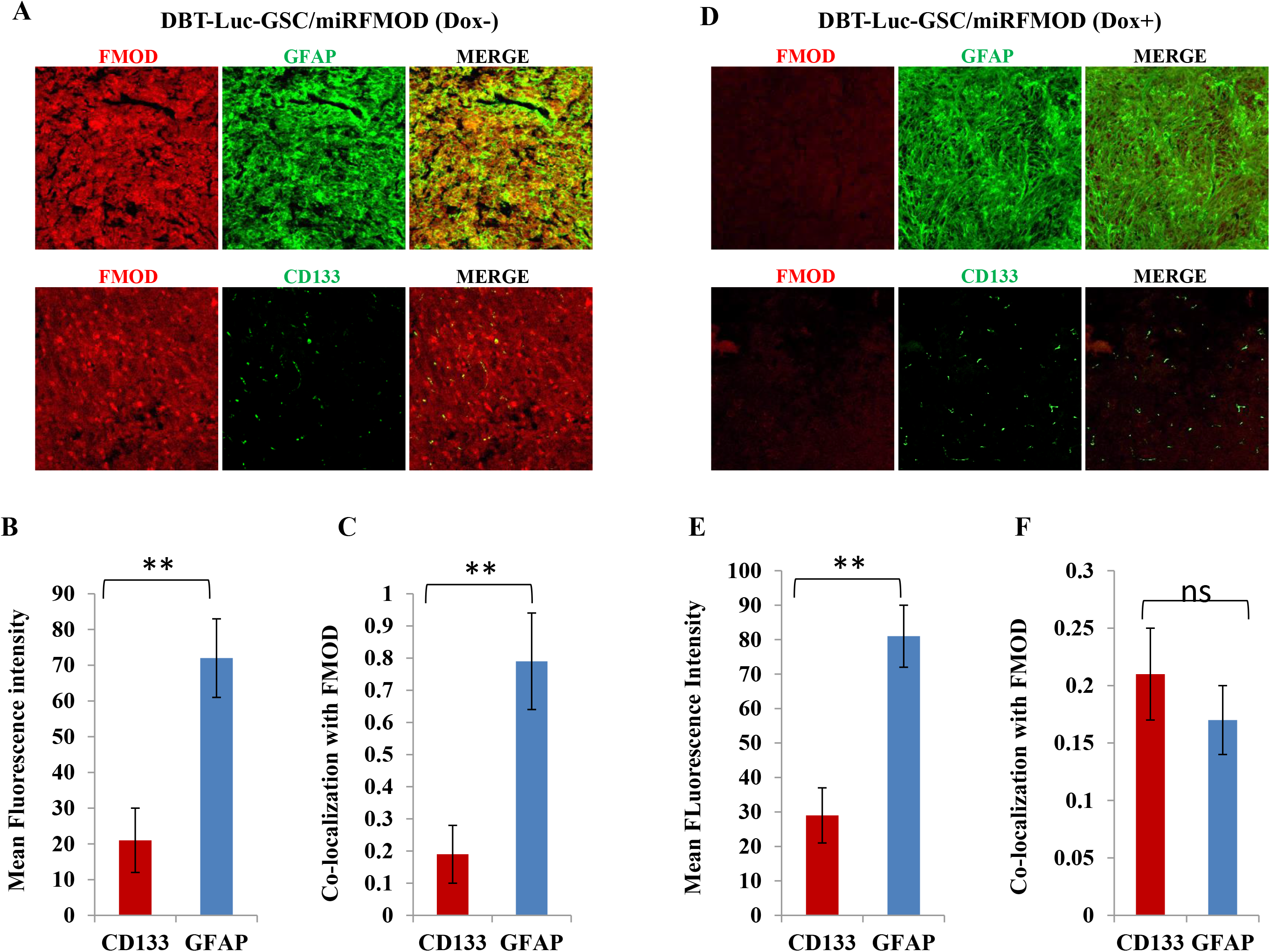
FMOD-silencing does not hamper the differentiation potential of tumor cells in murine glioma models. **A.** Immunohistochemical analysis of brain-derived sections obtained from mice injected with DBT-Luc-GSC/miRFMOD cell show FMOD, GFAP, and CD133 expression, and co-localization of FMOD with GFAP, and not with CD133 (left, before doxycycline injection). **B.** Bar diagram quantifying mean fluorescence intensity of CD133 and GFAP in DBT-Luc-GSC/miRFMOD cells without doxycycline injection. **C.** Bar diagram quantifying co-localization of FMOD with CD133 and GFAP, respectively, in DBT-Luc-GSC/miRFMOD cells before doxycycline injection. **D.** Immunohistochemical analysis of brain-derived sections obtained from mice injected with DBT-Luc-GSC/miRFMOD cell show low FMOD, and high GFAP, and CD133 expression, and loss of co-localization of FMOD with GFAP (right, with doxycycline injection). **E.** Bar diagram quantifying mean fluorescence intensity of CD133 and GFAP in DBT-Luc-GSC/miRFMOD cells after doxycycline injection. **F.** Bar diagram quantifying co-localization of FMOD with CD133 and GFAP, respectively, in DBT-Luc-GSC/miRFMOD cells after doxycycline injection. Magnification for panels **A** and **D** =20 X, Scale= 100 μm. p-values for panels **B, C, E**, and **F** are calculated by Student’s t-test, p-value are represented by *, **, *** indicate values <0.05, <0.01, and <0.001 respectively.

**Supplementary Figure 26:**
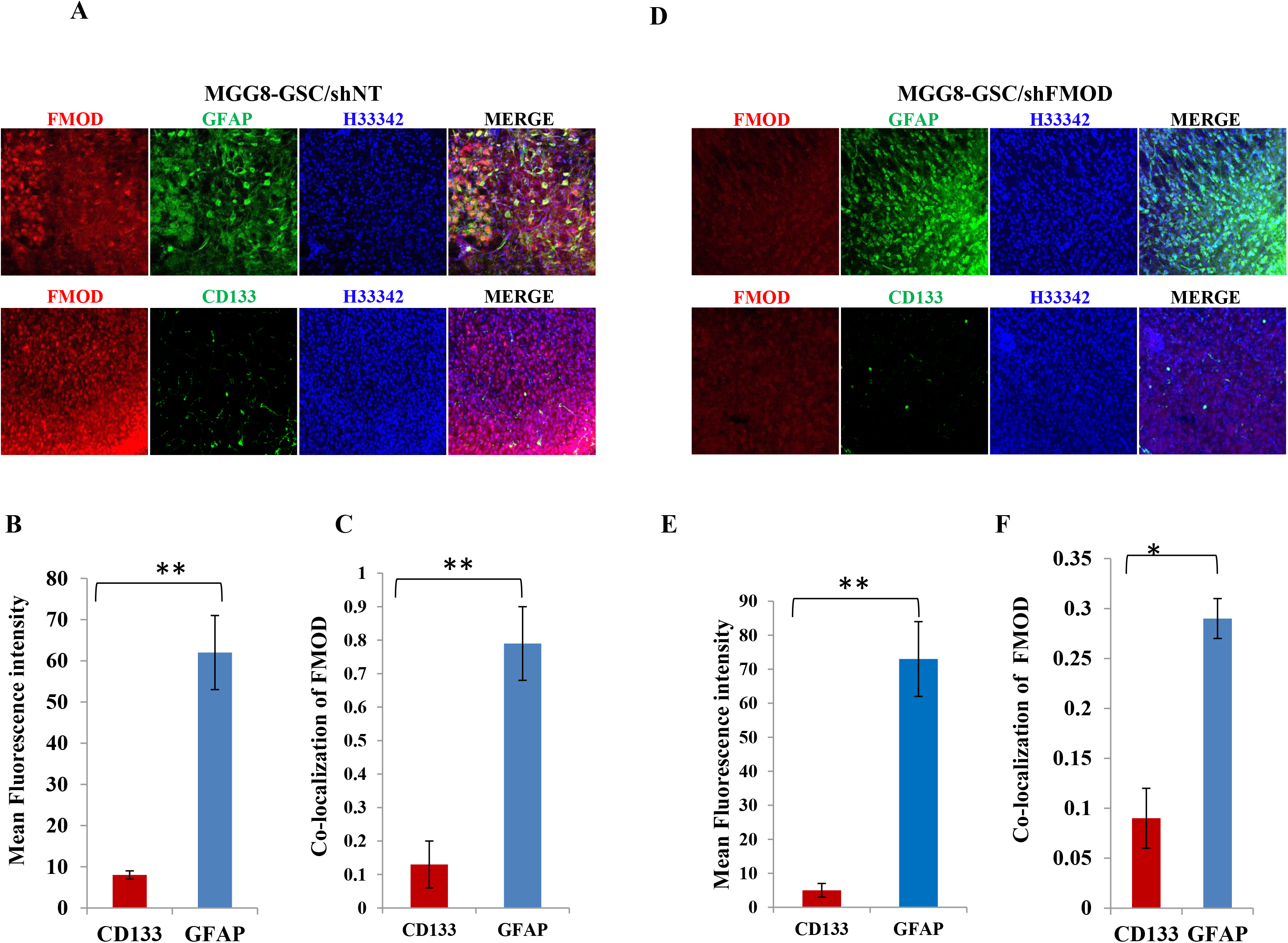
FMOD-silencing does not hamper the differentiation potential of tumor cells. **A.** Immunohistochemical analysis of brain-derived sections obtained from mice injected with MGG8-GSC/shNT cells show FMOD, GFAP, and CD133 expression, and co-localization of FMOD with GFAP, and not with CD133 (left). **B.** Bar diagram quantifying mean fluorescence intensity of CD133 and GFAP in MGG8-GSC/shNT cells. **C.** Bar diagram quantifying co-localization of FMOD with CD133 and GFAP, respectively, in MGG8-GSC/shNT cells. **D.** Immunohistochemical analysis of brain-derived sections obtained from mice injected with MGG8-GSC/shFMOD cells show low FMOD, and high GFAP, and CD133 expression, and loss of co-localization of FMOD with GFAP (right, after doxycycline injection). **E.** Bar diagram quantifying mean fluorescence intensity of CD133 and GFAP in MGG8-GSC/shFMOD cells. **F.** Bar diagram quantifying co-localization of FMOD with CD133 and GFAP, respectively, in MGG8-GSC/shFMOD cells. Magnification for panels **A** and **D** =20 X, Scale= 100 μm. p-values for panels **B, C, E**, and **F** are calculated by Student’s t-test, p-value are represented by *, **, *** indicate values <0.05, <0.01, and <0.001 respectively.

**Supplementary Figure 27:**
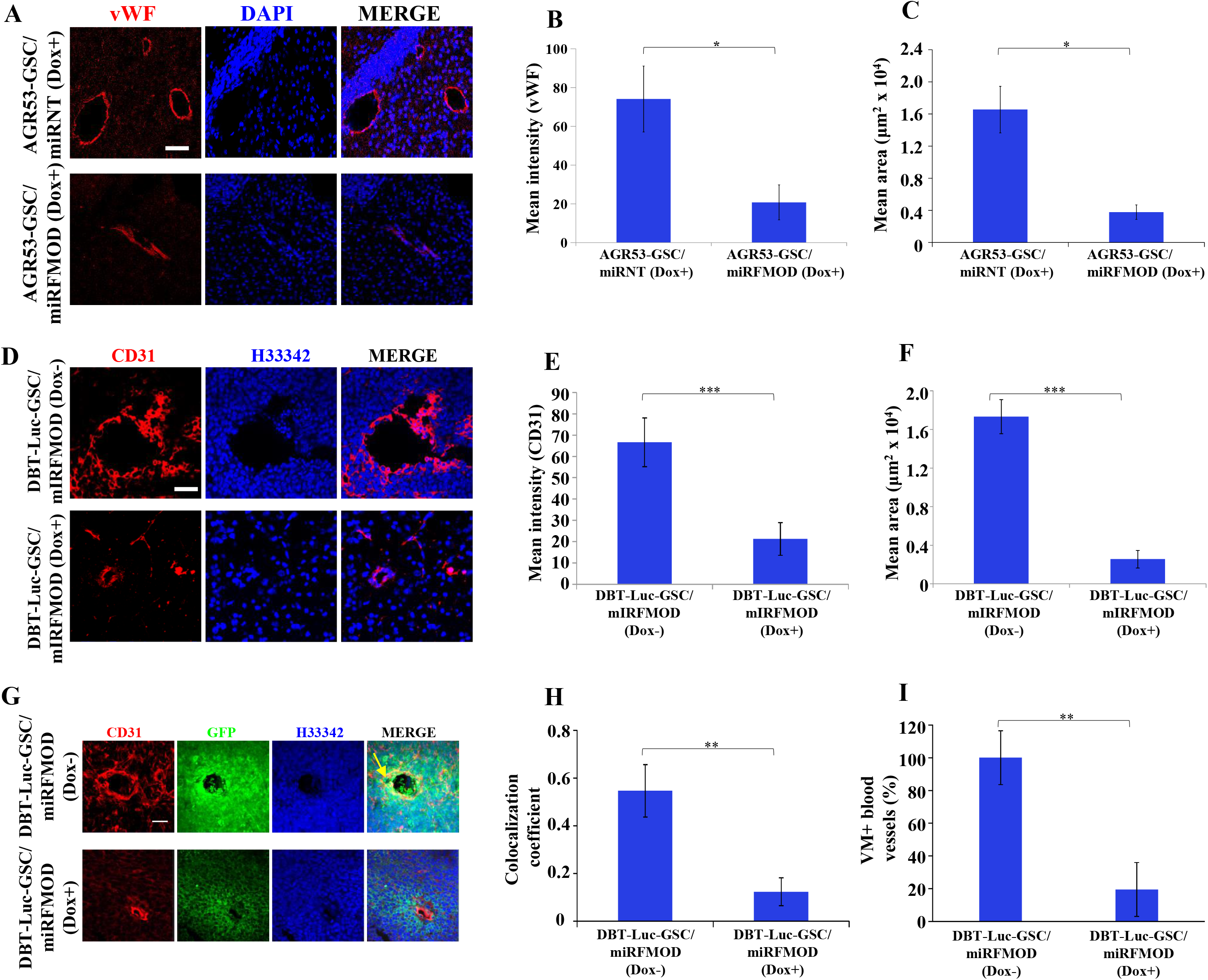
FMOD-silencing affects angiogenesis and vascular mimicry *in vivo.* **A.** Immunohistochemical analysis shows that the expression of von Willebrand Factor (vWF), a blood vessel marker, lining the blood vessels in both AGR53-GSC/miRNT and AGR53-GSC/miRFMOD groups, after doxycycline injection. Brain sections of two time points, day21 and day 55 were stained for the expression of vWF. At both time points, AGR53-GSC/miRFMOD shows lesser vWF staining than the AGR53-GSC/miRNT group. Magnification=20x, Scale= 100 μm. **B.** Bar diagram showing the mean intensity of vWF and **C.** The mean area of blood vessels in the AGR53-GSC/miRNT and AGR53-GSC/miRFMOD groups. **D.** Immunohistochemical analysis shows that the expression of CD31, lining the blood vessels in both DBT-Luc-GSC/miRFMOD(Dox-) and DBT-Luc-GSC/miRFMOD(Dox+) groups. **E.** Bar diagram showing the mean intensity of vWF and **F.** The mean area of blood vessels in the DBT-Luc-GSC/miRFMOD(Dox-) and DBT-Luc-GSC/miRFMOD(Dox+) groups. **G.** Immunohistochemical analysis showing overlap of CD31 (red) and GFP (green) expression in brains of mice injected with both DBT-Luc-GSC/miRFMOD cells, with or without doxycycline injection. Yellow allow indicates the region depicting the colocalization of both the two markers. **H.** Quantification of the colocalization coefficient in both DBT-Luc-GSC/miRFMOD(Dox-) and DBT-Luc-GSC/miRFMOD(Dox+) groups. **I.** Quantification of the VM+ blood vessels, indicating the measure of vascular mimicry (VM) in both DBT-Luc-GSC/miRFMOD(Dox-) and DBT-Luc-GSC/miRFMOD(Dox+) groups. p values calculated by unpaired t test with Welch’s correction are indicated. p value less than 0.05 is considered significant with *, **, *** representing p value less than 0.05, 0.01 and 0.001, respectively. ns stands for non-significant. Magnification = 20x, Scale=100 μm.

**Supplementary Figure 28:**
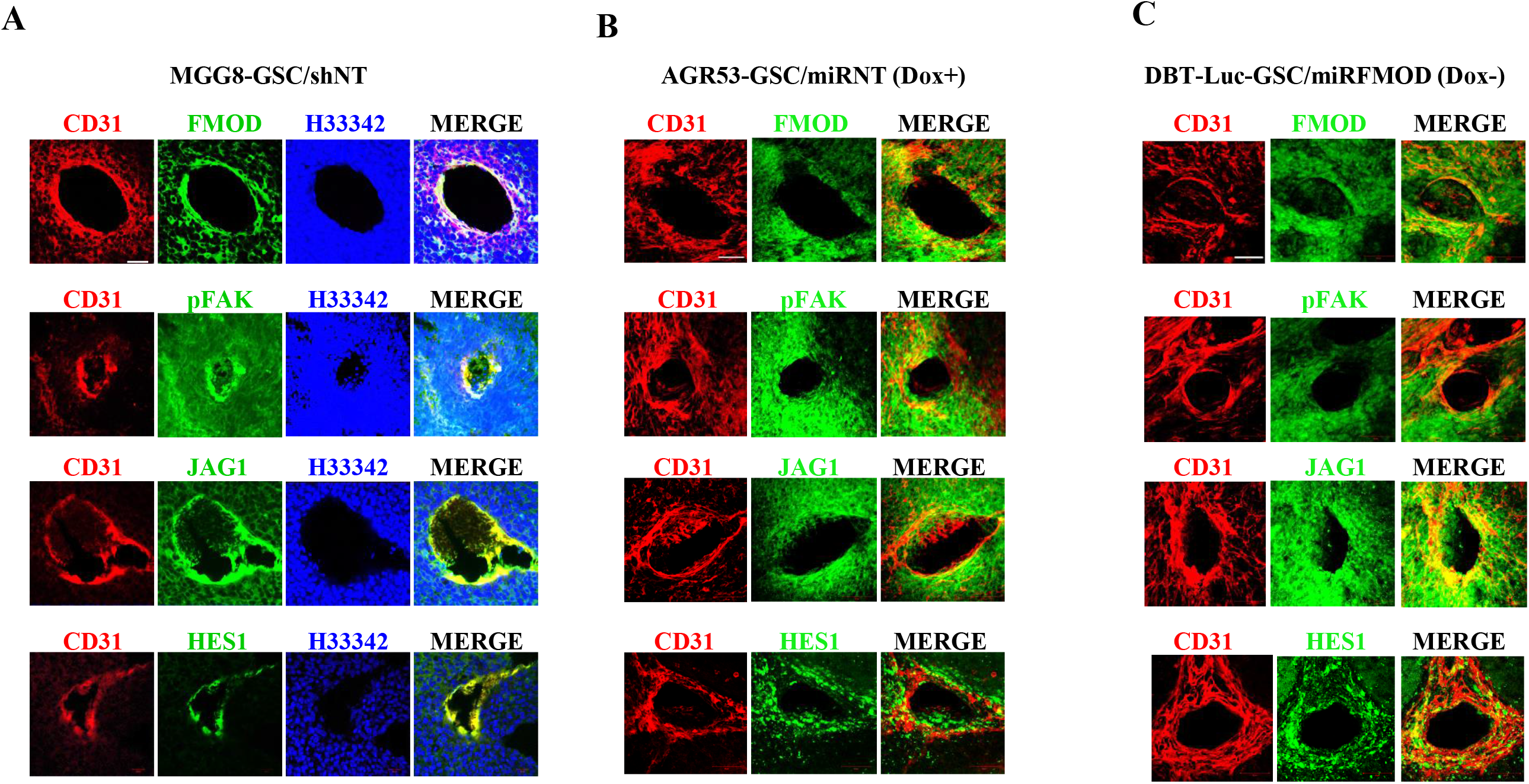
FMOD-dependent activation of the integrin-FAK-JAG1-HES1 signaling axis is maintained *in vivo*. **A.** Immunohistochemical analysis shows co-staining of CD31 with FMOD, pFAK, JAG1, and HES 1 in brain section derived from MGG8/shNT groups of animals. **B.** Immunohistochemical analysis shows co-staining of CD31 with FMOD, pFAK, JAG1, and HES 1 in brain section derived from AGR53-GSC/miRNT group of animals, after doxycycline injection. **C.** Immunohistochemical analysis shows co-staining of CD31 with FMOD, pFAK, JAG1, and HES 1 in brain section derived from DBT-Luc-GSC/miRFMOD group of animals, without doxycycline injection. Magnification = 20x, Scale=100 μm.

@@@@@@@@@@@@@@@@@@@@@@@@@@@@@@@@@@@@@@@@@@

